# Design of Tissue-Selective PROTACs Through Recruiting E3 Ligase Scaffolding Protein MAGEA11

**DOI:** 10.1101/2025.11.03.686228

**Authors:** Isabella E. Jacobsen, Rui Shi, Cole R. Scholtz, William C.K. Pomerantz, Gunda I. Georg

## Abstract

Proteolysis targeting chimeras (PROTACs) are an emerging therapeutic modality that induces protein degradation by recruiting E3 ligases. Most reported PROTACs recruit ubiquitously expressed E3 ligases, such as cereblon and the von Hippel-Lindau tumor suppressor. Of the additional 600+ E3 ligases, recruiting those with tissue-restricted expression is attractive for increasing the specificity of PROTACs. To this end, tissue-specific E3 ligases or E3 ligase-associated proteins that can be recruited for targeted protein degradation need to be identified. This work describes the first reported PROTAC that recruits the tissue-specific E3 ligase scaffolding protein MAGEA11. As an initial demonstration, a library of bromodomain and extra-terminal domain (BET)-targeting PROTACs that recruit MAGEA11 was synthesized. The library was screened in osteosarcoma U2OS cells, identifying lead compound 105B. 105B potently degrades BET proteins in U2OS osteosarcoma cell lines (BRD4 DC_50_ = 0.130 nM, D_max_ = 78%) and KYSE180 esophageal squamous cell carcinoma cell lines (DC_50_ = 40 nM, D_max_ = 70%), but shows no degradation in non-cancerous, MAGEA11-deficient HEK293T cells. Mechanistic studies confirmed 105B’s dependence on the ubiquitin-proteasome system and engagement of both MAGEA11 and BRD4. 105B decreased levels of BET-regulated gene products c-Myc, RUNX2, and KRT14; however, improvements are still necessary to affect selective cytotoxicity. This work reports the first example of a PROTAC recruiting a tissue-specific E3 ligase for cancer-restricted degradation of BET proteins and highlights the need for further development of MAGEA11-recruiting degraders.

## INTRODUCTION

Proteolysis-targeting chimeras (PROTACs) are an emerging class of therapeutics that hijack a cell’s ubiquitin-proteasome system to degrade proteins of interest. These heterobifunctional molecules are formed through linking a ligand for a protein of interest (POI) to a ligand capable of recruiting an E3 ligase or E3 ligase scaffolding protein. Upon binding and recruitment of both the POI and the E3 ligase (termed ternary complex formation), the E3 ligase transfers ubiquitin to the POI ultimately leading to proteasome-mediated degradation.^1^ The event-driven pharmacology of PROTACs has distinct advantages over traditional occupancy-driven therapeutics; namely, their catalytic function means they are typically effective at low concentrations, their targets have reduced potential for resistance mutations, and they can be used to target traditionally undruggable proteins.^2^ Substantial progress has been made in the development of PROTACs, with multiple PROTACs now in clinical trials for cancer and central nervous system disorders.^3^

One key advantage of PROTACs is that they have the potential to impart additional selectivity for their targets. As one approach, the length and composition of the linker have been modified to increase a PROTAC’s selectivity for a particular protein in a family of structurally similar proteins (e.g. BET family proteins) or a certain protein isoform (e.g. p38α/*δ*) by inducing favorable POI/E3 ligase interactions.^4-7^ Photo-controlled degraders, or PHOTACs, have been developed to direct degradation to a certain tissue.^8-10^ To enhance degradation in cancer, PROTACs have also been conjugated to cell-surface-targeting ligands or antibodies for cancer biomarkers, such as DHFR and HER2, or designed to recruit E3 ligases overexpressed in cancer, such as IAP, for improving the therapeutic window.^11-13^

A largely unexplored area in developing selective PROTACs is the recruitment of tissue-specific E3 ligases. The majority of reported PROTACs recruit one of two ubiquitously expressed E3 ligases: cereblon and Von Hippel-Lidau tumor suppressor (VHL).^14^ As a tissue-selective alternative, there has been growing interest in the development of a PROTAC that could recruit an E3 ligase exclusively expressed in cancer cells.^14^ Targeting the anti-cancer B-cell lymphoma extra-large protein (BCL-xL) with a VHL-based PROTAC demonstrates the feasibility of tissue-specific E3 ligase recruitment.^13^ The reported PROTAC degrades BCL-xL in cancer cells, where VHL is highly expressed, with limited degradation of BCL-xL in platelet cells, reducing risks of thrombocytopenia. Given this precedence, cancer-specific E3 ligases are desirable for tissue-selective PROTAC degradation.

The Melanoma Antigen Gene (MAGE) family of cancer-testes antigens, some of which scaffold E3 ligases to direct ubiquitination, are cancer-specific proteins that could be recruited for targeted protein degradation.^15^ MAGE family member A11 (MAGEA11) binds E3 ligase HUWE1 to direct ubiquitination of MAGEA11’s substrate protein, PCF11.^16^ In healthy cells, MAGEA11 is expressed exclusively in sperm and placental tissues and is associated with germ cell maturation.^17^ MAGEA11 is expressed aberrantly in a variety of cancers, including osteosarcoma, castration-resistant prostate carcinoma, esophageal squamous cell carcinoma (ESCC), and non-small cell lung carcinoma.^18-21^ Therefore, MAGEA11 recruitment to POIs is hypothesized to result in tissue-specific degradation in cancer, sperm, and placental tissues.

To probe MAGEA11’s ability to be recruited for PROTAC development, an initial proof-of-concept PROTAC was developed to degrade bromodomain and extra-terminal domain (BET) family proteins (including BRD2, 3, 4, and T). Bromodomain-containing protein 4 (BRD4) was chosen as the initial target due to its known role in tumorigenesis and the dose-limiting toxicities associated with its inhibition or degradation.^22^ BRD4 is an epigenetic reader protein that binds to acetylated lysine residues on histones to regulate gene transcription. Its binding leads to the recruitment of transcription factors, release of paused RNA polymerase II, and initiation of transcription elongation.^23^ All BET proteins contain two acetyl lysine binding pockets termed bromodomain 1 (BD1) and bromodomain 2 (BD2).^24^ Binding of BRD4 to acetylated lysine residues of histone H4 drives the expression of oncogenic transcription factor c-Myc in osteosarcoma and ESCC.^25, 26^ Inhibition and degradation of BRD4 in these cancer models cause tumor growth inhibition.^24, 26-29^ However, no BET-targeted therapies have progressed beyond clinical trials due to dose-limiting toxicities of BRD4 inhibition in healthy cells which cause side effects including thrombocytopenia and gastrointestinal events.^22^ Therefore, for a BRD4-targeting therapy to be clinically viable, it would need to have minimal effects in healthy cell lines.

Here, we report a tissue-specific PROTAC that recruits the cancer-(and placental/sperm-) specific E3 ligase scaffolding protein MAGEA11. The designed PROTAC links a MAGEA11 ligand to a BET ligand, causing HUWE1-mediated ubiquitination and degradation of BET proteins exclusively in cancer cells (**Figure 1**). We identify a hit PROTAC, 105B, which degrades BRD4 and other BET proteins in osteosarcoma and ESCC cell lines at low nanomolar concentrations and has no effect in non-cancerous cell lines. This proof-of-concept study is the first example of a PROTAC that recruits a cancer-specific E3 ligase, demonstrating the possibility of imparting PROTAC selectivity via this approach.

**Figure 1.**
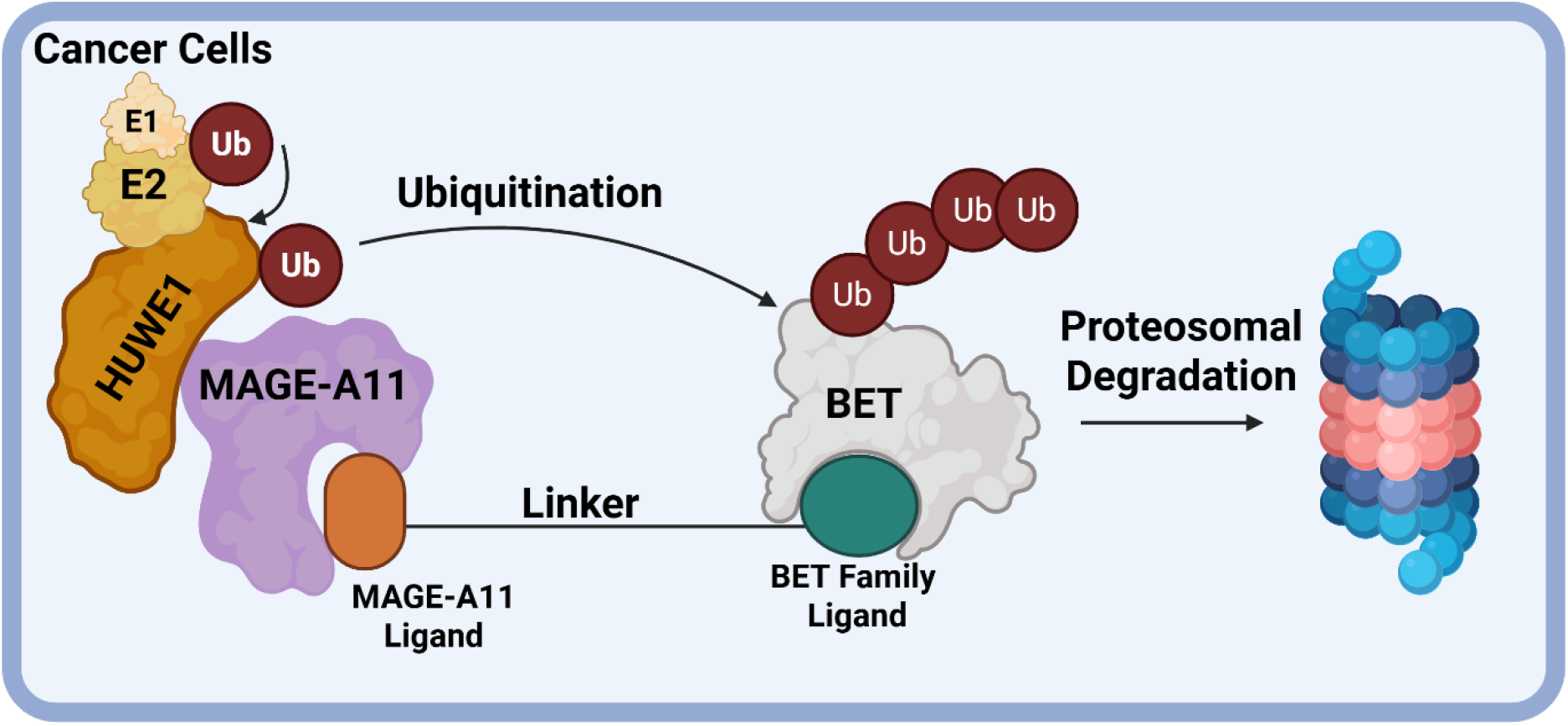
Design of a BET targeting PROTAC recruiting MAGEA11.

## RESULTS

### Design of a MAGEA11 Recruiting PROTAC and Identification of Lead 105B

MAGEA11-recruiting PROTACs were designed based on known MAGEA11 and BET family ligands. A ligand for MAGEA11 was reported by Yang et. al in 2020 (SJ1008066; **Figure 2a**).^27^ This ligand is hypothesized to bind at the PCF11 binding site of MAGEA11 with an IC_50_ of 130 nM, but no crystal structure of the ligand bound to MAGEA11 has been published. The structure-activity relationship of SJ1008066 demonstrated that the addition of large hydrophobic groups at the 7-position of the quinoline had minimal effect on its binding. These results indicate the 7-position is a permissive site for linker attachment. The BET ligand (+)-JQ1 was chosen due to its well-documented use in PROTACs and tolerance of linker addition at the tert-butyl carboxyl group (e.g., VHL-recruiting PROTAC MZ-1, **Figure 2b**).^4^ As no PROTAC recruiting MAGEA11 has been reported, a library of 14 PROTACs was designed and synthesized with various linker lengths, compositions, and attachment moieties (**Figure 2c**). PEG and alkyl linkers between 3-28 atoms in length were attached to (+)-JQ1 via ester or amide bond formation and to SJ1008066 via triazole formation (**Supplementary Scheme 1**), providing a large sampling of linker lengths and compositions.

**Figure 2.**
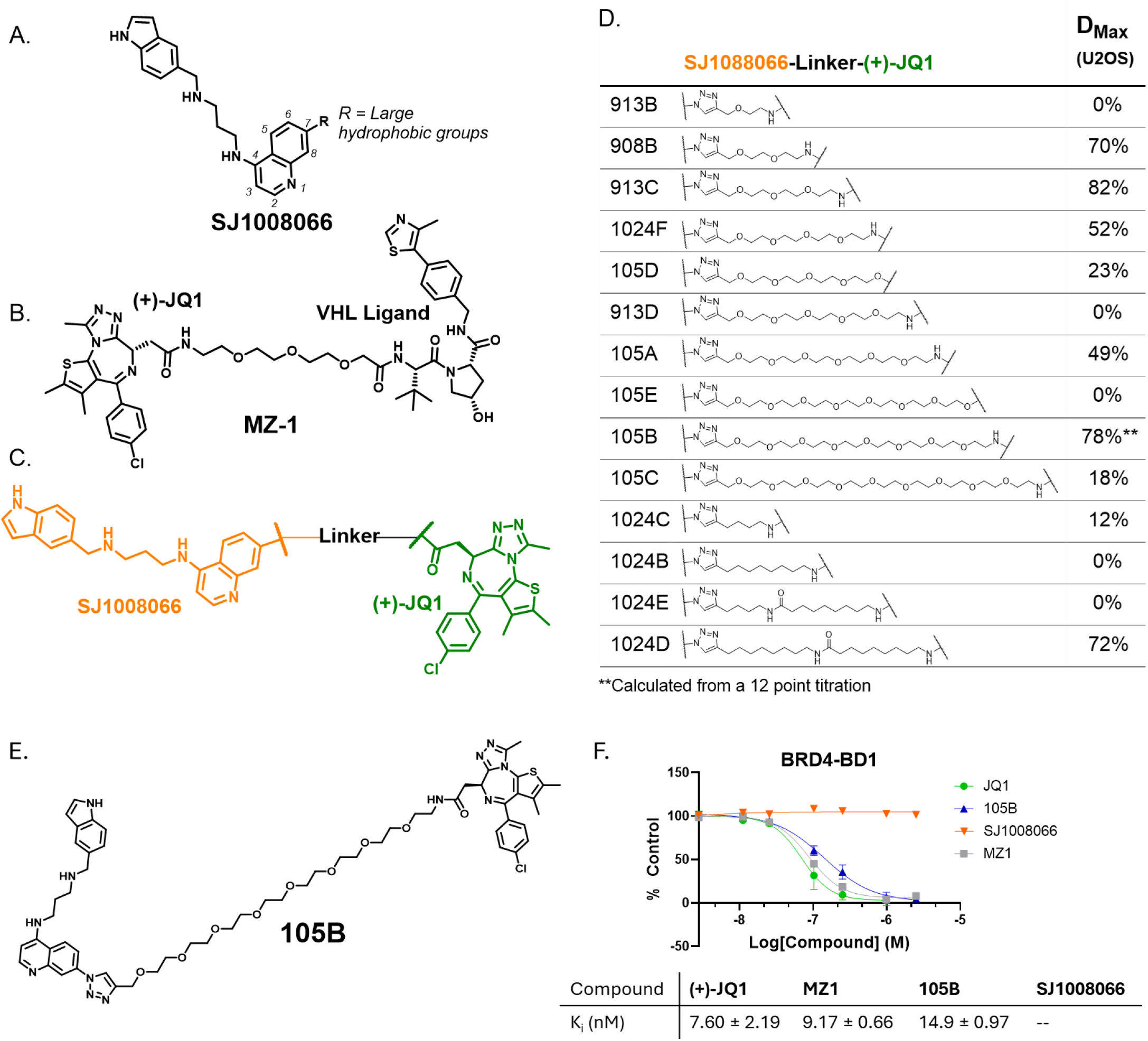
Design of a library of BET family-targeting PROTACs recruiting MAGEA11. A. MAGEA11 ligand SJ1008066 tolerates the addition of large hydrophobic groups to position 7 of the quinoline moiety.^27^ B. Known BET degrader MZ-1 targets BET proteins with (+)-JQ1. C. A library of degraders was designed with SJ1008066 linked to (+)-JQ1. D. Screening results of PROTAC library. E. Structure of lead compound 105B. F. 105B binds to BRD4 with a K_i_ of 14.9 ± 0.97 nM. K_i_ data calculated from a fluorescence polarization competitive inhibition assay with BRD4-BD1, n=3.

The PROTACs were initially screened in U2OS osteosarcoma cells due to their known sensitivity to BRD4 inhibition, degradation, and MAGEA11 expression, confirmed via western blot (**Supplementary Figure S1**).^25, 28, 29^ The PROTACs were screened in a 24 hour, 4-point titration from 10 - 0.01 µM concentrations. The BRD4 levels in U2OS cells after treatment were assessed using western blot analysis and compared to those of a known pan-BET degrader, MZ1, at 1 µM and a 0.1% DMSO vehicle control (**Figure 2d; Supplementary Figure S2**). From the screen, we selected 105B to further pursue due to its dose-dependent degradation of BRD4 (**Supplementary Figure S2**). Though 913C initially showed more degradation of BRD4, follow-up experiments did not show a dose-dependent effect. 105B features an 8 PEG-based linker with triazole and amide connections to SJ1008066 and (+)-JQ1, respectively (**Figure 2e**). The target engagement and binding affinity of 105B with the first bromodomain of (BRD4-BD1) was quantified with a fluorescence polarization assay to verify that linker attachment did not perturb binding. The K_i_ of 105B binding to BRD4-BD1 is 14.9 ± 0.97 nM, approximately two-fold higher than (+)-JQ1 and MZ-1 (K_i_ of 7.60 ± 2.19 nM and 9.17 ± 0.66 nM; **Figure 2f**), demonstrating minimal perturbation.

BRD4 degradation with 105B was then characterized in full dose-response experiments in multiple cell lines. In U2OS cells at 24 hours, 105B degraded BRD4 with a DC_50_ of 130 ± 30 pM and a D_max_ of 78 ± 2% (**Figure 3a; Supplementary Figure S3**). 105B exhibited a hook effect above ∼41 nM. 105B was also tested in KYSE180 cells, an ESCC cell line that is known to undergo transcriptional regulation by BRD4 and BRDT and expresses MAGEA11.^30,29^ 105B degraded BRD4 with a DC_50_ of 40 ± 11 nM and a D_max_ of 70 ± 3 % in KYSE180 cells (**Figure 3b; Supplementary Figure S4**). No significant hook effect was observed in these cell lines at concentrations up to 10 µM. Given the high potency of 105B in two cancer cell lines, the tissue specificity of 105B was next probed. 105B was tested in HEK293T cells, which are non-cancerous and do not express MAGEA11. At 24 hours, 105B does not significantly degrade BRD4 in HEK293T cells at any concentration tested (**Figure 3c; Supplementary Figure S5**). Together these results show that 105B is highly potent in two cancer cell lines expressing MAGEA11 and demonstrates no BRD4 degradation in HEK293T cells.

**Figure 3.**
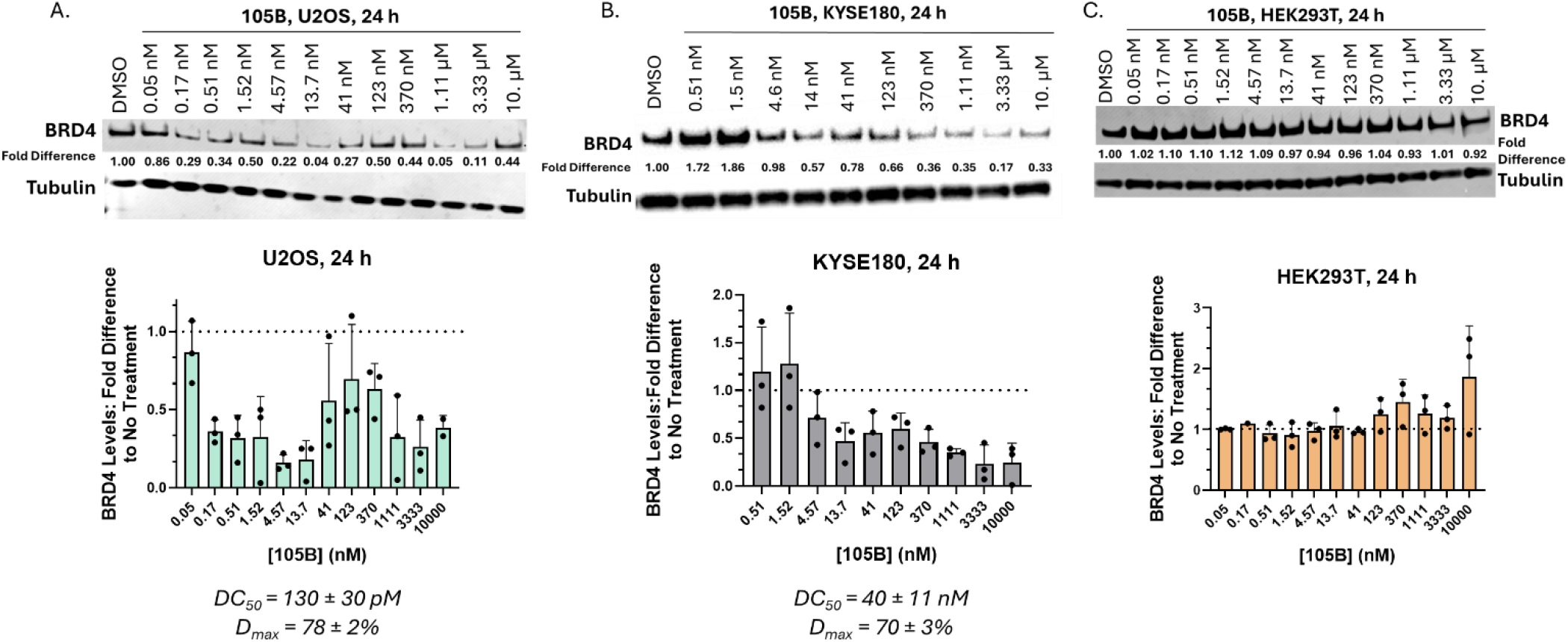
Characterization of 105B in U2OS, KYSE180, and HEK293T Cells. A. 105B degrades BRD4 in U2OS cells with a DC_50_ of 130 ± 30 pM and a D_max_ of 78 ± 2%. C. 105B degrades BRD4 in KYSE180 cells with a DC_50_ of 40 ± 11 nM and a D_max_ of 70 ± 3%. D. 105B does not degrade BRD4 in HEK293T cells at any concentration tested. All values calculated from triplicate western blot experiments; representative western blot shown. Graphical data represents mean ± standard deviation of triplicate data, n=3.

### 105B is Dependent on the Ubiquitin Proteasome System and MAGEA11 and BRD4 Engagement

With an established range of effective concentrations of 105B in U2OS lines, we next sought to test the time and proteasome dependence of 105B. U2OS cells were incubated with 105B at 10 nM and lysed every 30 min for the first two hours, every two hours until eight hours, and then at 24 hours (**Figure 4a; Supplementary figure S6**). 105B degraded BRD4 within four hours, with a variable amount of rebound at 24 hours, likely attributed to the resynthesis of BRD4 in cells. To test its dependence on the ubiquitin-proteasome system, 105B was dosed at both 25 nM and 100 nM in the presence of the proteasome inhibitor MG132 (10 µM) and the NEDDylation inhibitor MLN4924 (1 µM). Cells were incubated for only six hours given the known toxicity of MG132 at extended time points.^31^ Addition of both MG132 and MLN4924 restored BRD4 levels to those of the vehicle control (p<0.05), demonstrating that 105B is dependent on the function of the ubiquitin proteasome system (**Figure 4b; Supplementary Figure S7**).

**Figure 4.**
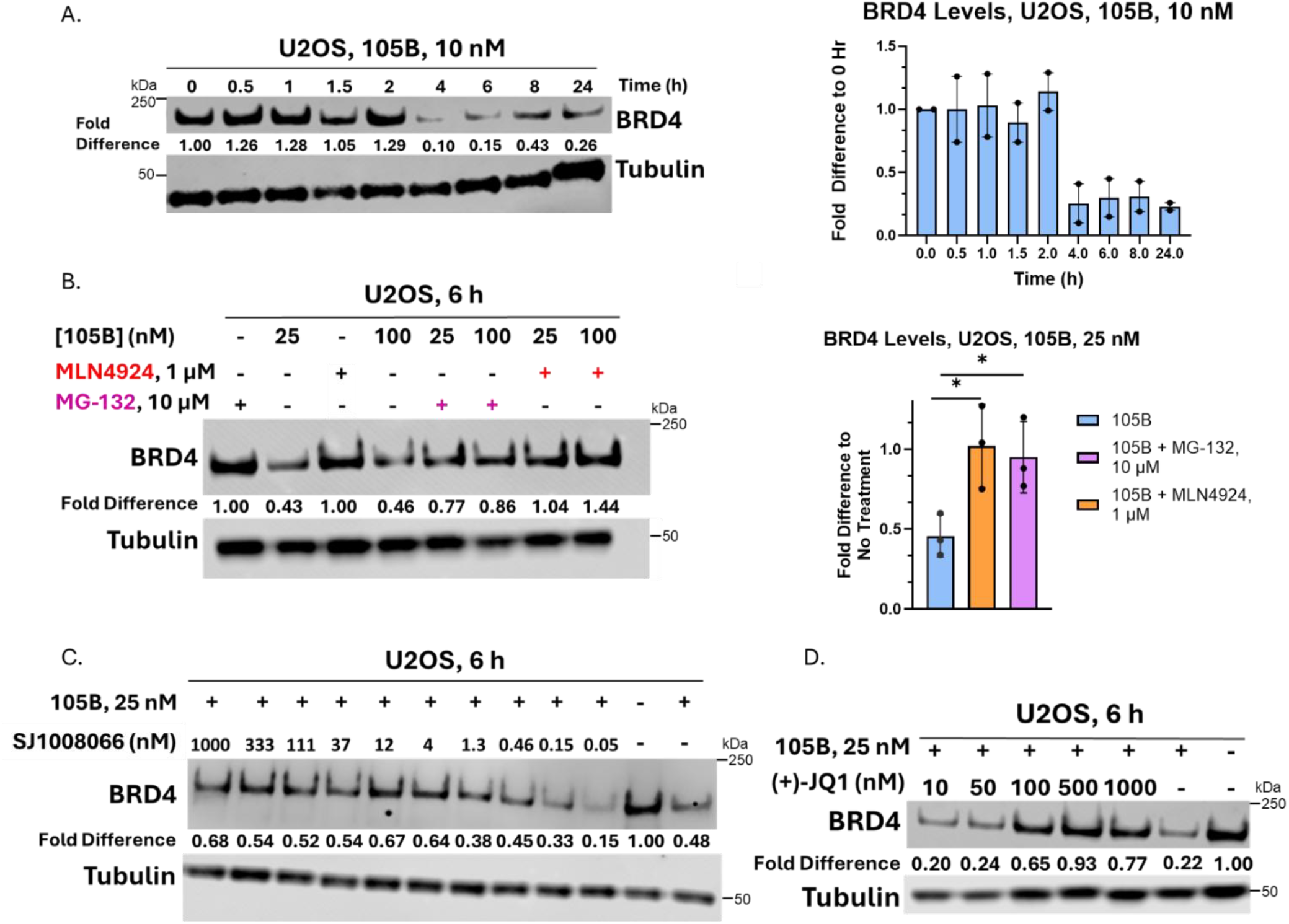
Time and Proteasome Dependence of 105B. A. Time course experiment with 105B at 10 nM shows BRD4 degradation by hour four. The graph represents biological duplicates; a representative western blot is shown. B. Incubation of cells with proteasome inhibitor MG132 and NEDDylation inhibitor MLN4924 inhibits the function of 105B. The experiment was repeated in biological triplicate; replicates are quantified in a bar graph, and a representative western blot is shown. Statistical significance was determined using an unpaired t-test in GraphPad Prism 10. * Represents p-value < 0.05. C. Addition of excess SJ1008066 partially reduces degradation of BRD4 with 105B at concentrations above 4 nM. D. Addition of (+)-JQ1 in excess of 105B reduces degradation of BRD4 at concentrations above 50 nM.

To probe its engagement of BRD4 and MAGEA11, 105B was dosed at 25 nM with increasing concentrations of (+)-JQ1 and SJ1008066 and incubated for six hours (synthesis of SJ1008066 **Supplementary Scheme S2**). SJ1008066 outcompeted the degrader at concentrations larger than 4 nM but did not fully restore BRD4 levels to those of the vehicle (**Figure 4c; Supplementary Figure S8a**). (+)-JQ1 outcompeted the degrader at concentrations above 50 nM and restored BRD4 levels to control levels at 500 nM (**Figure 4d; Supplementary Figure 8b**). These dependencies confirm that the observed decrease in BRD4 levels following treatment with 105B is caused by its recruitment of MAGEA11 and the subsequent ubiquitination and degradation of BRD4 by the 26S proteasome.

### 105B Degrades all BET Family Proteins and Decreases Oncogenic Protein Levels in U2OS and KYSE180 Cells

Since (+)-JQ1 binds to all BET family proteins, 105B could degrade all BET family proteins present in U2OS cells. To test this, 105B was incubated with U2OS cells for 24 hours in a 12-point, 3-fold titration from 10 µM – 0.05 nM concentrations, and BRD2 and BRD3 levels were probed with western blot. BRDT is not expressed in U2OS cells. 105B degraded both BRD2 and BRD3 at concentrations between 0.17 nM – 41 nM, consistent with the degradation trends of BRD4 (**Supplementary Figure 9a**). Unlike with BRD4, BRD2 degradation did not demonstrate a dramatic hook effect. Although some levels of BRD2 increased at 41 and 123 nM, degradation was restored from 370 nM – 10 µM. Given the precedent that BET proteins degrade at different rates, the degradation of BRD2 was assessed over time.^32^ BRD2 degradation happened at a similar rate to BRD4 degradation, with degradation starting between 4-6 hours. (**Supplementary Figure 9b**).

Given the potency and selectivity of 105B, its impact on oncogene expression in U2OS was initially probed. In U2OS cells, BRD4 regulates the transcription of oncogenes c-Myc and RUNX2.^25^ C-Myc levels were tested in U2OS cells after 24-hour incubation with 105B at four representative concentrations. 105B decreased c-Myc levels by over 80% at 100 nM and 1 µM concentrations (**Figure 5a; Supplementary Figure S10a**). However, (+)-JQ1 at 10 µM and MZ-1 at 1 µM had minimal effect on c-Myc levels. Some studies have reported downregulation of c-Myc transcript levels in HeLa and U2OS cells in the presence of MZ-1 and (+)-JQ1 at short time points.^4, 25^ We suspect that our observation of minimal perturbation of c-Myc levels at 24 hours is likely due to BRD4’s negative regulation of c-Myc via its phosphorylation capacity (**Figure 5a, Supplementary Figure S10b**).^33^ The transcription factor RUNX2 is regulated by BRD4 in U2OS cells to a greater extent than c-Myc.^25^ The effect of degradation with 105B on RUNX2 levels was therefore probed in U2OS cells. 105B degradation resulted in a reduction of RUNX2 levels of ∼50% for 105B at 1 uM, and ∼70% for (+)-JQ1 at 10 µM and ∼90% for MZ-1 at 1 µM (**Figure 5a; Supplementary figure S10a**,**b**). Given the consistent downregulation of RUNX2 with the control compounds, the downregulation observed with 105B is likely due to the degradation of BRD4.

**Figure 5.**
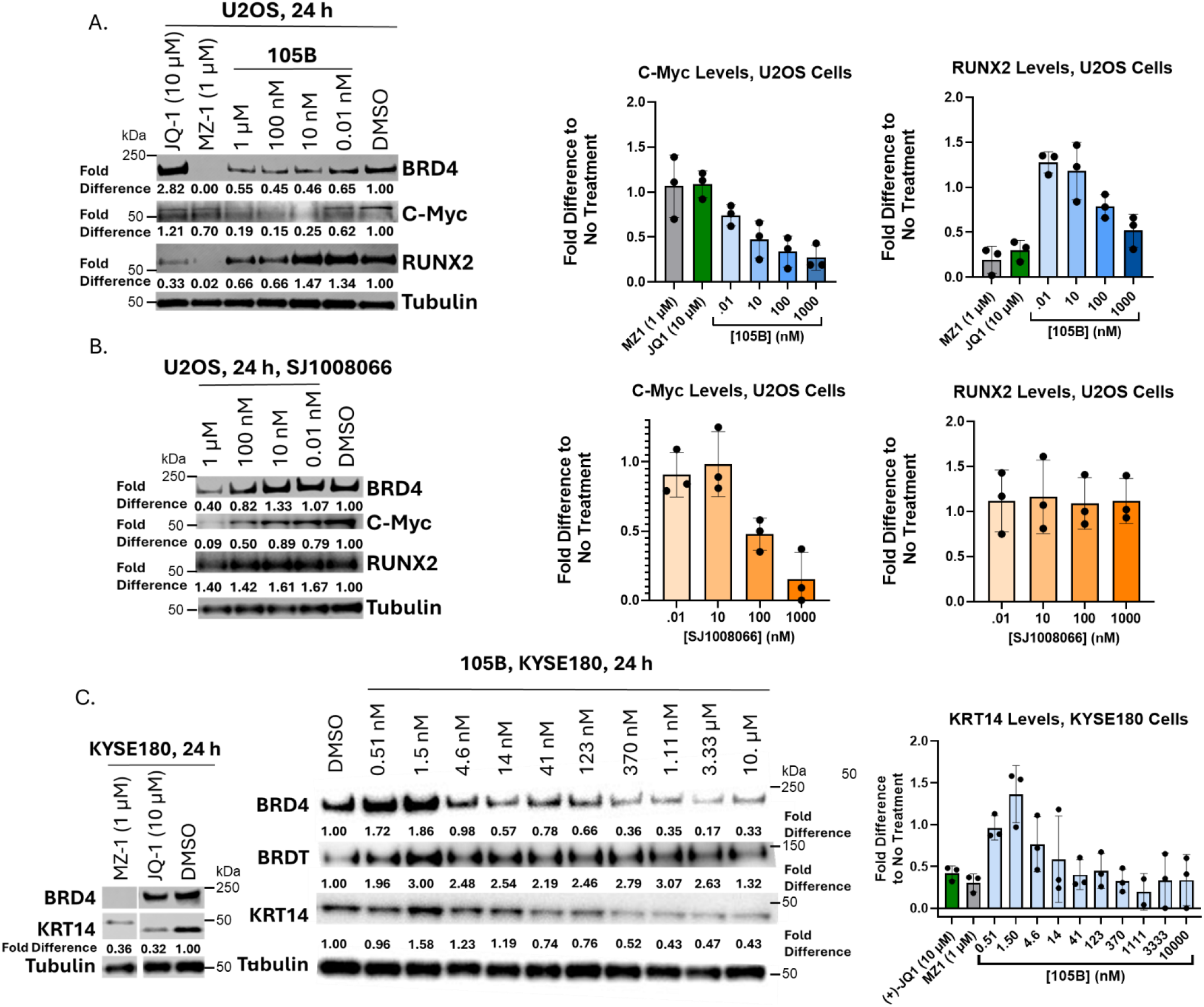
105B and SJ1008066 Downregulate Downstream Genes Regulated by BRD4 in U2OS Cells. A. 105B downregulates c-Myc and RUNX2 in U2OS cells. Graphical representation of triplicate data ± standard deviation and representative western blot shown. B. MAGEA11 inhibitor SJ1008066 downregulates c-Myc and shows downregulation of BRD4 but does not affect RUNX2 levels in U2OS cells. Graphical representation of triplicate data ± standard deviation and representative western blot shown. C. MZ-1 and (+)-JQ-1 reduce KRT14 levels in KYSE180 cells. 105B does not degrade BRDT but does decrease KRT14 levels in KYSE180 cells. Graphical representation of triplicate data ± standard deviation and representative western blot shown.

Since 105B significantly decreased c-Myc levels, while the two BRD4-targeting controls did not, it was hypothesized that the MAGEA11 engagement of 105B could have regulatory effects on c-Myc. To test this, U2OS cells were incubated with SJ1008066 for 24 hours, and the levels of c-Myc, BRD4, and RUNX2 were probed. As hypothesized, SJ1008066 significantly downregulated c-Myc at concentrations as low as 100 nM, though SJ1008066 did not have significant effects on RUNX2 levels (**Figure 5b; Supplementary Figure S11**). SJ1008066 also decreased BRD4 levels at 1 µM. Though the mechanism is unclear regarding BRD4 downregulation, the observation could explain why a decrease in BRD4 was seen at concentrations beyond the hook effect of 105B (**Figure 3a**) and why SJ1008066 competition experiments with 105B did not fully restore BRD4 levels at high concentrations (**Figure 4c**). The observed decrease of c-Myc by SJ1008066 could be a consequence of BRD4 downregulation by MAGEA11 inhibition or a direct consequence of MAGEA11 inhibition. No interaction between MAGEA11 and c-Myc or BRD4 has been reported at this time; however, these results point to a dual mechanism for 105B’s downregulation of BRD4 and c-Myc levels.

In the ESCC cell line KYSE180, BRDT and BRD4 are colocalized at super-enhancer regions that regulate expression of migration-associated protein KRT14.^30^ We found that both MZ-1 at 1 µM and (+)-JQ1 at 10 µM decreased KRT14 levels by about 70% after 24-hour incubation in KYSE180 cells (**Figure 5c; Supplementary Figure S12a**). 105B did not degrade BRDT at any concentrations tested (**Figure 5c; Supplementary Figure S12b**). Nonetheless, 105B did decrease KRT14 levels in a dose-dependent manner. KRT14 depletion peaked at 50-60% at concentrations between 370 nM – 10 µM (**Figure 5c**), suggesting that KRT14 levels can be modulated with 105B and does not require BRDT degradation.

The effects of 105B on cell viability were investigated with CellTiter-Glo 2.0 (Promega). Despite significant c-Myc downregulation, 105B had no significant effect on U2OS viability at any concentration after 72 hours of incubation, though positive controls MZ-1, (+)-JQ1, and SJ1008066 were effective at concentrations similar to those reported in the literature (**Supplementary Figure S13a**).^25, 27^ In KYSE180 cells, a 30% reduction in viability was observed at 10 µM 105B (**Supplementary Figure S13b**). However, a similar reduction of viability was seen in HEK293T cells, suggesting that the observed toxicity is not on-target (**Supplementary Figure S13c**). We hypothesize that this lack of toxicity in cancer cells is because 105B does not fully degrade any of the BET proteins in either U2OS or KYSE180 cells, allowing the cells to survive on the remaining BET proteins expressed.

## DISCUSSION

We have identified and characterized the first example of a PROTAC harnessing the E3 ligase scaffolding protein MAGEA11. A VHL-recruiting PROTAC targeting MAGE family member MAGE-A3 was recently reported, but no PROTAC recruiting MAGE proteins to degrade POIs has been reported.^34^ The PROTAC 105B has a sub-nanomolar DC_50_ in U2OS cells and a double-digit nanomolar DC_50_ in KYSE180 cells, but exhibits no BRD4 degradation in HEK293T cells. 105B has a DC_50_ comparable to the most potent reported BRD4 degraders, such as QCA570 (degradation as low as 10 pM) and ARV-825 (degradation as low as 30 pM).^35, 36^ However, 105B does not fully degrade BRD4 at any concentration tested and has a limited window of effective concentrations before exhibiting hook effects.

Our mechanistic studies confirmed that degradation with 105B is reliant on the ubiquitin-proteasome system and occurs between 4-24 hours. The speed and duration of BRD4 degradation with PROTACs can vary according to the E3 ligase recruited. For example, in HEK293T cells the BRD4 degrader dBET6 (cereblon-recruiting) reaches its peak D_Max_ around five hours and rebounds around 10 hours; whereas, MZ-1 (VHL-recruiting) peaks around one hour and maintains degradation through 24 hours.^32^ Degradation with 105B therefore follows a similar trend to dBET6 with degradation between 4-8 hours and rebound starting between 8-24 hours.

In both U2OS cells and KYSE180 cells, 105B decreased levels of proteins regulated by BRD4 transcription. We also discovered that MAGEA11 inhibition with SJ1008066 decreased BRD4 and c-Myc levels in U2OS cells. No direct connection has yet been made between MAGEA11 and BRD4/c-Myc. The observed effect could be due to the interaction between MAGEA11 and transcription factor E2F1, which upregulates c-Myc. MAGEA11 complexes with and stabilizes E2F1, so inhibition of this interaction could decrease c-Myc transcription.^37, 38^ This relationship between MAGEA11 and c-Myc could be further explored in future studies as a mechanism for targeting c-Myc in cancer.

Though 105B decreased oncogenic protein levels in U2OS and KYSE180 cells, 105B did not significantly decrease the cell viability of the cancer cell lines. We suspect that this observation is due to 105B not fully degrading any of the BET proteins at any concentration. Improvements to this compound need to be made to reach full BET degradation, such as incorporation of more rigid linkers for improved ternary complex formation.^39^

## CONCLUSION

This report details the synthesis and evaluation of the first PROTACs recruiting the cancer-specific E3 ligase scaffolding protein MAGEA11. A library of 14 PROTACs was synthesized and tested, and lead molecule 105B was identified. 105B degrades BET proteins BRD4, 3, and 2 at sub-nanomolar concentrations in U2OS cells and nanomolar concentrations in KYSE180 cells, but 105B does not degrade BRD4 proteins in MAGEA11-negative HEK293T cells. Mechanistic studies confirmed the time and ubiquitin-proteasome dependence of 105B. 105B decreased levels of oncogenic proteins c-Myc, RUNX2, and KRT14, but had no effect on cell viability. This first demonstration of a MAGEA11-recruiting PROTAC expands the toolbox of E3 ligase-associated proteins that can be recruited for targeted protein degradation and sets the stage for improvements upon this design for increased efficacy. PROTACs targeting different disease-associated proteins using the MAGEA11 ligand will be the subject of future work.

## Supporting information

Supplemental Figures and Characterization Data

## LIST OF ABBREVIATIONS

BCL-xL: B-cell lymphoma extra-large
BET: Bromodomain and extra-terminal domain
BD1, BD2: Bromodomain 1, bromodomain 2
BRD2, BRD3, BRD4, BRDT: Bromodomain-containing proteins 2, 3, 4, and testis-specific (T)
c-Myc: Myc proto-oncogene protein
DC_50_: Half-maximal degradation concentration
DHFR: Dihydrofolate reductase
D_max_: Maximum degradation achieved
DMSO: Dimethyl sulfoxide
E2F1: E2F transcription factor 1
ESCC: Esophageal squamous cell carcinoma
HER2: Human epidermal growth factor receptor 2
HUWE1: HECT, UBA, and WWE domain-containing E3 ubiquitin-protein ligase 1
IAP: Inhibitor of apoptosis protein
K_i_: Inhibition constant
KRT14: Keratin 14
MAGE: Melanoma antigen gene family
MAGEA11: Melanoma antigen family member A11
PEG: Polyethylene glycol
PHOTAC: Photo-controllable proteolysis-targeting chimera
PCF11: Cleavage and polyadenylation factor 11
POI: Protein of interest
PROTAC: Proteolysis-targeting chimera
RUNX2: Runt-related transcription factor 2
VHL: von Hippel–Lindau tumor suppressor protein

## ACKNOLWLEDGEMENTS

This research was supported by the NIH MIRA award R35 GM140837-01 (W.C.K.P.). I.E.J. was supported by an NIH T32 Chemical Biology Interface training grant, 5T32GM132029-06. C.R.S was supported by an NIH T32 Biotechnology training grant, T32GM008347. We thank the Steven A. Johnsen lab for providing the KYSE180 cell lines. BioRender.com was used in figure creation (graphical abstract and Figure 1).

## For Table of Contents Only

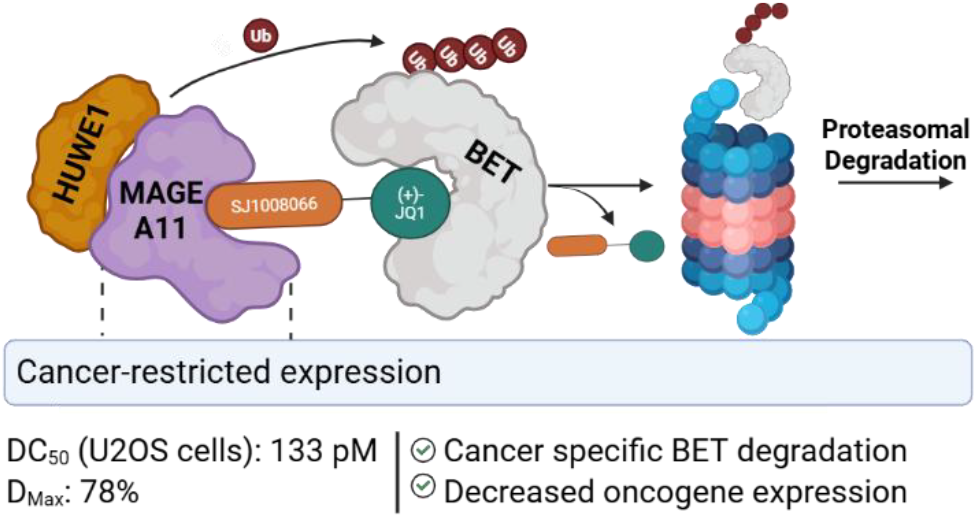

Synopsis: PROTAC degraders recruiting the cancer-specific E3 ligase scaffolding protein MAGEA11 to target BET family proteins were synthesized. Compound105B causes cancer-specific BET degradation.

