## Supplemental Figures and Characterization Data for "Design of Tissue-Selective PROTACs Through Recruiting E3 Ligase Scaffolding Protein MAGEA11"

#### Biological Procedures

##### Materials and Equipment

(+)-JQ1, MZ1, MG132, and MLN4924 were purchased as lyophilized powders from MedChem Express.

##### Cell Culture

All cell lines were cultured in a humidified environment at 37 °C with 5% $CO_2$. U2OS cells (ATCC HTB-96) were cultured in McCoy's 5A Media (ATCC 30-2007) with 5% 100 µg/mL penicillin-streptomycin solution and 10% fetal bovine serum. KYSE180 cells were cultured in RPMI 1640 media (ATCC 30-2001) with 5% 100 µg/mL penicillin-streptomycin solution and 10% fetal bovine serum. HEK293T cells (ATCC CRL-2316) were grown in DMEM (ATCC 30-2002) with 5% 100 µg/mL penicillin-streptomycin solution and 10% fetal bovine serum. For treatment, cells were plated in 6-well plates (Corning Costar 3516) at a density of 200,000 cells per well and allowed to adhere overnight. The compounds to be tested were dissolved in DMSO to a concentration of 10 mM and further diluted in media to achieve a final concentration of 1% DMSO. A 10x solution of the compounds was added to the cell plates at the desired concentration, and the cells were incubated for the desired time. The media was then removed, and the cells were washed with 1 mL of cold PBS. The cells were lysed with RIPA buffer (Sigma Aldrich) supplemented with cOmplete mini protease inhibitor cocktail (Roche, 1 tablet in 10 mL RIPA).

##### Western Blot

The soluble and insoluble layers of cellular lysates were separated via centrifugation (15000 rpm, 4 °C, 15 min). The soluble fraction was then removed, and the total protein concentration of each lysate was determined via a BCA assay kit (Thermo Fischer). The total protein levels of each lysate were then normalized by dilution with RIPA buffer, and 4x NuPAGE LDS sample buffer (Invitrogen) was added to each sample. The samples were boiled for 5 minutes, and the proteins were separated on a NuPAGE 3-8% tris-acetate gel (Invitrogen). The proteins were transferred to a PVDF membrane (BioRad) using a BioRad semi-dry Turbo Blot transfer system for 10 minutes, blocked with 5% milk in tris-buffered saline with 0.1% Tween 20 (TBST) for 1 h, and incubated with the appropriate antibody at 4 °C for 16 h. The membranes were then washed with TBST (10 mL, 5 x 3 min), incubated with the appropriate secondary antibody for 1 h, and washed with TBST (10 mL, 5 x 3 min). The proteins were visualized with SuperSignal West substrate (either Pico or Femto -Thermo Fischer) and a Licor Omega or BioRad ChemiTouch imaging system. Protein bands were quantified with ImageJ software and normalized to the appropriate loading control. Select blots were stripped with Restore™ PLUS Western Blot Stripping Buffer (Thermo Scientific) according to the manufacturer's protocol.

| Protein Target | Species | Manufacturer (product #) | Dilution | Substrate |
|---|---|---|---|---|
| BRD4 | Rabbit | Cell Signaling Technology (E2A7X) | 1:1000 | Pico |
| BRD2 | Rabbit | Cell Signaling (D89B4) | 1:1000 | Femto |
| BRD3 | Rabbit | Bethyl (BLR069G) | 1:1000 | Femto |
| BRDT | Rabbit | Cell Signaling (E6T6E) | 1:1000 | Pico |
| c-Myc | Rabbit | Cell Signaling (D84C12) | 1:1000 | Femto |
| RUNX2 | Rabbit | Cell Signaling (D1H7) | 1:1000 | Femto |
| KRT14 | Rabbit | Cell Signaling (46251S) | 1:1000 | Pico |
| MAGEA11 | Rabbit | Cell Signaling (E2F1K) | 1:1000 | Femto |
| Tubulin | Rabbit | Cell Signaling (2148) | 1:2000 | Pico |
| Actin | Mouse | Millipore Sigma (A3853) | 1:2000 | Pico |
| Vinculin | Mouse | Invitrogen (14-9777-82) | 1:1000 | Femto |
| Mouse- HRP Conjugate | Goat | Invitrogen (31430) | 1:2000 | NA |
| Rabbit- HRP Conjugate | Goat | Invitrogen (31460) | 1:2000 | NA |

**Fluorescence Polarization**

A his-tagged BRD4-BD1 was recombinantly expressed (residues 44-168), and a fluorescence polarization assay was performed as previously described.[1]

**Cell Viability**

Cells were seeded in a 384-well white opaque plate (Corning 3570) at a density of 500 cells/well in 25 µl total volume and allowed to adhere overnight. To each well was added a 10x stock of the desired compound at the desired concentration and titration, and the cells were incubated with the compound for 72 h. Viability was measured with CellTiter-Glo® 2.0 Cell Viability Assay (Promega) as follows: The cell plates and CellTiter-Glo® 2.0 reagent were allowed to equilibrate to room temperature for 30 minutes. CellTiter-Glo® 2.0 reagent (25 µl) was added to each well, and the plate was shaken on an orbital shaker for 2 minutes, then allowed to equilibrate undisturbed at room temperature for 10 minutes. The luminescence of each well was measured for 1 second with a Tecan Spark. The raw luminescence values were normalized to a vehicle control and plotted with GraphPad Prism 10 to calculate a $GI_{50}$ value.

**Supplementary Figures (S1-S13)**

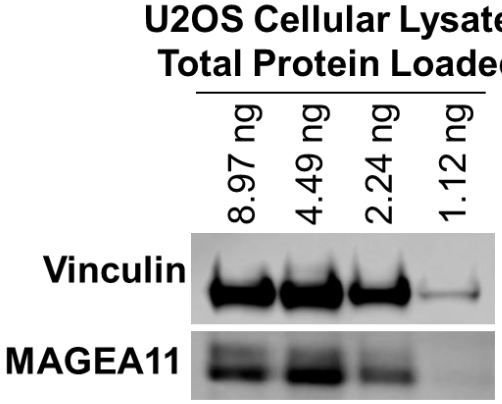

**Supplementary Figure S1.** Western blot confirmation of MAGEA11 expression in U2OS cells.

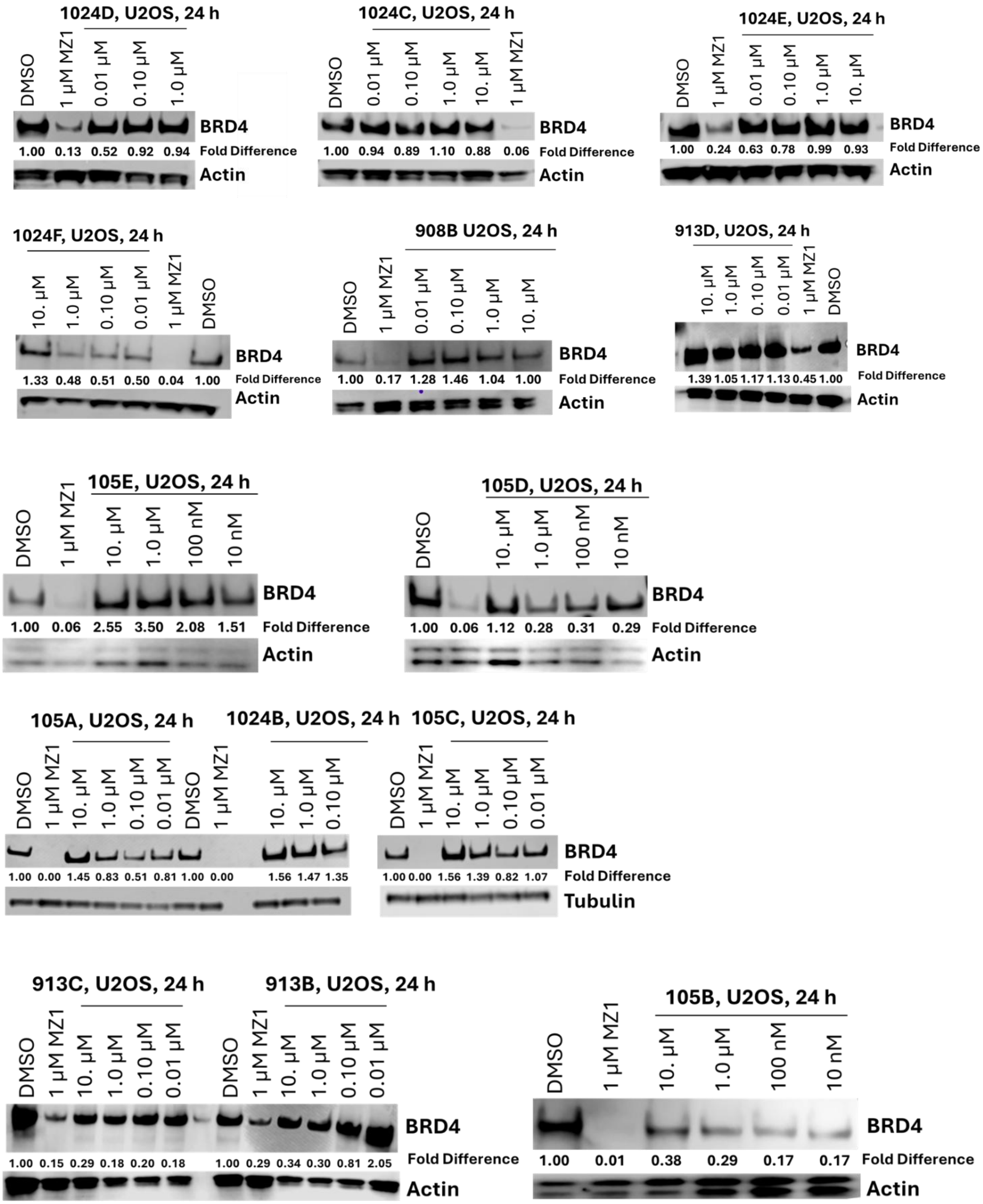

**Supplementary Figure S2.** Screening data for the PROTACs library.

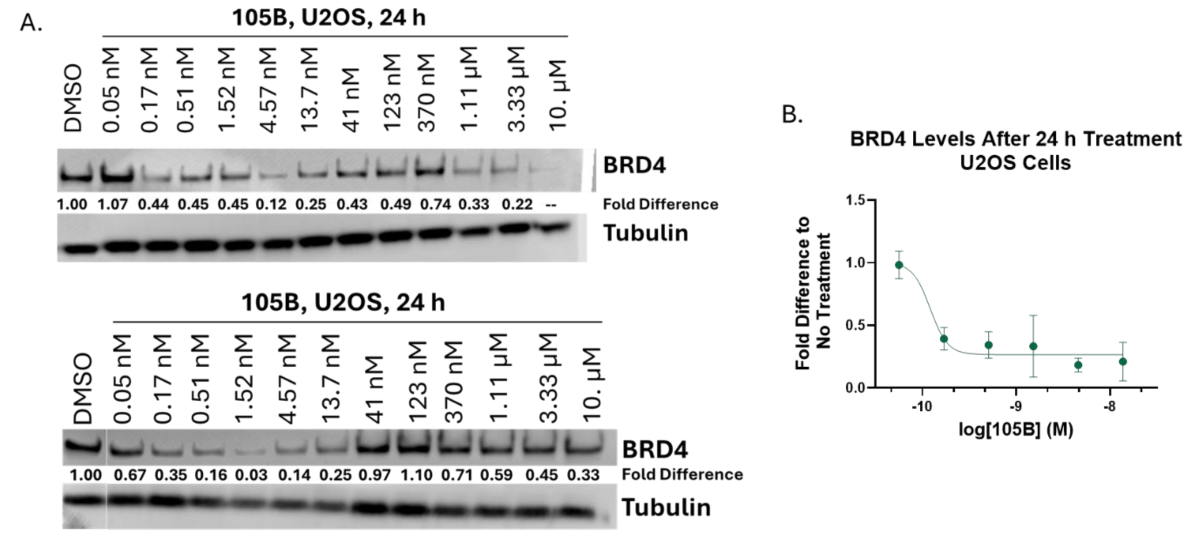

**Supplementary Figure S3.** Additional replicates of 12-point dose response curves of 105B in U2OS cells at 24 hours. A. Western blots of quantified BRD4 degradation. B. Dose-response curve used to calculate $DC_{50}$ of 105B using GraphPad Prism 10. Each point represents the mean ± standard deviation of three experimental replicates. Points past the hook effect (> 41 nM) were excluded from the analysis.

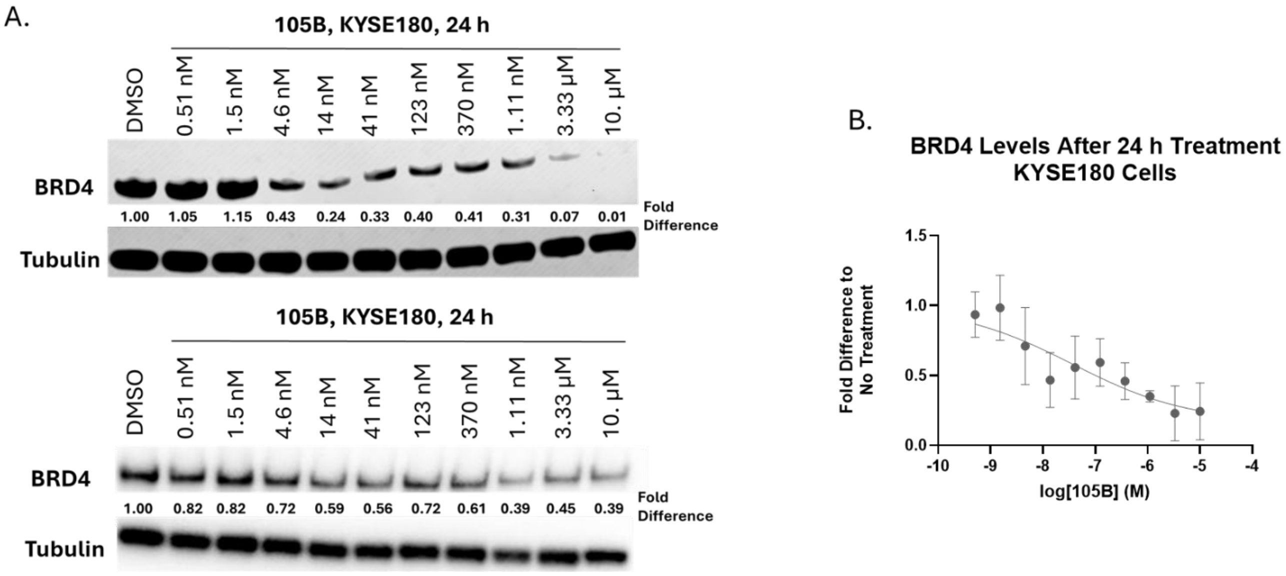

**Supplementary Figure S4.** Additional replicates of 12-point dose response curves of 105B in KYSE180 cells at 24 hours. A. Western blots of quantified BRD4 degradation. B. Dose-response curve used to calculate $DC_{50}$ of 105B using GraphPad Prism 10. Each point represents the mean ± standard deviation of three experimental replicates.

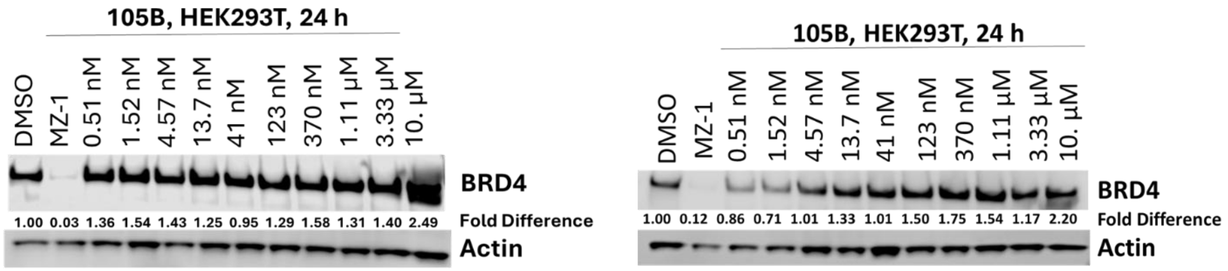

**Supplementary Figure S5.** Additional western blot replicates of 10-point dose response curves of 105B in HEK293T cells at 24 hours.

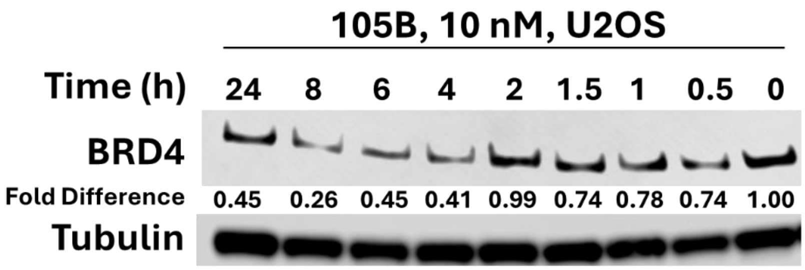

**Supplementary Figure S6**. Replication of the time course degradation experiment.

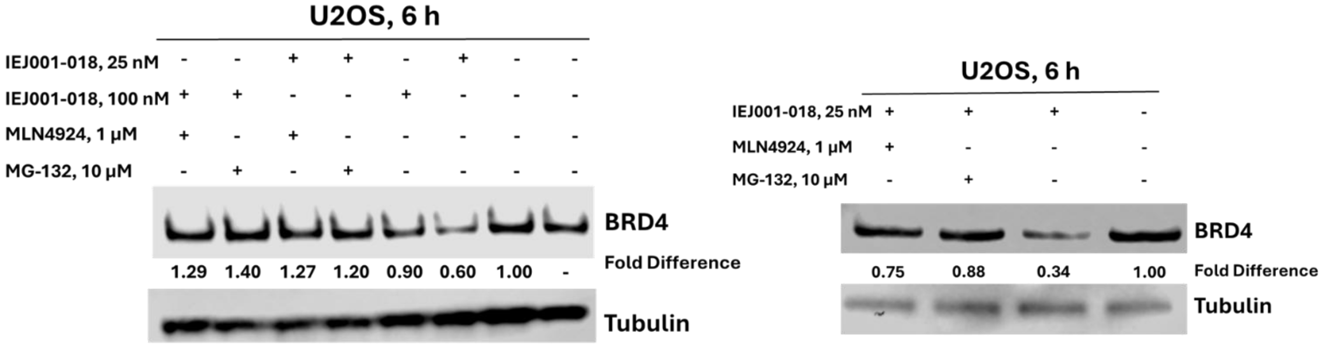

**Supplementary Figure S7.** Replicates of the MG132 and MLN4924 experiments.

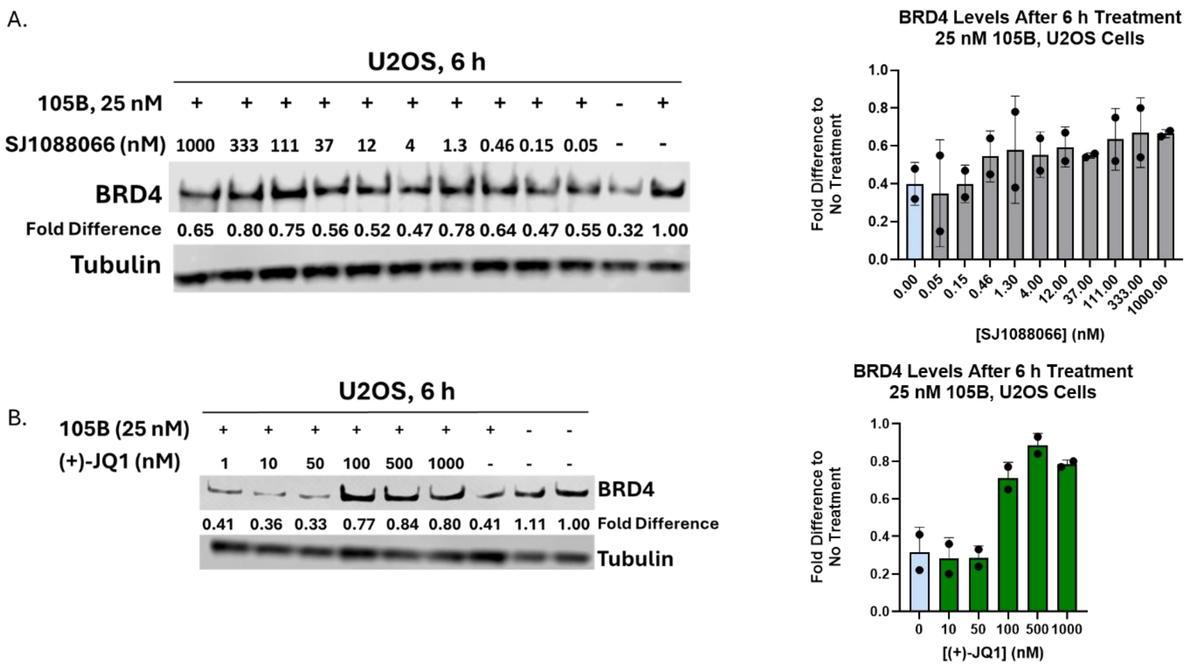

**Supplementary Figure S8**. Replicates of the ligand competition experiments. A. Replicate of the competition with increasing concentrations of SJ1088066. The graph represents the mean and standard deviation of two replicates. B. Replicate of competition with (+)-JQ1. The graph represents the mean and standard deviation of two replicates.

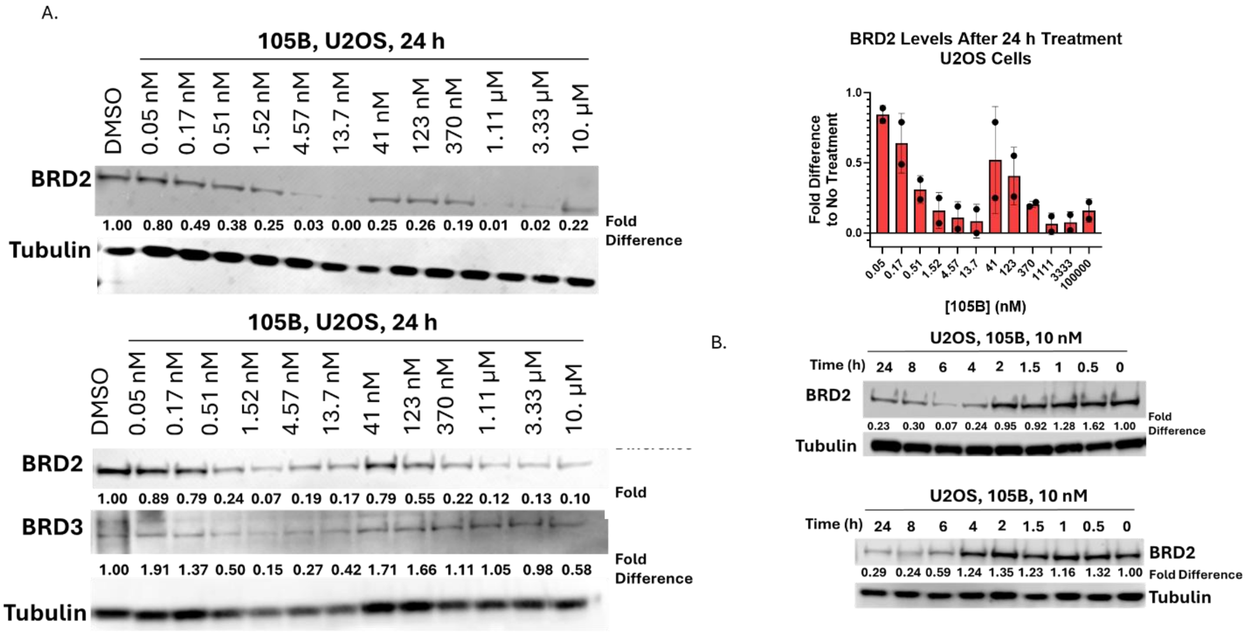

**Supplementary Figure S9.** Degradation of BRD2 and BRD3 in U2OS cells with 105B. A. Replicates of 12-point dose response of BRD2/3 degradation with 105B. The graph represents the mean of two replicates ± SD. B. Replicates of time course degradation of BRD2.

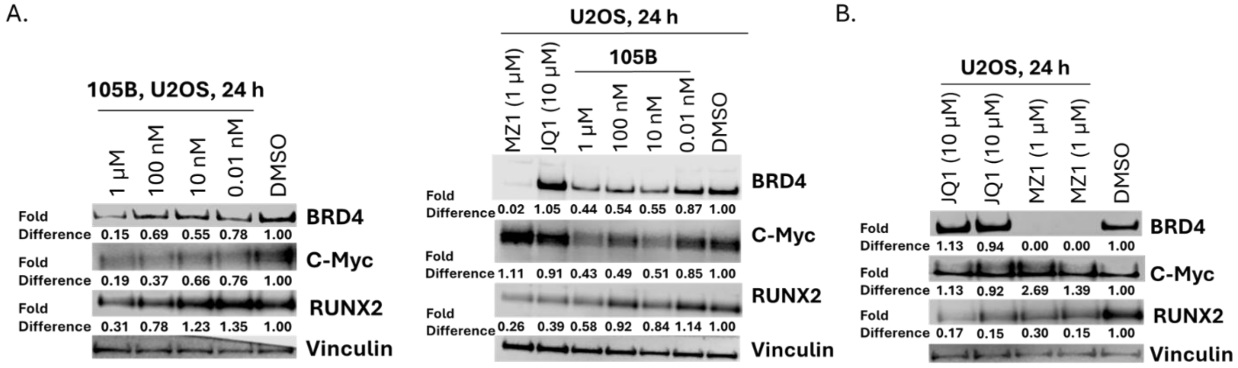

**Supplementary Figure S10.** Replicates of c-Myc and RUNX2 western blots in U2OS cells. A. Replicates of c-Myc and RUNX2 levels in U2OS cells after incubation with 105B for 24 hours. B. Replicates of c-Myc and RUNX2 levels in U2OS cells after incubation with controls (+)-JQ1 and MZ-1 for 24 hours.

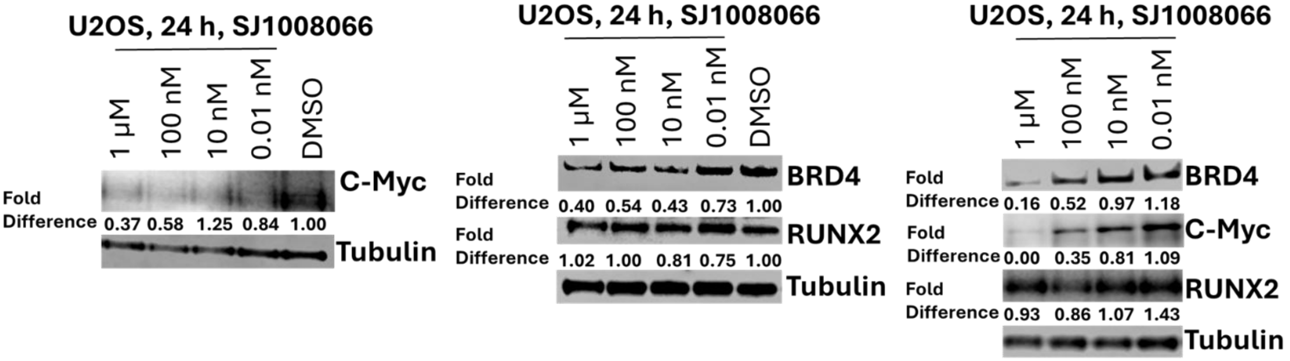

**Supplementary Figure S11.** Replicates of western blots probing levels of BRD4, c-Myc, and RUNX2 in U2OS cells treated with SJ1008066.

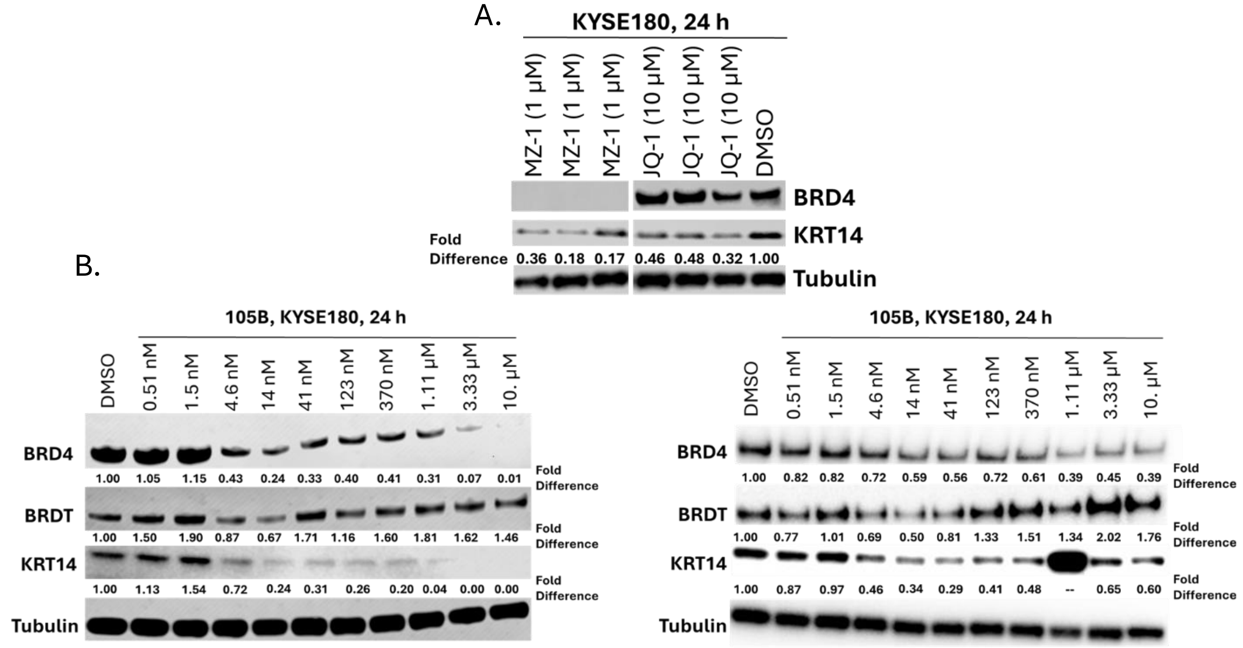

**Supplementary Figure S12.** A. Replicates of western blots of KRT14 and BRD4 levels in KYSE180 cells treated with (+)-JQ1 and MZ-1. B. Replicates of western blots probing levels of BRDT and KRT14 in KYSE180 cells. KRT14 levels in points 1.11 µM- 10 µM were not used in replicate 3 due to insoluble cell debris in the 1.11 µM lane, distorting the signal.

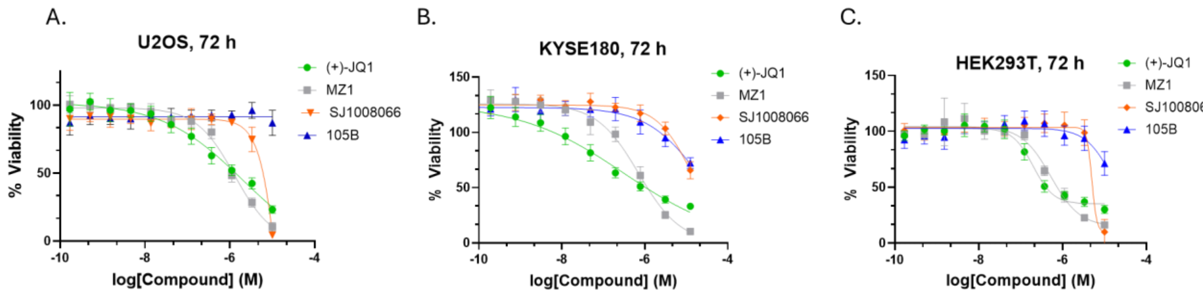

| Compound | $GI_{50}$ (μM) |
|---|---|
| (+)-JQ1 | 1.32 ± 0.27 |
| MZ1 | 1.31 ± 0.1 |
| SJ1008066 | 5.22 ± 0.39 |
| 105B | -- |

| Compound | $GI_{50}$ (μM) |
|---|---|
| (+)-JQ1 | 0.364 ± 0.147 |
| MZ1 | 0.717 ± 0.070 |
| SJ1008066 | >10 |
| 105B (% viability at 10 μM) | 72.3 ± 2.2 |

| Compound | $GI_{50}$ (μM) |
|---|---|
| (+)-JQ1 | 0.194 ± 0.020 |
| MZ1 | 0.572 ± 0.126 |
| SJ1008066 | 7.16 ± 1.85 |
| 105B (% viability at 10 μM) | 71.3 ± 9.4 |

**Supplementary Figure S13.** CellTiter-Glo cell viability data for: A. U2OS cells, B. KYSE180 cells, and C. HEK293T cells. All curves represent triplicate data as the mean ± SD.

#### Instrumentation and Materials

**Chemicals:** (+)-JQ1 acid, 7-bromo-4-chloro quinoline, and (1*S*,2*S*)-*N*-*N*'-dimethylcyclohexane-1,2-diamine were purchased from Ambeed. All amine and propyl functionalized PEG linkers and indole-5-carbaldehyde were purchased from Astatech. All other chemicals, including solvents, were purchased from Sigma Aldrich. All chemicals were used from suppliers without further purification. **Safety statement:** Sodium azide poses an explosion risk if concentrated under reduced pressure.

**HPLC:** Purity of all compounds was obtained with a C18 column (201HS54 Vydac) on an UltiMate 3000 HPLC with a 0-50% gradient of ACN in $H_2O$ w/0.1% TFA. 1 mM sample stocks in DMSO were injected onto HPLC and purity was calculated from the 254 nm wavelength UV trace.

**NMR:** NMR was measured on a Bruker Avance 700/850-MHz, Bruker Avance NEO 600-MHz, or a Bruker Avance III 400/100-MHz NMR instrument. Chemical shifts are reported in parts per million (ppm). Splitting patterns are designed as s, singlet; d, doublet; t, triplet; q, quartet; m, multiplet; br, broad singlet. Chemical shifts are reported in ppm relative to $C_2D_6SO$ (2.50 ppm for $^1H$, 39.5 ppm for $^{13}C$). NMR spectra were processed with the MestReNova or TopSpin programs.

**HRMS:** HRMS was measured on a Sciex X500R QTOF-MS in MeOH.

**LCMS:** Characterization with LCMS was done with an Agilent 1260 Infinity II instrument with a 3x150 mm C18 column with a 5-100% gradient of ACN in $H_2O$ w/0.1% formic acid.

### Synthetic Route and Characterization Data for PROTACs and SJ1008066

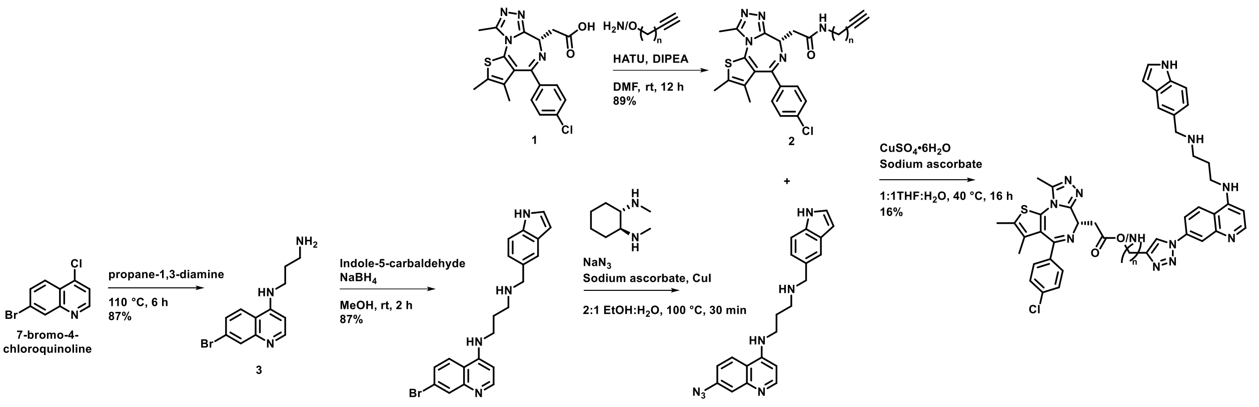

**Supplementary Scheme 1:** General Scheme for the Synthesis of PROTACs.

Intermediates used in PROTAC synthesis and an example linker for 105B:

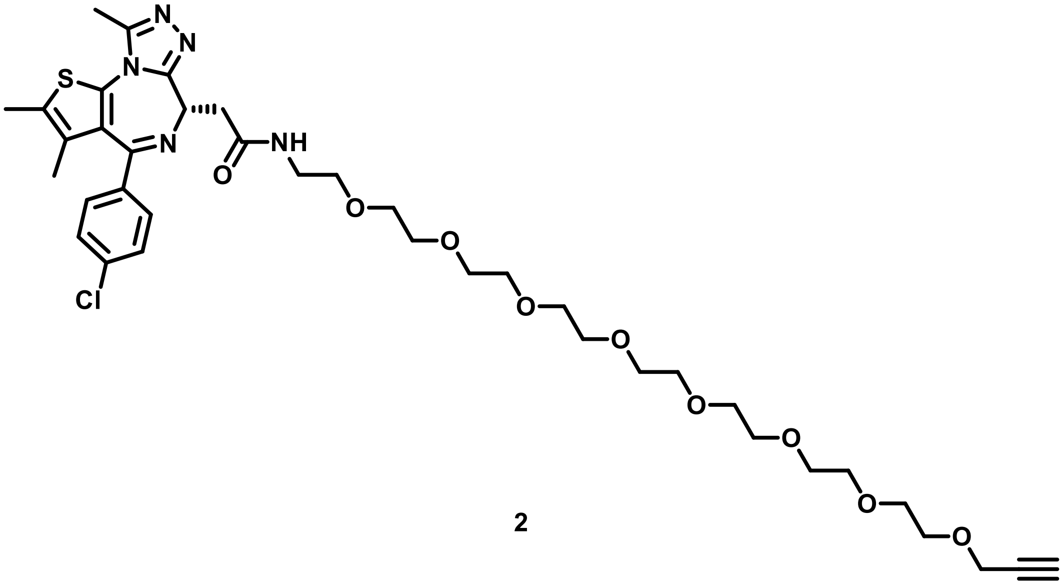

2

**(*S*)-2-(4-(4-Chlorophenyl)-2,3,9-trimethyl-6*H*-thieno[3,2-f][1,2,4]triazolo[4,3-a][1,4]diazepin-6-yl)-*N*-(3,6,9,12,15,18,21,24-octaoxaheptacos-26-yn-1-yl)acetamide (2)**. Propargyl-PEG8-amine (125 mg, 1 eq), (+)-JQ1 carboxylic acid (123 mg, 1 eq), and HATU (128 mg, 1.1 eq) were dissolved in DMF (3.07 mL, 0.1 M). DIPEA (0.160 mL, 3 eq) was added, and the reaction was stirred at room temperature for 16 h. The crude reaction mixture was diluted in $H_2O$ (5 mL). The product was extracted with DCM (3 x 10 mL), washed with brine (1 x 10 mL), and dried over anhydrous $MgSO_4$. The mixture was concentrated *in vacuo* and purified via flash column chromatography (silica gel, 0-20% MeOH in DCM w/1% TEA) to afford **2** as a yellow oil (89% yield). $^1$H NMR (400 MHz, $C_2D_6SO$ w/0.1% TMS, standard residual internal $C_2HD_5SO$ δ 2.50) δ 8.27 (br, 1H), 7.54 – 7.32 (m, 4H), 4.50 (t, *J* = 7.1 Hz, 1H), 4.13 (d, *J* = 2.5 Hz, 2H), 3.61 – 3.38 (m, 29H), 3.27-3.19 (m, 6H obscured by water), 2.66 (m, 1H), 2.59 (s, 3H), 2.41 (s, 3H), 2.32 (m, 1H), 1.62 (s, 3H). LCMS (ESI) calcd for $C_{38}H_{52}ClN_5O_9S$ 790.4, found 791.3 $[M+H]^+$.

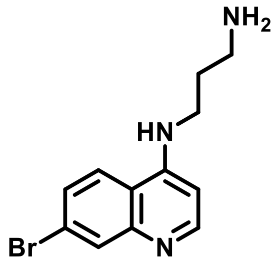

**3**

***N*$^1$-(7-Bromoquinolin-4-yl)propane-1,3-diamine (3).** 7-Bromo-4-chloroquinoline (1 g, 1 eq) was dissolved in 1,3-propanediamine (3 mL, 5.8 eq) and heated to 100 °C for 16 h. The reaction was cooled to room temperature, and 1 M NaOH (10 mL) and water (15 mL) were added, forming a white precipitate. The precipitate was filtered to afford **3** as an off-white powder. The product was used without further purification (Yield 87%). $^1$H NMR (400 MHz, $C_2D_6SO$ w/0.1% TMS, standard residual internal $C_2D_5HSO$ δ 2.50) δ 8.37 (d, *J* = 5.4 Hz, 1H), 8.15 (d, *J* = 9.0 Hz, 1H), 7.93 (d, *J* = 2.1 Hz, 1H), 7.54 (d, *J* = 8.9, 2H), 7.54 (br, 1 H, overlapping with doublet), 6.48 (d, *J* = 5.5 Hz, 1H), 3.31 (t, *J* = 7.0 Hz, 2H, obscured by water), 2.68 (t, *J* = 6.5 Hz, 2H), 1.72 (p, *J* = 6.7 Hz, 2H). LCMS (ESI) calcd for $C_{12}H_{14}BrN_3$ 280.2 found, 281.0 $[M+H]^+$.

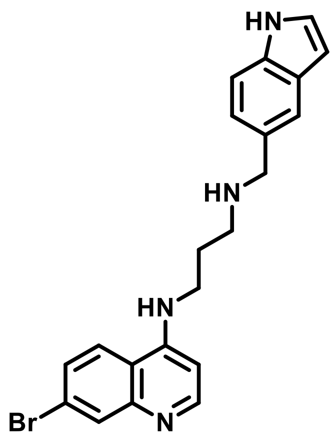

**4**

***N*[1]-((1*H*-Indol-5-yl)methyl)-*N*3-(7-bromoquinolin-4-yl)propane-1,3-diamine (4).** Compound **3** (750 mg, 1 eq) and indole-5-carbaldehyde (389 mg, 1 eq) were dissolved in MeOH (27 mL, 0.1 M) and stirred for 1 h at rt. Sodium borohydride (50.6 mg, 0.5 eq) was added, and the reaction was stirred for an additional h. The reaction was quenched with $H_2O$, and the MeOH was removed *in vacuo*. The remaining aqueous layer was extracted with ethyl acetate (3 x 10mL), washed with brine (1x10 mL), and dried over anhydrous $NaSO_4$. The reaction mixture was concentrated *in vacuo* and purified via reverse phase flash column chromatography (C18, 5-100% ACN in $H_2O$) to afford **4** as an off-white powder (87% yield). $^1$H NMR (400 MHz, $C_2D_6SO$ w/0.1% TMS, standard residual internal $C_2D_5HSO$ δ 2.50) δ 11.01 (s, 1H), 8.36 (d, *J* = 5.4 Hz, 1H), 8.01 (d, *J* = 9.0 Hz, 1H), 7.93 (d, *J* = 2.1 Hz, 1H), 7.67 (br, 1H), 7.48 (d, *J* = 1.5 Hz, 1H), 7.41 – 7.28 (m, 3H), 7.09 (dd, *J* = 8.3, 1.7 Hz, 1H), 6.45 (d, *J* = 5.5 Hz, 1H), 6.35 (t, *J* = 2.5 Hz, 1H), 3.77 (s, 2H), 3.32 (d, *J* = 13.5 Hz, 2H. Note: obscured by water), 2.69 (t, *J* = 6.4 Hz, 2H), 1.87-1.80 (m, 2H). $^{13}$C NMR (400 MHz, $C_2D_6SO$) δ 152.3, 150.6, 149.8, 135.5, 131.4, 131.2, 128.0, 126.9, 125.8, 125.8, 124.45, 122.4, 122.3, 119.8, 118.1, 111.5, 101.3, 99.0, 54.2, 47.4, 41.9, 40.6, 40.4, 40.2, 40.0, 39.8, 39.6, 39.4, 28.1. LCMS (ESI) calcd for $C_{21}H_{21}BrN_4$ 409.3, found 410.1 $[M+H]^+$

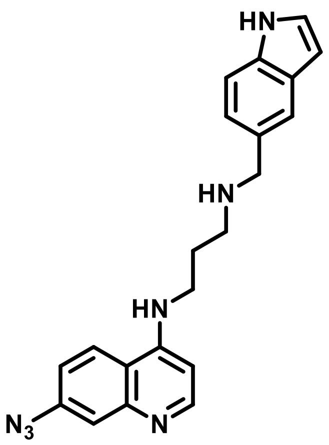

**5**

***N*$^1$-((1*H*-Indol-5-yl)methyl)-*N*3-(7-azidoquinolin-4-yl)propane-1,3-diamine (5)**. To a flask under nitrogen was added **4** (50 mg, 1 eq), sodium azide (23.8 mg, 3 eq), sodium ascorbate (2.42 mg, 0.1 eq), copper iodide (2.33 mg, 0.1 eq), and (1*S*,2*S*)-N1,N2-dimethylcyclohexane-1,2-diamine (5.21 mg, 0.3 equiv.). The reaction mixture was dissolved in ethanol and water (2:1 ratio, 2 mL, 0.01 M) and heated in a microwave reactor (100 °C) for 30 minutes. The reaction was allowed to cool to room temperature and diluted with brine (5 mL). The product was extracted with THF (3 x 10 mL), washed with brine (10 mL), and dried with anhydrous $MgSO_4$. The product was concentrated under reduced pressure and purified via reverse phase flash column chromatography (C18, 5-100% ACN in $H_2O$) to afford **6** as a white powder**.** Identity confirmed via LCMS due to the inability to completely remove solvent *in vacuo*. LCMS (ESI) calcd for $C_{21}H_{21}N_7$ 371.5, found 372.2 $[M+H]^+$.

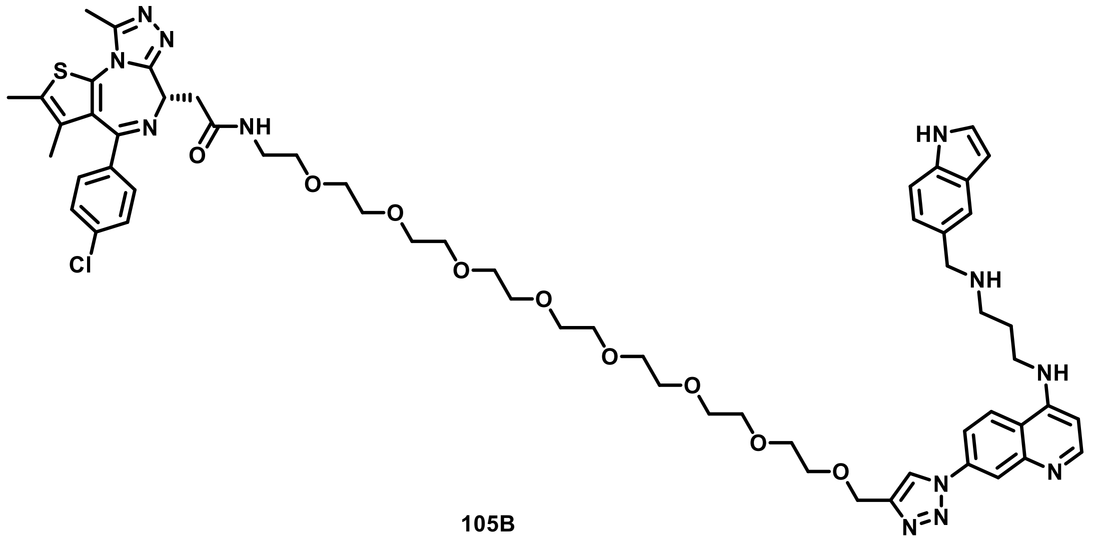

105B

**(*S*)-*N*-(1-(1-(4-((3-(((1*H*-Indol-5-yl)methyl)amino)propyl)amino)quinolin-7-yl)-1*H*-1,2,3-triazol-4-yl)-2,5,8,11,14,17,20,23-octaoxapentacosan-25-yl)-2-(4-(4-Chlorophenyl)-2,3,9-trimethyl-6*H*-thieno[3,2-f][1,2,4]triazolo[4,3-a][1,4]diazepin-6-yl)acetamide** (**105B**). Compound **5** (10 mg, 1.1 eq) and **2** (19.3 mg, 1 eq) were dissolved in $H_2O$/THF (1 mL total, 50:50 ratio, 0.18 M). $CuSO_4$•$6H_2O$ (0.6 mg, 0.1 eq) and sodium ascorbate (1 mg, 0.2 eq) were added, and the reaction was stirred at 45 °C for 16 h. The crude reaction mixture was concentrated in vacuo, diluted in MeOH (3 mL), and purified with ACCQPrep HP150 (Teledyne) equipped with a C18 column (Kinetex) (5-100% ACN in $H_2O$ w/0.1% TFA) to afford the TFA salt of **105B** as an off-white powder (16% yield). $^1$H NMR (700 MHz, , $C_2D_6SO$, standard residual internal $C_2HD_5SO$ δ 2.50), δ 11.28 (s, 1H), 9.52 (t, *J* = 6.1 Hz, 1H), 9.08 (s, 2H), 9.04 (br, 1H), 8.72 (d, *J* = 9.3 Hz, 1H), 8.65 (d, *J* = 7.1 Hz, 1H), 8.53 (d, *J* = 2.2 Hz, 1H), 8.34 – 8.27 (m, 2H), 7.67 (d, *J* = 1.5 Hz, 1H), 7.51 – 7.35 (m, 6H), 7.19 (dd, *J* = 8.3, 1.6 Hz, 1H), 6.96 (d, *J* = 7.2 Hz, 1H), 6.46 – 6.40 (m, 1H), 4.68 (s, 2H), 4.52 (dd, *J* = 8.2, 5.9 Hz, 1H), 4.23 (t, *J* = 5.6 Hz, 2H), 3.66-3.65 (m, 4H), 3.61 – 3.56 (m, 2H), 3.55 – 3.39 (m, 28H), 3.33 – 3.18 (m, 4H), 3.10 (br, 2H), 2.59 (s, 3H), 2.39 (s, 3H), 1.61 (s, 3H). $^{13}$C NMR (850 MHz, $C_2D_6SO$) δ 169.7, 163.1, 155.5, 145.9, 145.9, 143.5, 139.4, 138.6, 136.74, 136.67, 136.0, 135.4, 135.3, 132.2, 130.9, 130.2, 129.9, 129.7, 128.5, 127.7, 126.5,

126.0, 122.73, 122.68, 122.1, 121.9, 118.2, 116.1, 111.7, 109.7, 101.2, 98.8, 69.9, 69.82, 69.81, 69.77, 69.8, 69.7, 69.31, 69.25, 63.4, 53.8, 51.0, 43.8, 40.5, 38.7, 37.5, 24.4, 14.0, 12.7, 11.3. HRMS $[M+2H]^{2+}$ calcd for $C_{59}H_{73}ClN_{12}O_9S$ 581.2589. Found 581.2562. Purity: 96%.

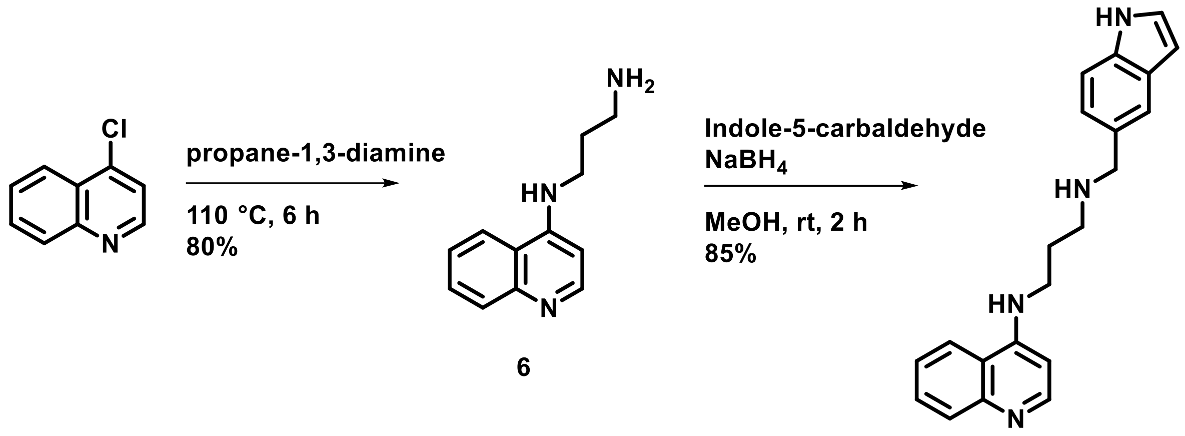

SJ1008066

**Supplementary Scheme 2:** Synthetic route to SJ1008066.

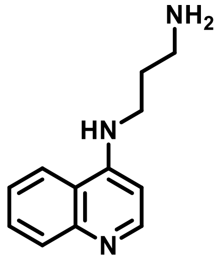

**6**

***N*[1]-(Quinolin-4-yl)propane-1,3-diamine (6).** 4-Chloroquinoline (1 g, 1 eq) was dissolved in 1,3-propanediamine (3mL, 5.8 eq) and heated to 100 °C for 16 h. The reaction was cooled to room temperature, and water (15 mL) was added, forming a white precipitate. The precipitate was filtered to afford **6** as an off-white powder. The product was used without further characterization, and its identity was confirmed via LCMS (80 % yield). LCMS(ESI): calcd for $C_{12}H_{15}N_3$ 201.3, found 202.1 $[M+H]^+$.

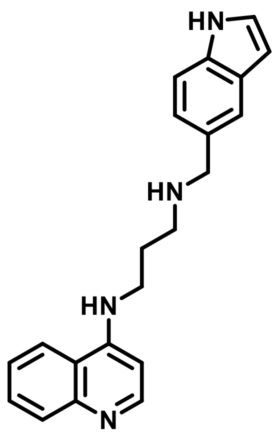

**N[1]-((1*H*-Indol-5-yl)methyl)-*N*3-(quinolin-4-yl)propane-1,3-diamine (SJ1008066).** Compound **6** (1 g, 1 eq) and indole-5-carbaldehyde (721 mg, 1 eq) were dissolved in MeOH (50 mL, 0.1 M) and stirred for 1 h at rt. Sodium borohydride (94 mg, 0.5 eq) was added, and the reaction was stirred for an additional h. The reaction was quenched with $H_2O$ (10 mL), the MeOH was removed *in vacuo,* and the remaining aqueous layer was lyophilized overnight. The reaction mixture was purified via reverse phase flash column chromatography (C18, 5-100% ACN in water) to afford **5** as an off-white powder (85%). $^1$H NMR (400 MHz, $CD_3OD$, standard residual internal $CHD_2OD$ δ 3.31) δ 8.33 (d, *J* = 5.5 Hz, 1H), 7.98 (d, *J* = 8.6 Hz, 1H), 7.79 (d, *J* = 8.5 Hz, 1H), 7.62 (t, *J* = 6.9, 1H) 7.52 (s, 1H), 7.37-7.33 (m, 2H), 7.23 (d, *J* = 3.5 Hz, 1H), 7.10 (d, *J* = 8.2 Hz, 1H), 6.50 (d, J = 5.5 Hz, 1H), 6.39 (d, *J* = 3.8 Hz, 1H), 3.89 (s, 2H), 3.43 (t, *J* = 5.9 Hz, 2H), 2.85 (t, *J* = 6.7 Hz, 2H), 2.08 – 1.87 (m, 2H). $^{13}$C NMR (600 MHz, $CD_3OD/C_2D_6SO$) δ 150.4, 149.2, 146.6, 34.8, 128.5, 128.2, 127.4, 126.7, 124.2, 123.6, 121.3, 120.4, 119.4, 118.11, 110.3, 110.3, 97.1, 52.8, 40.3, 26.4. Note: one $^1$HNMR peak expected around 4.5 ppm was obscured by the MeOH peak. HRMS $[M+H]^+$ calcd for $C_{21}H_{22}N_4$: 331.1923, found 331.1901. Purity: 97%.

<u>Characterization of PROTAC library</u>

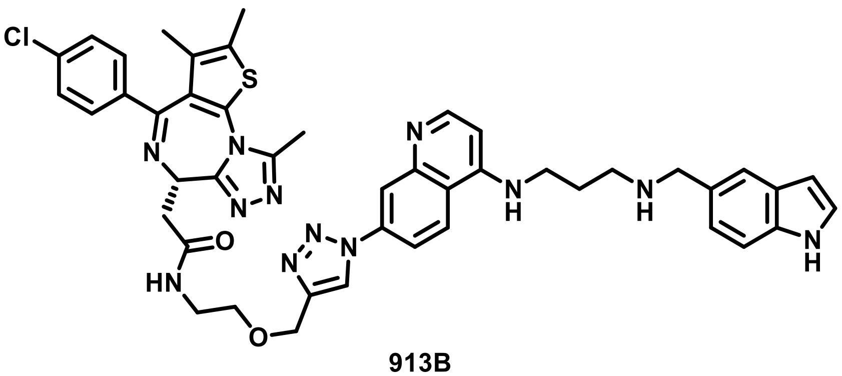

**913B**

**(*S*)-*N*-(2-((1-(4-((3-(((1*H*-Indol-5-yl)methyl)amino)propyl)amino)quinolin-7-yl)-1*H*-1,2,3-triazol-4-yl)methoxy)ethyl)-2-(4-(4-chlorophenyl)-2,3,9-trimethyl-6*H*-thieno[3,2-f][1,2,4]triazolo[4,3-a][1,4]diazepin-6-yl)acetamide (913B).** $^{1}$H NMR (400 MHz, $C_2D_6SO$ w/0.1% TMS, standard residual internal $C_2HD_5SO$ δ 2.50) δ 11.24 (s, 1H), 9.07 (s, 1H), 8.50 - 8.48(m, 2H), 8.33 (m, 2H) 8.05 (dd, $J$ = 9.1, 2.3 Hz, 1H), 7.69 (s, 1H), 7.49 – 7.34 (m, 6H), 7.23 (dd, $J$ = 8.4, 1.7 Hz, 1H), 6.60 (d, $J$ = 5.7 Hz, 1H), 6.44 (d, $J$ = 2.7 Hz, 1H), 4.70 (s, 2H), 4.51 (dd, $J$ = 8.1, 6.0 Hz, 1H), 4.20 (s, 2H), 3.59 (t, $J$ = 5.8 Hz, 2H), 3.44 (s, 2H, obscured by water), 3.25-3.16 (m, 2H, obscured by water), 3.05 (t, $J$ = 7.4 Hz, 3H), 2.58 (s, 3H), 2.38 (s, 3H), 2.00-2.01 (m, 2H), 1.59 (s, 3H), 1.24 (s, 2H). LCMS calcd for $C_{45}H_{45}ClN_{12}O_2S$: 853.4, found: 853.3 $[M]^{+}$. Purity: 98%.

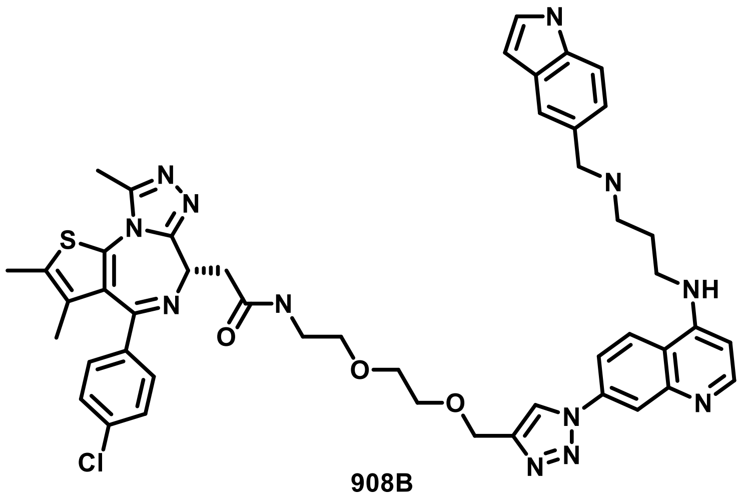

**908B**

**(*S*)-*N*-(2-(2-((1-(4-((3-(((1*H*-Indol-5-yl)methyl)amino)propyl)amino)quinolin-7-yl)-1*H*-1,2,3-triazol-4-yl)methoxy)ethoxy)ethyl)-2-(4-(4-chlorophenyl)-2,3,9-trimethyl-6*H*-thieno[3,2-f][1,2,4]triazolo[4,3-a][1,4]diazepin-6-yl)acetamide (908B)**. $^{1}H$ NMR (400 MHz, $C_2D_6SO$ w/0.1% TMS, standard residual internal $C_2HD_5SO$ δ 2.50) δ 11.25 (s, 1H), 9.11 (br, 1H), 9.02 (s, 1H), 8.57 (d, *J* = 9.2 Hz, 1H), 8.50 (d, *J* = 5.8 Hz, 1H), 8.33-8.30 (m, 2H), 8.08 (dd, *J* = 9.2, 2.4 Hz, 2H), 7.71 (s, 1H), 7.52 – 7.31 (m, 6H), 7.30 – 7.18 (m, 1H), 6.65 (d, *J* = 5.9 Hz, 1H), 6.44 (t, *J* = 2.6 Hz, 1H), 4.68 (s, 2H), 4.52 (dd, *J* = 8.0, 6.1 Hz, 1H), 4.20 (s, 2H), 3.74 – 3.55 (m, 4H), 3.51-3.46 (m, 5H), 3.27 – 3.14 (m, 4H, note: obscured by water), 3.04 (t, *J* = 7.3 Hz, 2H), 2.57 (s, 3H), 2.35 (s, 3H), 2.09 (q, *J* = 7.3 Hz, 2H), 1.56 (s, 3H). Note: 4 expected hydrogens obscured by water. LCMS calcd for $C_{47}H_{49}ClN_{12}O_3S$ 897.5, found 897.4 $[M]^+$. Purity: 98%.

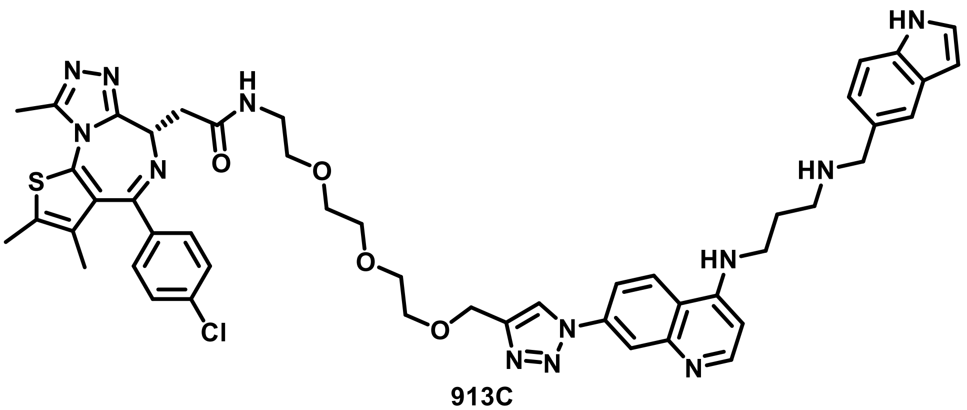

913C

**(*S*)-*N*-(2-(2-(2-((1-(4-((3-(((1*H*-Indol-5-yl)methyl)amino)propyl)amino)quinolin-7-yl)-1*H*-1,2,3-triazol-4-yl)methoxy)ethoxy)ethoxy)ethyl)-2-(4-(4-chlorophenyl)-2,3,9-trimethyl-6*H*-thieno[3,2-f][1,2,4]triazolo[4,3-a][1,4]diazepin-6-yl)acetamide (913C).** $^{1}H$ NMR (400 MHz, $C_2D_6SO$ w/0.1% TMS, standard residual internal $C_2HD_5SO$ δ 2.50) δ 11.25 (s, 1H), 9.72 (s, 1H),

9.28 (s, 2H), 9.07 (s, 1H), 8.91 (d, *J* = 9.2 Hz, 1H), 8.61 (d, *J* = 7.0 Hz, 1H), 8.53 (d, *J* = 2.2 Hz, 1H), 8.36 – 8.20 (m, 2H), 7.71 (s, 1H), 7.55 – 7.34 (m, 6H), 7.27 (dd, *J* = 8.4, 1.7 Hz, 1H), 6.93 (d, *J* = 7.1 Hz, 1H), 6.52 (s, 1H), 6.43 (s, 1H), 4.68 (s, 2H), 4.49 (dd, *J* = 8.0, 5.9 Hz, 1H), 4.19 (s, 2H), 3.72 – 3.63 (m, 4H), 3.59 (dd, *J* = 5.9, 3.3 Hz, 2H), 3.55 (s, 4H), 3.46 (t, *J* = 5.9 Hz, 3H), 3.33 (s, 16H), 3.29-3.17 (m, 4H, obscured by water), 3.05 (s, 2H), 2.57 (s, 3H), 2.38 (s, 3H), 2.12 (d, *J* = 7.1 Hz, 2H), 1.59 (s, 3H), 1.23 (s, 1H). Calcd for $C_{49}H_{53}ClN_{12}O_4S$ 941.6, found 941.3 $[M]^+$. Purity: 98%.

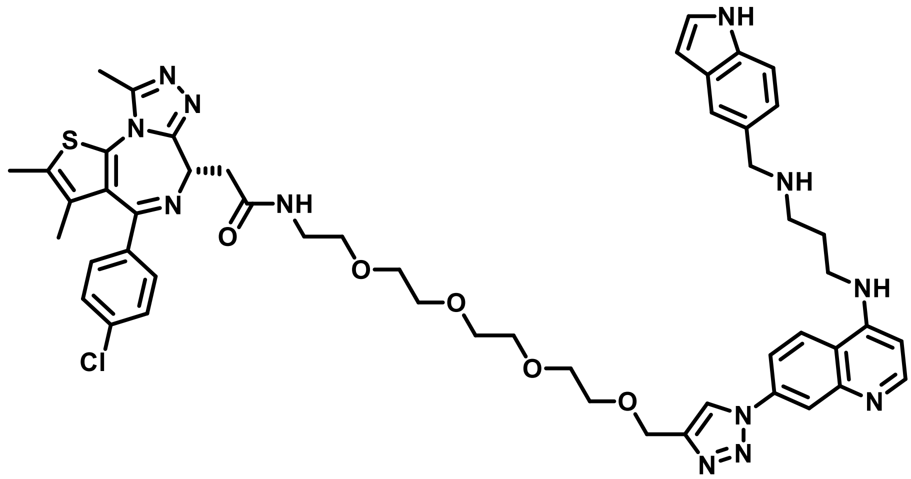

**1024F**

**(*S*)-*N*-(1-(1-(4-((3-(((1*H*-Indol-5-yl)methyl)amino)propyl)amino)quinolin-7-yl)-1*H*-1,2,3-triazol-4-yl)-2,5,8,11-tetraoxatridecan-13-yl)-2-(4-(4-Chlorophenyl)-2,3,9-trimethyl-6*H*-thieno[3,2-f][1,2,4]triazolo[4,3-a][1,4]diazepin-6-yl)acetamide (1024F).** $^1$H NMR (400 MHz, $C_2D_6SO$ w/0.1% TMS, standard residual internal $C_2HD_5SO$ δ 2.50) δ 11.24 (s, 1H), 9.62 (br, 1H), 9.22 (br, 2H), 9.07 (s, 1H), 8.88 (d, *J* = 9.2 Hz, 1H), 8.61 (d, *J* = 6.9 Hz, 1H), 8.52 (d, *J* = 2.3 Hz, 1H), 8.31 – 8.20 (m, 2H), 7.71 (s, 1H), 7.50 – 7.37 (m, 8H), 7.26 (d, *J* = 8.5 Hz, 1H), 6.92 (d, *J* = 7.0 Hz, 1H), 6.52 (s, 1H), 6.43 (t, *J* = 2.5 Hz, 1H), 4.67 (s, 2H), 4.49 (dd, *J* = 8.1, 6.0 Hz, 1H), 4.20 (s, 2H), 3.69-3.63 (m, 5H, note: integral inflated by water), 3.59 – 3.56 (m, 3H), 3.52 (s, 10H),

3.45 (t, $J$ = 5.9 Hz, 5H; integral inflated by water), 3.25 – 3.17 (m, 3H), 3.05 (s, 3H), 2.58 (s, 3H), 2.39 (s, 3H), 2.11-2.09 (m, 2H), 1.60 (s, 3H). LCMS calcd for $C_{51}H_{57}ClN_{12}O_5S$ 985.6, found 986.4 $[M+H]^+$. Purity: 89%.

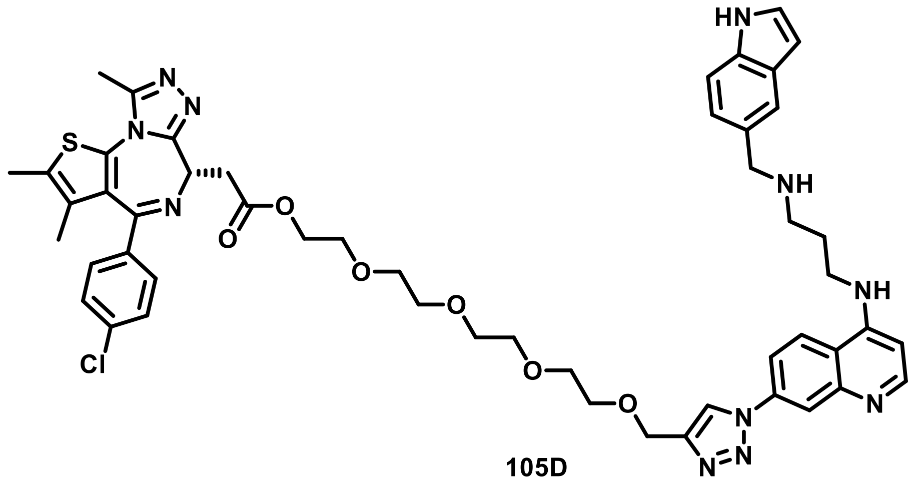

105D

**1-(1-(4-((3-(((1*H*-Indol-5-yl)methyl)amino)propyl)amino)quinolin-7-yl)-1*H*-1,2,3-triazol-4-yl)-2,5,8,11-tetraoxatridecan-13-yl (*S*)-2-(4-(4-Chlorophenyl)-2,3,9-trimethyl-6*H*-thieno[3,2-f][1,2,4]triazolo[4,3-a][1,4]diazepin-6-yl)acetate (105D).** $^1$H NMR (400 MHz, $C_2D_6SO$ w/0.1% TMS, standard residual internal $C_2HD_5SO$ δ 2.50) δ 11.24 (s, 1H), 9.79 (br, 1H), 9.26 (br, 1H), 9.07 (s, 1H), 8.91 (d, $J$ = 9.2 Hz, 1H), 8.63 (d, $J$ = 6.9 Hz, 1H), 8.54 (s, 1H), 8.27-8.25 (m, 1H), 7.71 (s, 1H), 7.56 – 7.33 (m, 6H), 7.27 (d, $J$ = 8.4 Hz, 1H), 6.95 (d, $J$ = 7.2 Hz, 1H), 6.52-6.48 (m, 2H), 4.67 (s, 2H), 4.48 (t, $J$ = 7.2 Hz, 1H), 4.23 – 4.16 (m, 3H), 3.66 – 3.45 (m, 18H; integral obscured by water), 3.07-3.03 (m, 2H), 2.59 (s, 3H), 2.41 (s, 3H), 2.13-2.10 (m, 2H), 1.62 (s, 3H), 1.23 (s, 1H). LCMS calcd for $C_{51}H_{56}ClN_{11}O_6S$ 986.6, found 1008.3 $[M + Na]^+$. Purity: 97%.

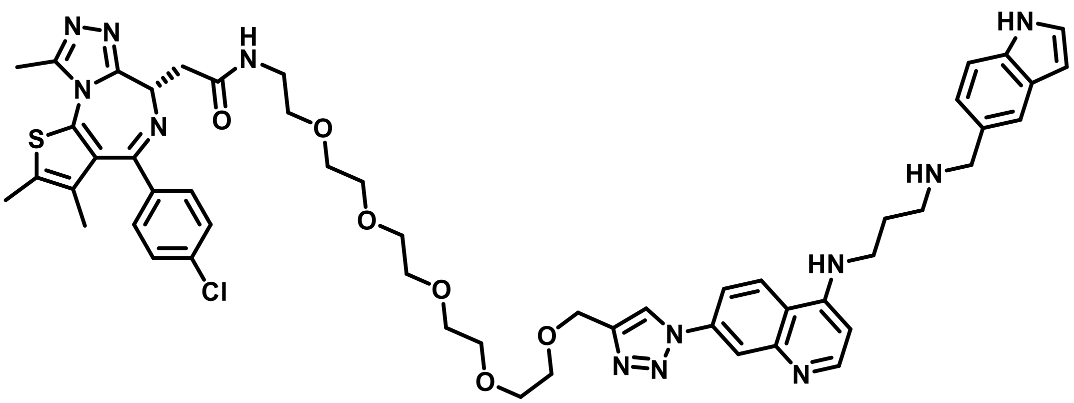

**913D**

**(*S*)-*N*-(1-(1-(4-((3-(((1*H*-Indol-5-yl)methyl)amino)propyl)amino)quinolin-7-yl)-1*H*-1,2,3-triazol-4-yl)-2,5,8,11,14-pentaoxahexadecan-16-yl)-2-(4-(4-Chlorophenyl)-2,3,9-trimethyl-6*H*-thieno[3,2-f][1,2,4]triazolo[4,3-a][1,4]diazepin-6-yl)acetamide (913D).** $^{1}$H NMR (400 MHz, $C_2D_6SO$ w/0.1% TMS, standard residual internal $C_2HD_5SO$ δ 2.50) δ 11.25 (s, 1H), 9.17 (br, 1H), 9.06 (s, 1H), 8.77 (d, *J* = 9.1 Hz, 1H), 8.57 (d, *J* = 6.5 Hz, 1H), 8.45 (d, *J* = 2.2 Hz, 1H), 8.28 (t, *J* = 5.6 Hz, 1H), 8.19 (dd, *J* = 9.1, 2.3 Hz, 1H), 7.71 (s, 1H), 7.51 – 7.34 (m, 6H), 7.26 (dd, *J* = 8.4, 1.7 Hz, 1H), 6.82 (d, *J* = 6.5 Hz, 1H), 6.43 (t, *J* = 2.5 Hz, 1H), 4.67 (s, 2H), 4.50 (dd, *J* = 8.1, 6.0 Hz, 1H), 4.20 (s, 2H), 3.65 (dd, *J* = 5.8, 3.4 Hz, 2H), 3.63 – 3.55 (m, 4H), 3.55 – 3.48 (m, 12H), 3.44 (t, *J* = 5.9 Hz, 2H; obscured by water), 3.26 – 3.20 (m, 4H: obscured by water), 3.06-3.03 (m, 2H), 2.58 (s, 3H), 2.39 (s, 3H), 2.10 (m, 2H), 1.60 (s, 3H). LCMS calcd for $C_{53}H_{61}ClN_{12}O_6S$ 1029.6, found 1029.4 $[M]^{+}$. Purity: 97%.

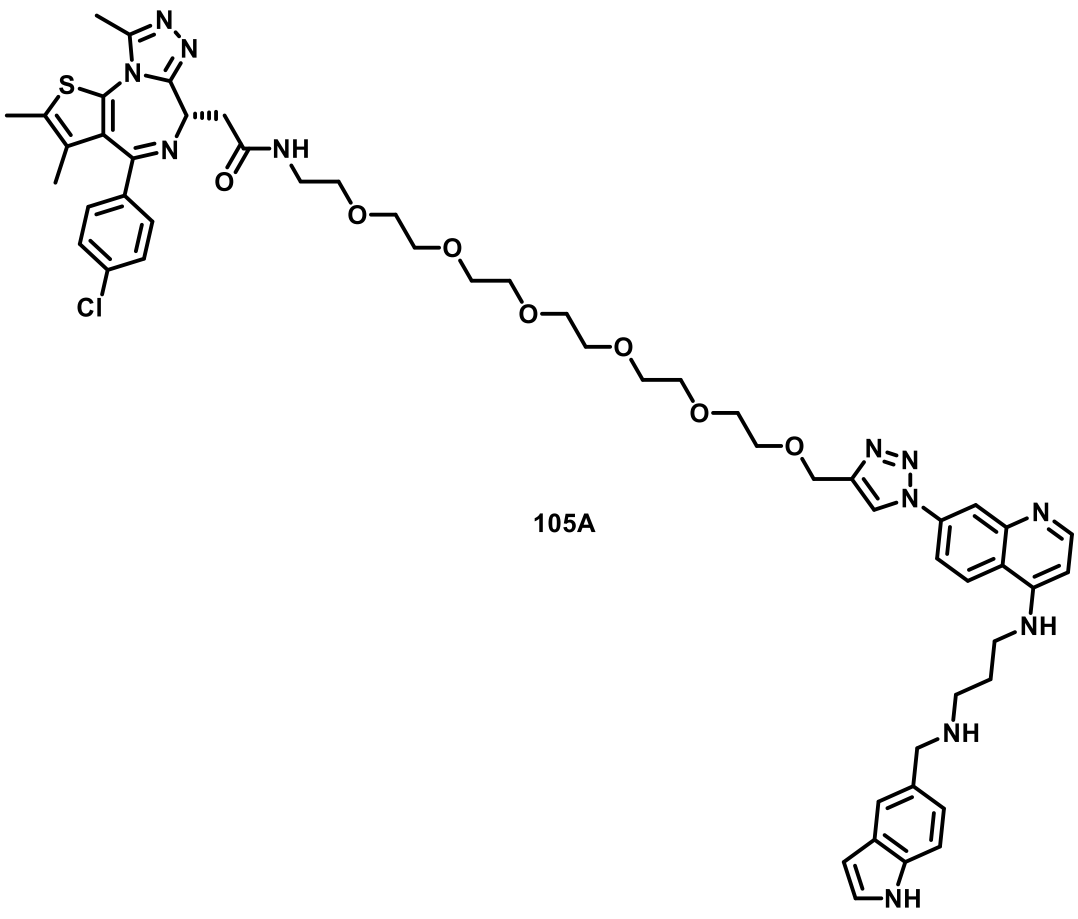

105A

**(*S*)-*N*-(1-(1-(4-((3-(((1*H*-Indol-5-yl)methyl)amino)propyl)amino)quinolin-7-yl)-1*H*-1,2,3-triazol-4-yl)-2,5,8,11,14,17-hexaoxanonadecan-19-yl)-2-(4-(4-Chlorophenyl)-2,3,9-trimethyl-6*H*-thieno[3,2-f][1,2,4]triazolo[4,3-a][1,4]diazepin-6-yl)acetamide (105A).** $^{1}$H NMR (400 MHz, $C_2D_6SO$ w/0.1% TMS, standard residual internal $C_2HD_5SO$ δ 2.50) δ 11.25 (s, 1H), 9.82 (br, 1H), 9.27 (br, 1H), 9.08 (s, 1H), 8.93 (d, *J* = 9.2 Hz, 1H), 8.69 – 8.41 (m, 2H), 8.30-8.26 (m, 2H), 7.57 – 7.35 (m, 6H), 7.28 (d, *J* = 8.5 Hz, 1H), 6.95 (d, *J* = 7.1 Hz, 1H), 6.48 (m, 1H), 4.68 (s, 2H), 4.50 (t, *J* = 7.2 Hz, 1H), 4.20 (s, 2H), 3.71 – 3.62 (m, 4H), 3.60 – 3.55 (m, 3H), 3.54 – 3.45 (m, 16H), 3.44 (m, 5H, note: obscured by water), 3.06 (m, 2H), 2.59 (s, 3H), 2.40 (s, 3H),

2.12 (m, 2H), 1.61 (s, 3H), 1.24 (s, 2H). LCMS calcd for $C_{55}H_{65}ClN_{12}O_7S$ 1073.7, found 1095.3 $[M+Na]^+$. Purity: 96%.

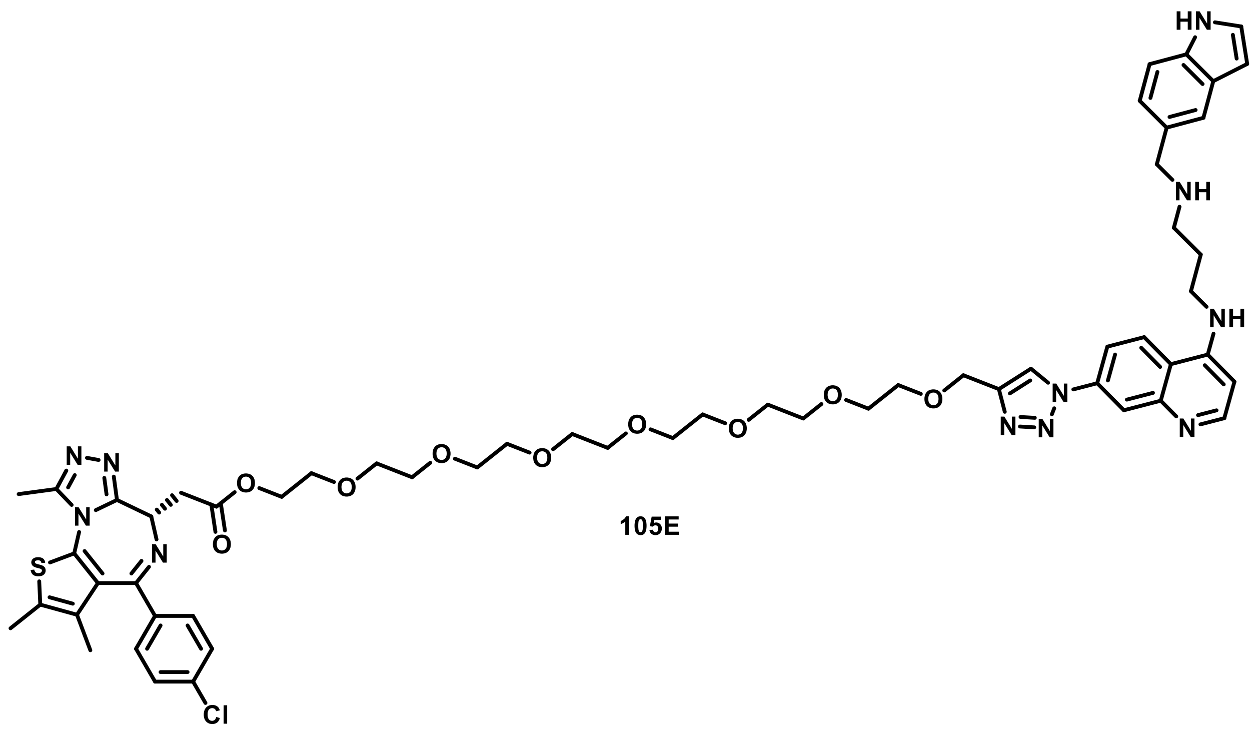

**105E**

**1-(1-(4-((3-(((1*H*-Indol-5-yl)methyl)amino)propyl)amino)quinolin-7-yl)-1*H*-1,2,3-triazol-4-yl)-2,5,8,11,14,17,20-heptaoxadocosan-22-yl (*S*)-2-(4-(4-Chlorophenyl)-2,3,9-trimethyl-6*H*-thieno[3,2-f][1,2,4]triazolo[4,3-a][1,4]diazepin-6-yl)acetate (105E).** $^1$H NMR (400 MHz, $C_2D_6SO$ w/0.1% TMS, standard residual internal $C_2HD_5SO$ δ 2.50) δ 11.24 (s, 1H), 9.66 (br 1H), 9.15 (s, 1H), 9.07 (s, 1H), 8.87 (d, *J* = 9.1 Hz, 1H), 8.63 (d, *J* = 6.8 Hz, 1H), 8.51 (d, *J* = 2.1 Hz, 1H), 8.27 (d, *J* = 8.8 Hz, 1H), 7.71 (s, 1H), 7.56 – 7.33 (m, 6H), 7.32 – 7.20 (m, 1H), 6.94 (d, *J* = 7.0 Hz, 1H), 6.52 (s, 1H), 4.68 (s, 2H), 4.48 (t, *J* = 7.1 Hz, 1H), 4.26 – 4.13 (m, 4H), 3.69 – 3.54 (m, 10H), 3.50-3.47 (m, 21H, note: integral inflated by water), 3.06 (br, 2H), 2.59 (s, 3H), 2.41 (ts, 3H), 2.10 (br, 2H), 1.62 (s, 3H), 1.23 (s, 2H). LCMS calcd for $C_{57}H_{68}ClN_{11}O_9S$ 1118.75, found 1140.3 $[M + Na]^+$. Purity: 91%.

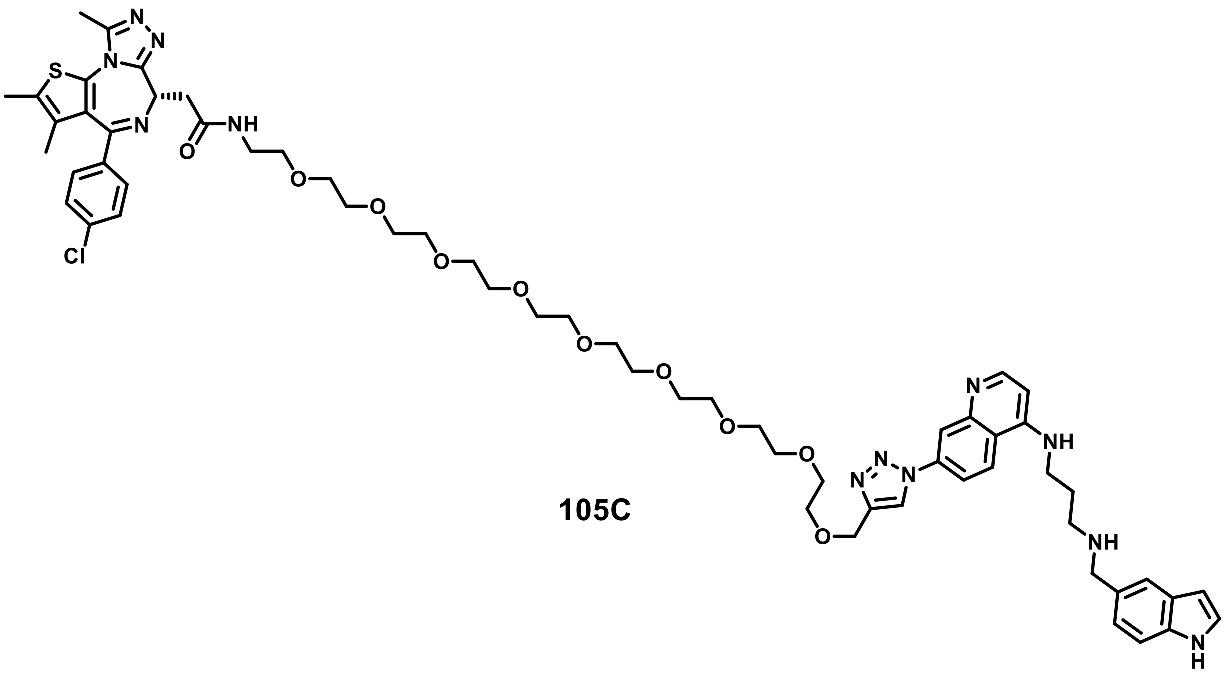

**105C**

**(*S*)-*N*-(1-(1-(4-((3-(((1*H*-Indol-5-yl)methyl)amino)propyl)amino)quinolin-7-yl)-1*H*-1,2,3-triazol-4-yl)-2,5,8,11,14,17,20,23,26-nonaoxaoctacosan-28-yl)-2-(4-(4-Chlorophenyl)-2,3,9-trimethyl-6*H*-thieno[3,2-f][1,2,4]triazolo[4,3-a][1,4]diazepin-6-yl)acetamide (105C).** $^1$H NMR (400 MHz, $C_2D_6SO$ w/0.1% TMS, standard residual internal $C_2HD_5SO$ δ 2.50) δ 11.26 (s, 1H), 9.90 (br, 1H), 9.38 (br, 1H), 9.02 (m, 2H), 8.65 – 8.43 (m, 2H), 8.29 (t, *J* = 6.6 Hz, 1H), 8.15 (br, 1H), 7.56 – 7.27 (m, 7H), 6.95 (dd, *J* = 7.3, 3.2 Hz, 1H), 6.47 (d, *J* = 39.7 Hz, 1H), 6.92 (s, 1H), 6.42 (s, 1H), 4.67 (d, *J* = 7.3 Hz, 2H), 4.50 (t, *J* = 7.0 Hz, 1H), 4.20 (s, 2H), 3.68 – 3.63 (m, 3H), 3.55 – 3.41 (m, 32H), 3.05 (m, 3H), 2.58 (s, 3H), 2.40 (d, *J* = 5.2 Hz, 3H), 2.13 (s, 3H), 1.61 (d, *J* = 5.1 Hz, 3H), 1.30 – 1.16 (m, 1H). Note: 8 hydrogens obscured by water. LCMS calcd for $C_{61}H_{77}ClN_{12}O_{10}S$ 1205.9, found 1205.5 $[M]^+$. Purity: 96%.

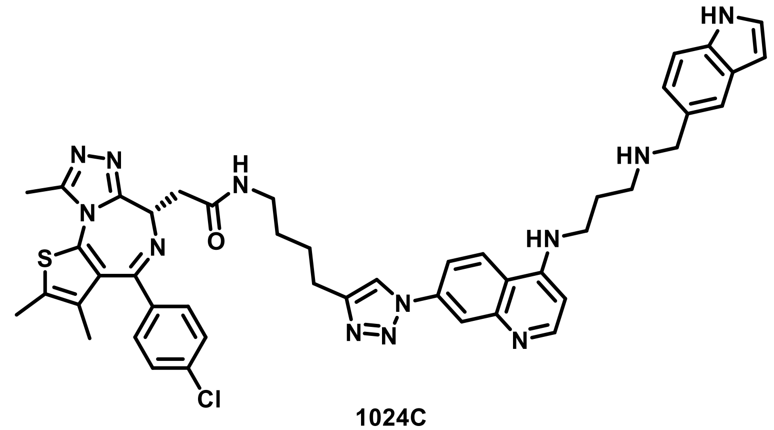

**1024C**

**(*S*)-*N*-(4-(1-(4-((3-(((1*H*-Indol-5-yl)methyl)amino)propyl)amino)quinolin-7-yl)-1*H*-1,2,3-triazol-4-yl)butyl)-2-(4-(4-Chlorophenyl)-2,3,9-trimethyl-6*H*-thieno[3,2-f][1,2,4]triazolo[4,3-a][1,4]diazepin-6-yl)acetamide (1024C).** $^{1}$H NMR (400 MHz, $C_2D_6SO$ w/0.1% TMS, standard residual internal $C_2HD_5SO$ δ 2.50) δ 11.16 (1H, s), 8.74 (1H, s), 8.42-8.36 (2H, m) 8.19-8.14 (2H, m)f, 7.93- 7.91 (1H, dd), 7.61 (1H, s), 7.41-7.31 (5H, m) 7.17- 7.15 (1H, dd), 6.48-6.47 (1H, d) 6.36 (1H, s), 4.47-4.44 (1H, t), 4.09 (2H, s), 3.21 (appx. 5H, obscured by water), 2.92 (2H, t), 2.70 (2H, t), 2.53 (3H, s) 2.30 (3H, s) 2.01- 1.94 (2H, m), 1.84- 1.65 (2H, m), 1.54-1.47 (1H, m), 1.50 (3H, s). $^{13}$C NMR (600 MHz, $C_2D_6SO$ w/0.1% TMS, standard residual internal $C_2HD_5SO$) δ 169.9, 163.5, 155.6, 152.5, 150.4, 150.3, 148.7, 137.3, 137.2, 136.3, 135.7, 132.7, 131.1, 130.6, 130.3, 130.0, 128.9, 128.0, 126.8, 124.7, 123.2, 122.3, 120.8, 118.6, 116.2, 112.0, 101.6, 54.4, 51.8, 40.6, 40.4, 40.2, 40.0, 39.8, 39.6, 39.4, 38.7, 38.2, 29.2, 26.5, 25.2, 14.5, 13.1, 11.8. LCMS calcd for $C_{46}H_{47}ClN_{12}OS$ 851.5, found 851.3 $[M]^{+}$. Purity: 97%.

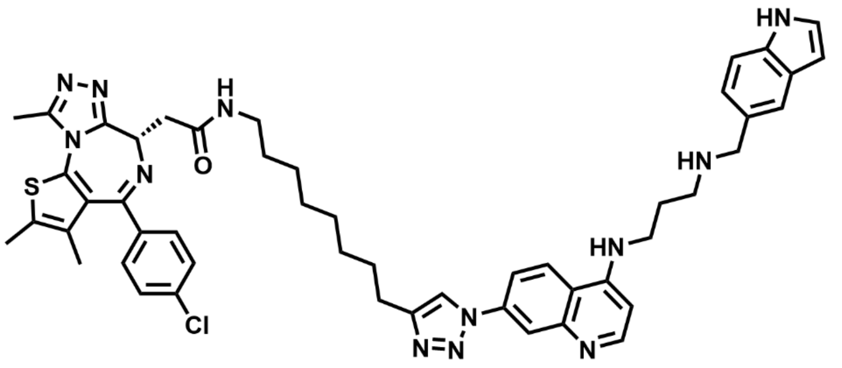

**(*S*)-*N*-(8-(1-(4-((3-(((1*H*-Indol-5-yl)methyl)amino)propyl)amino)quinolin-7-yl)-1*H*-1,2,3-triazol-4-yl)octyl)-2-(4-(4-Chlorophenyl)-2,3,9-trimethyl-6*H*-thieno[3,2-f][1,2,4]triazolo[4,3-a][1,4]diazepin-6-yl)acetamide (1024B).** $^{1}$H NMR (400 MHz, $C_2D_6SO$ w/0.1% TMS, standard residual internal $C_2HD_5SO$ δ 2.50) δ 11.24 (s, 1H), 9.10 (br, 2H), 8.81 (s, 1H), 8.70 (d, *J* = 9.1 Hz, 1H), 8.56 (d, *J* = 6.3 Hz, 1H), 8.39 (d, *J* = 2.2 Hz, 1H), 8.16 (m, 2H), 7.70 (s, 1H), 7.56 – 7.34 (m, 6H), 7.25 (dd, *J* = 8.3, 1.7 Hz, 1H), 6.77 (d, *J* = 6.4 Hz, 1H), 6.44 (t, *J* = 2.5 Hz, 1H), 4.50 (dd, *J* = 8.1, 5.9 Hz, 1H), 4.20 (s, 2H), 3.56 (s, 2H), 3.24 – 3.01 (m, 6H obscured by water), 2.73 (t, *J* = 7.5 Hz, 2H), 2.58 (s, 3H), 2.39 (s, 3H), 2.09 (m, 2H), 1.68 (t, *J* = 7.4 Hz, 2H), 1.60 (s, 3H), 1.45 (m, 2H), 1.31 (m, 9H). LCMS calcd for $C_{50}H_{55}ClN_{12}OS$ 907.6, found 907.4 $[M]^{+}$. Purity: 98%

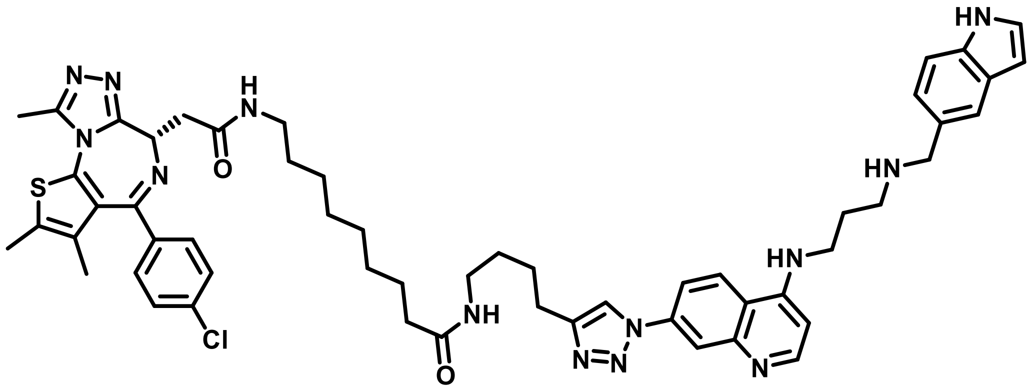

**1024E**

**(*S*)-*N*-(4-(1-(4-((3-(((1*H*-Indol-5-yl)methyl)amino)propyl)amino)quinolin-7-yl)-1*H*-1,2,3-triazol-4-yl)butyl)-9-(2-(4-(4-Chlorophenyl)-2,3,9-trimethyl-6*H*-thieno[3,2-f][1,2,4]triazolo[4,3-a][1,4]diazepin-6-yl)acetamido)nonanamide (1024E).** $^{1}$H NMR (400 MHz, $C_2D_6SO$ w/0.1% TMS, standard residual internal $C_2HD_5SO$ δ 2.50) δ 11.24 (s, 1H), 9.17 (br, 2H), 8.82 (d, *J* = 11.0 Hz, 2H), 8.59 (d, *J* = 6.7 Hz, 1H), 8.46 (s, 1H), 8.19 (mi, 2H), 7.77 (d, *J* = 5.8 Hz, 1H), 7.71 (s, 1H), 7.49 – 7.37 (m, 6H), 7.26 (d, *J* = 8.4 Hz, 1H), 6.87 (d, *J* = 6.8 Hz, 1H), 6.43 (d, *J* = 3.0 Hz, 1H), 4.49 (dd, *J* = 8.1, 5.9 Hz, 1H), 4.2920(s, 2H), 3.63 (m, 2H), 3.18 – 2.97 (m, 7H), 2.75 (t, *J* = 7.4 Hz, 2H), 2.58 (s, 3H), 2.40 (s, 3H), 2.14 – 2.00 (m, 4H), 1.74 – 1.64 (m, 2H), 1.60 (s, 3H), 1.54 – 1.35 (m, 6H), 1.22 (d, *J* = 6.3 Hz, 8H). $^{13}$C NMR (600 MHz, $C_2D_6SO$ w/0.1% TMS) δ 172.4, 169.8, 163.4, 155.6, 150.3, 149.1, 139.3, 137.2, 136.4, 135.7, 132.7, 131.2, 130.6, 130.3, 130.0, 128.9, 128.0, 126.9, 126.4, 123.4, 122.7, 122.4, 121.0, 117.9, 116.9, 112.0, 101.6, 99.2, 54.4, 51.3, 44.2, 40.6, 40.4, 40.2, 40.0, 39.8, 39.6, 39.4, 38.9, 38.5, 38.1, 35.9, 35.9, 29.7, 29.1, 26.8, 26.6, 25.8, 25.1, 24.7, 14.5, 13.2, 11.8. LCMS calcd for $C_{55}H_{64}ClN_{13}O_2S$ 1006.7, found 1006.5 $[M]^+$. Purity: 97%.

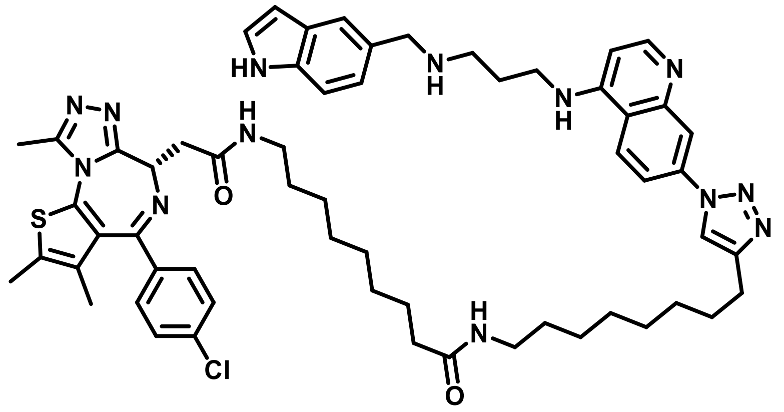

**1024D**

**(*S*)-*N*-(8-(1-(4-((3-(((1*H*-Indol-5-yl)methyl)amino)propyl)amino)quinolin-7-yl)-1*H*-1,2,3-triazol-4-yl)octyl)-9-(2-(4-(4-Chlorophenyl)-2,3,9-trimethyl-6*H*-thieno[3,2-f][1,2,4]triazolo[4,3-a][1,4]diazepin-6-yl)acetamido)nonanamide (1024D).** $^1$H NMR (400 MHz, $C_2D_6SO$ w/0.1% TMS, standard residual internal $C_2HD_5SO$ δ 2.50) δ 11.26 (s, 1H), 9.18 (br, 2H), 8.81 (s, 1H), 8.69 (d, *J* = 9.2 Hz, 1H), 8.53 (d, *J* = 6.2 Hz, 1H), 8.37 (d, *J* = 2.3 Hz, 1H), 8.26 – 8.01 (m, 2H), 7.72 (m, 2H), 7.55 – 7.36 (m, 6H), 7.27 (dd, *J* = 8.4, 1.7 Hz, 1H), 6.73 (d, *J* = 6.3 Hz, 1H), 6.44 (t, *J* = 2.4 Hz, 1H), 4.50 (dd, *J* = 8.1, 5.9 Hz, 1H), 4.20 (s, 2H), 3.55 (d, *J* = 7.0 Hz, 2H), 3.03 (m, predicted 5H obscured by water), 2.72 (t, *J* = 7.5 Hz, 2H), 2.58 (s, 3H), 2.39 (s, 3H), 2.09 (t, *J* = 7.3 Hz, 2H), 2.01 (t, *J* = 7.3 Hz, 2H), 1.67 (q, *J* = 7.4 Hz, 2H), 1.60 (s, 3H), 1.50 – 1.12 (m, 24H). LCMS calcd for $C_{59}H_{72}ClN_{13}O_2S$ 1062.8, found 1062.6 $[M]^+$. Purity: 96%.

### NMR Spectra

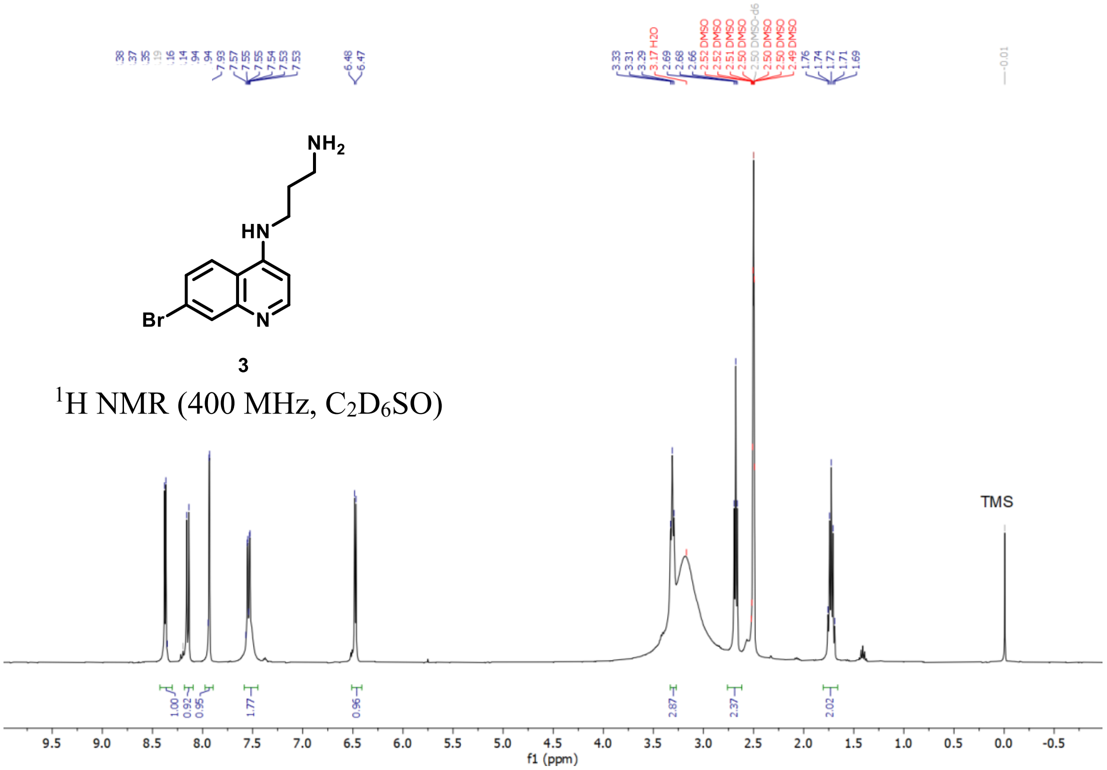

3
$^1$H NMR (400 MHz, $C_2D_6SO$)
TMS
f1 (ppm)

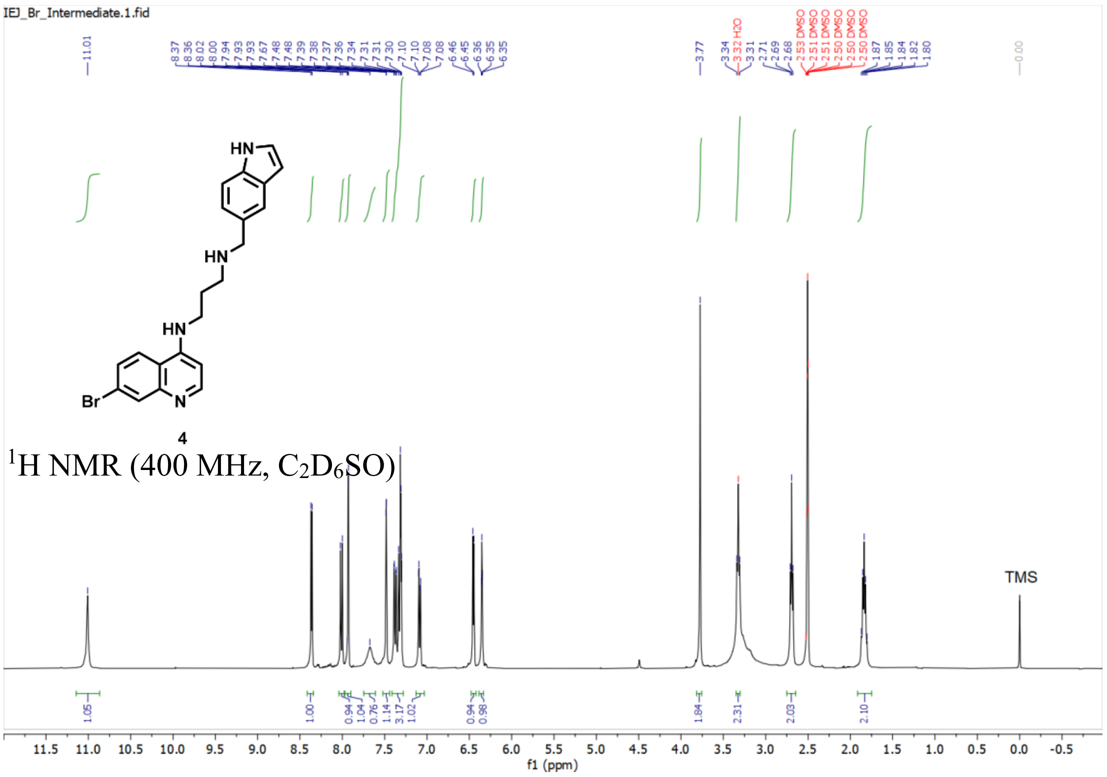

IEJ_Br_Intermediate.1.fid
4
$^1$H NMR (400 MHz, $C_2D_6SO$)
TMS
f1 (ppm)

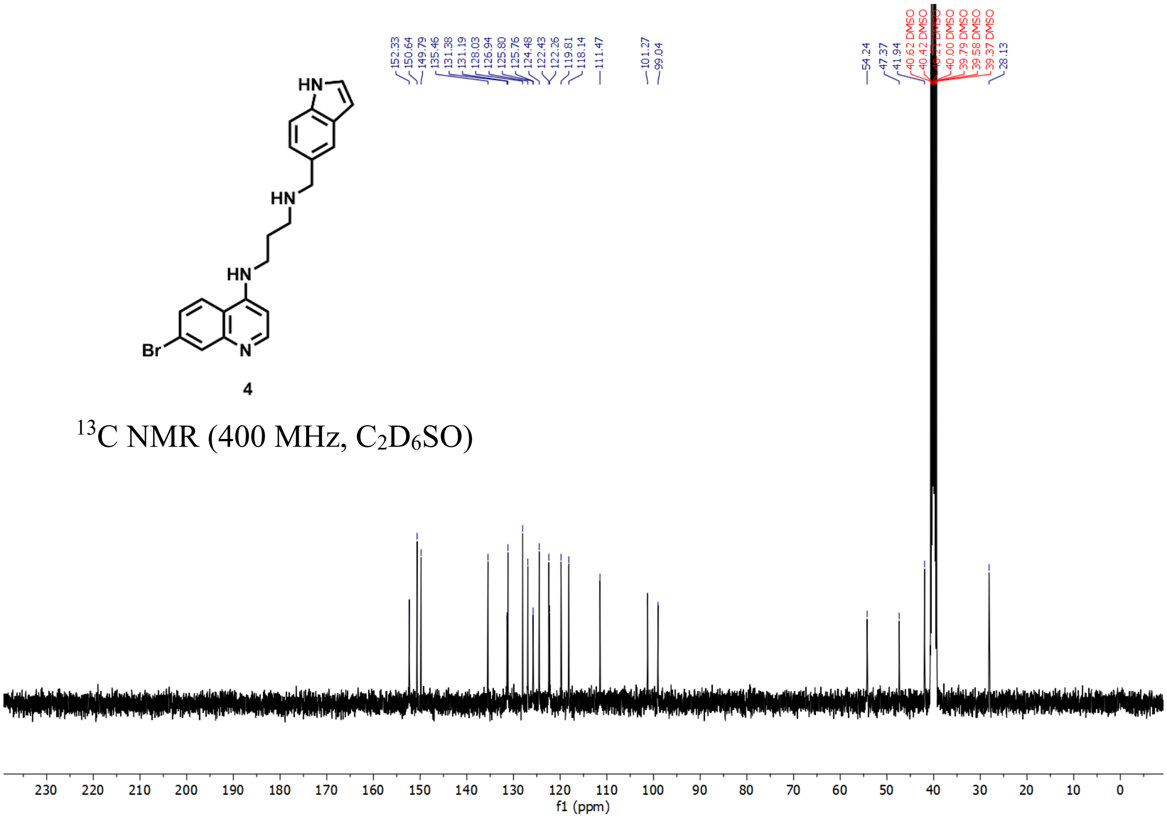

4

$^{13}C$ NMR (400 MHz, $C_2D_6SO$)

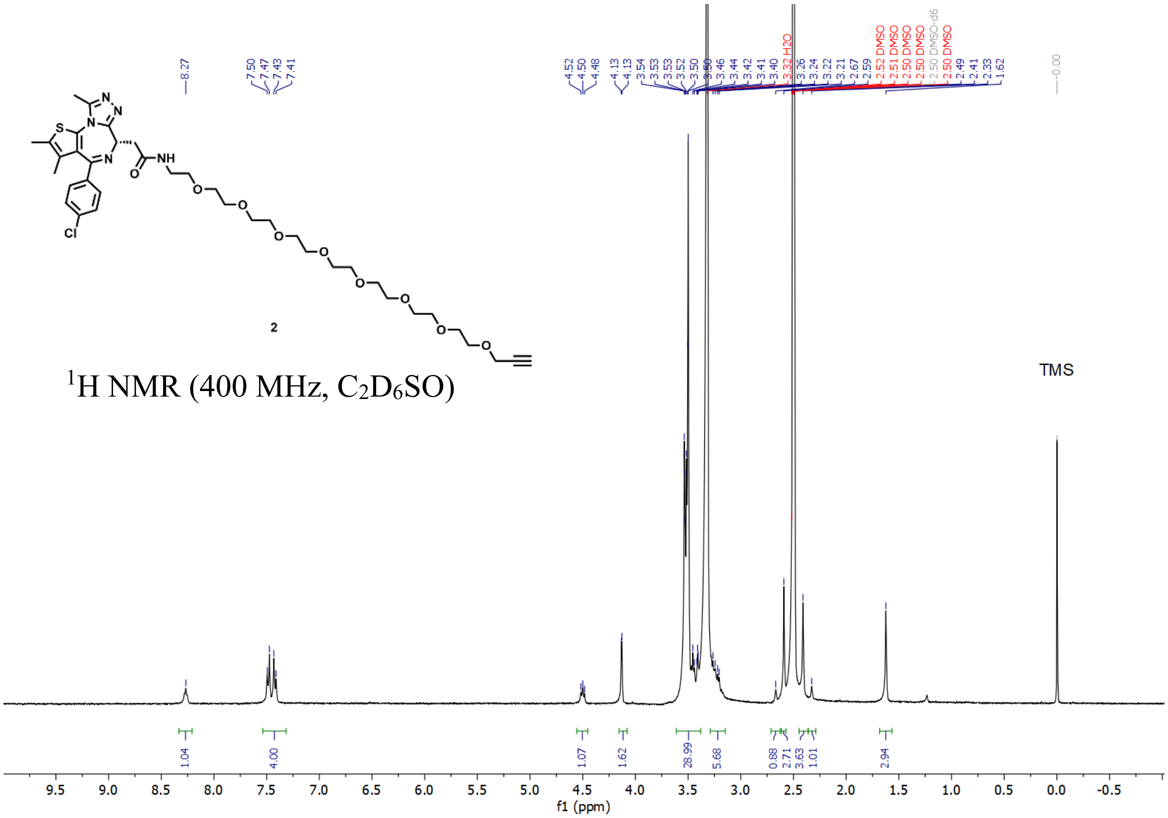

2

$^{1}H$ NMR (400 MHz, $C_2D_6SO$)

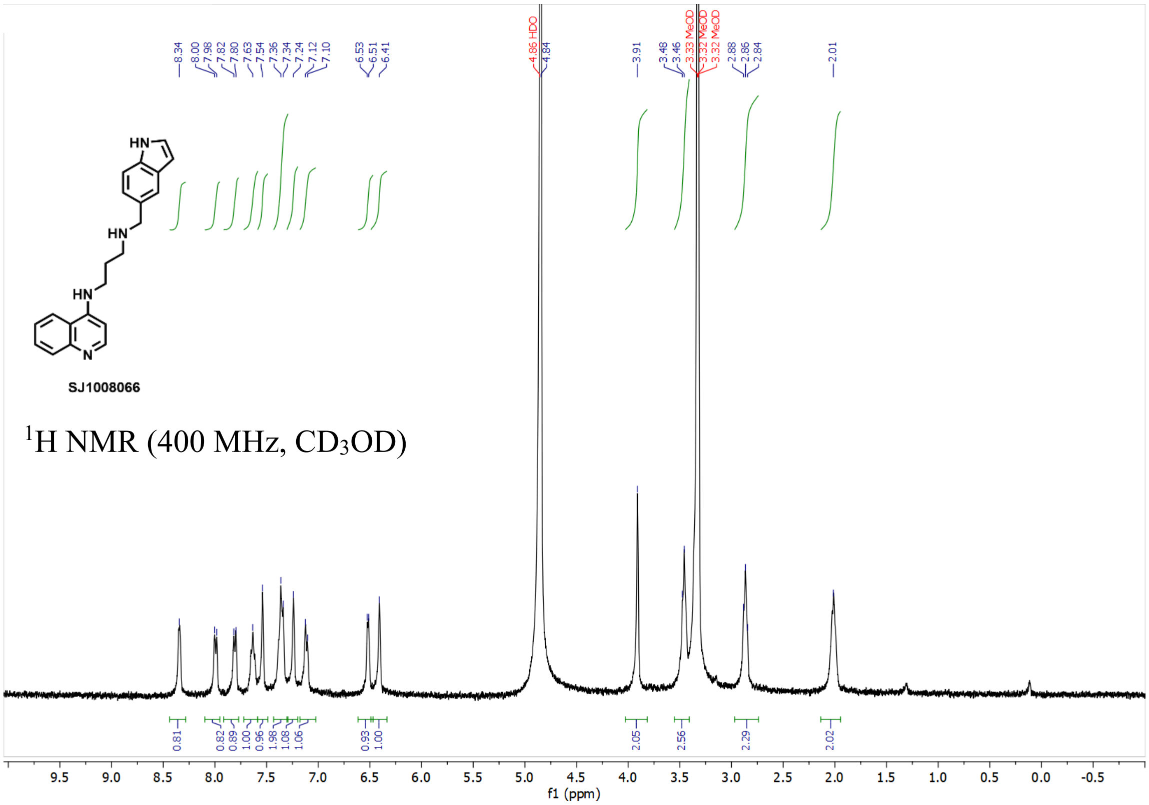

SJ1008066

$^{1}H$ NMR (400 MHz, $CD_3OD$)

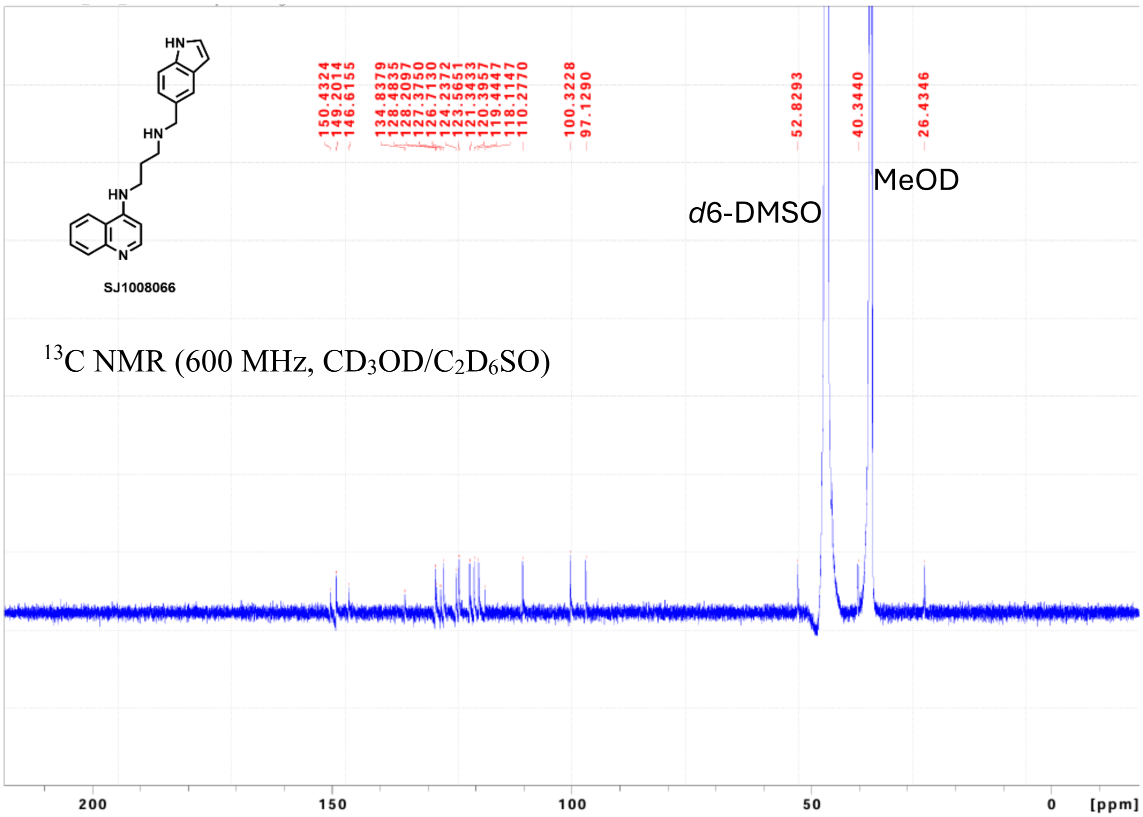

SJ1008066

$^{13}C$ NMR (600 MHz, $CD_3OD/C_2D_6SO$)

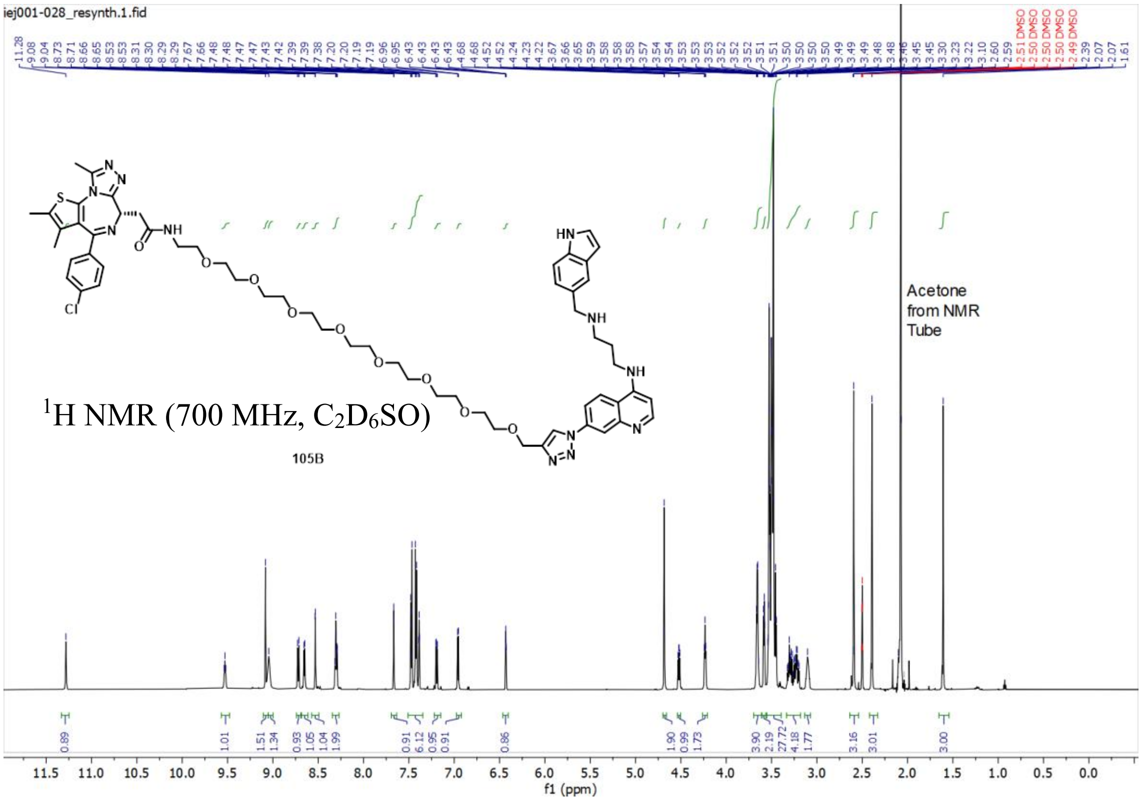
iej001-028_resynth.1.fid
Acetone from NMR Tube
1H NMR (700 MHz, C2D6SO)
105B
f1 (ppm)

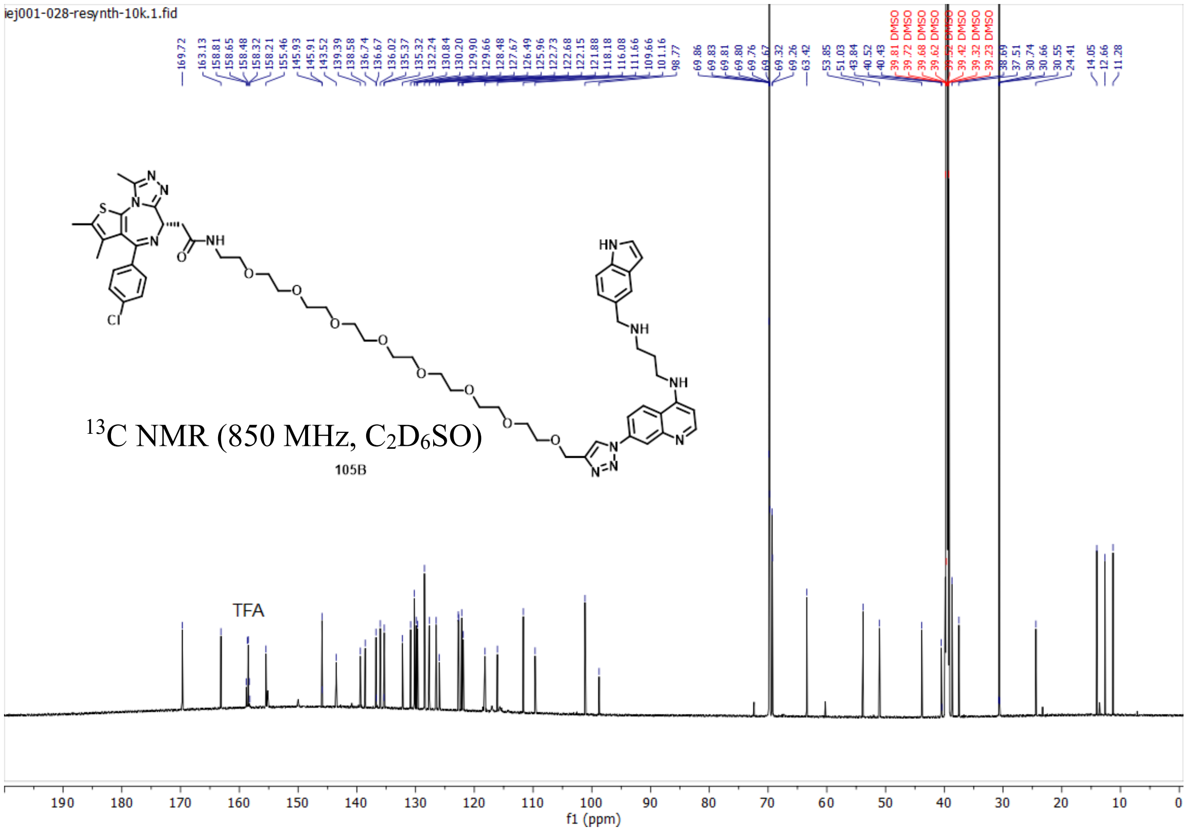
iej001-028-resynth-10k.1.fid
13C NMR (850 MHz, C2D6SO)
105B
TFA
f1 (ppm)

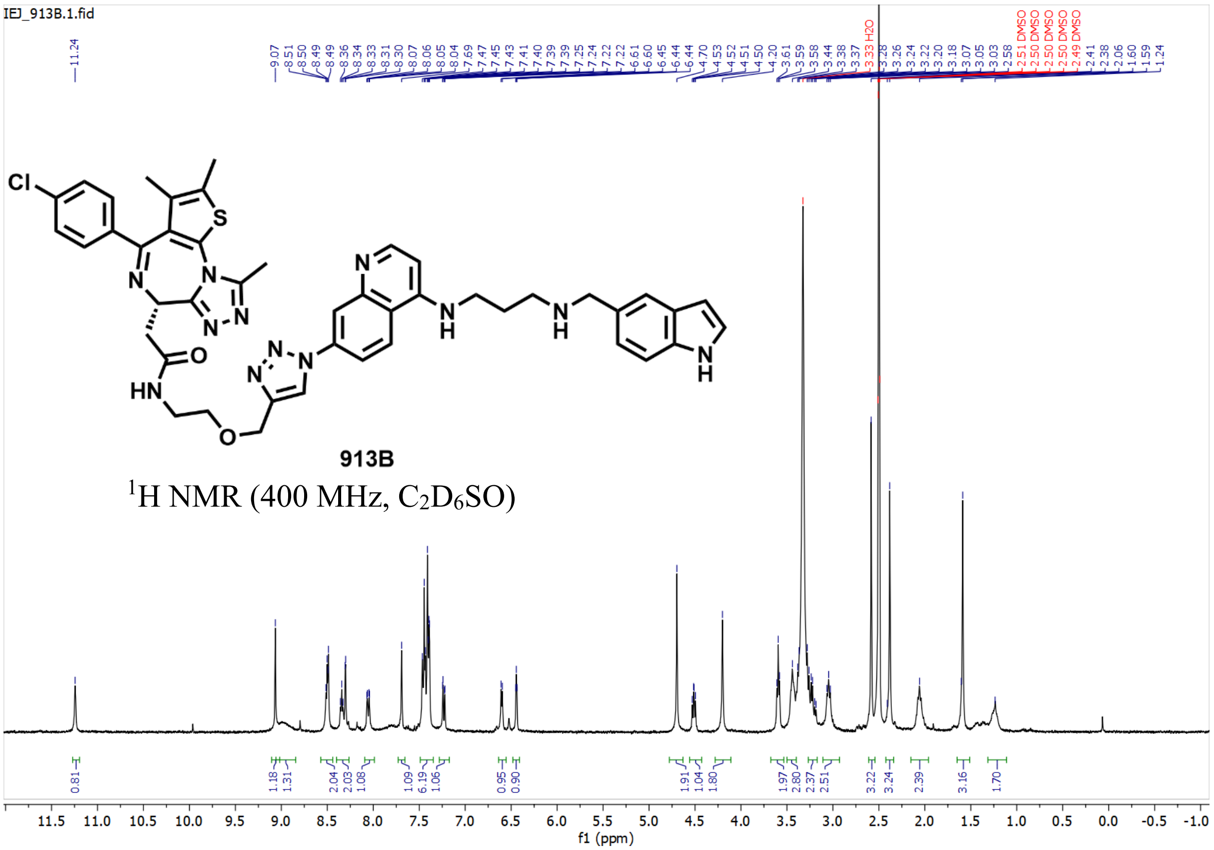
IEJ_913B.1.fid
913B
1H NMR (400 MHz, C2D6SO)
f1 (ppm)

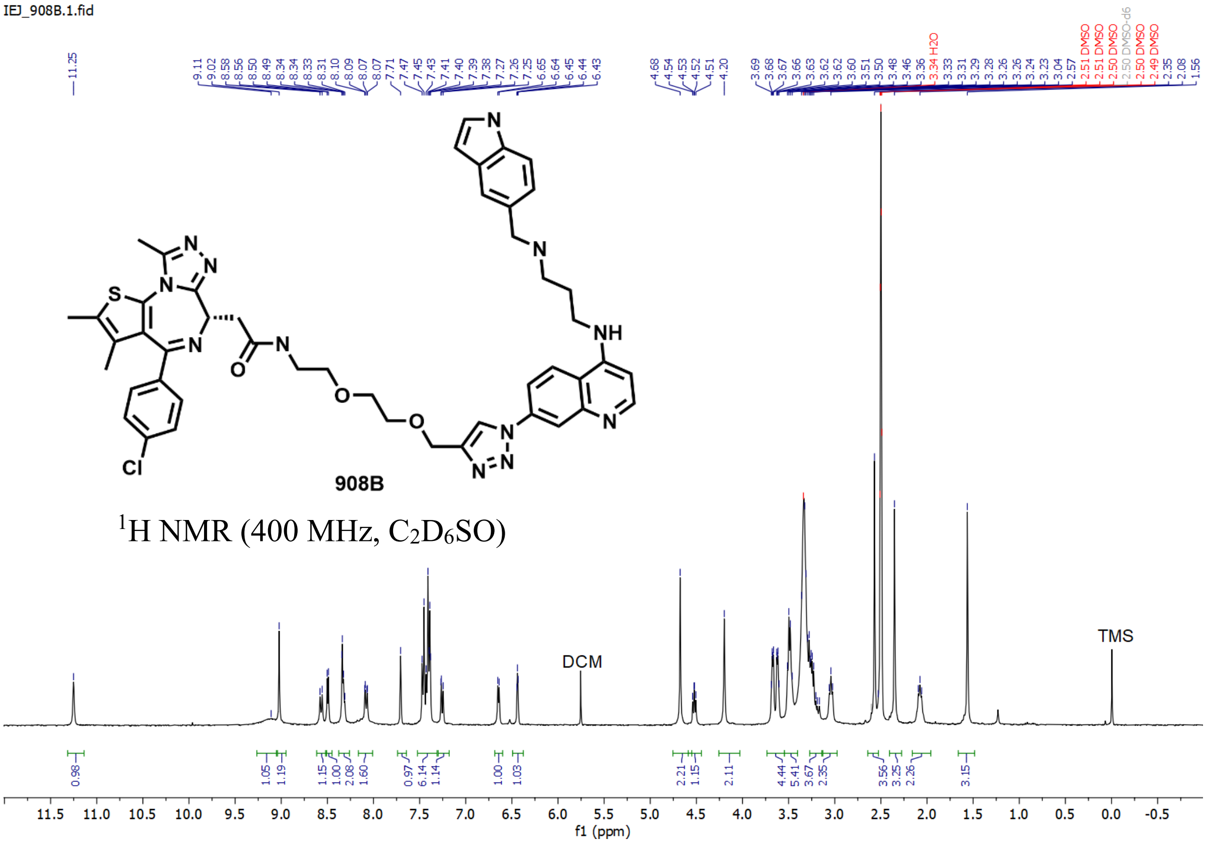
IEJ_908B.1.fid
908B
1H NMR (400 MHz, C2D6SO)
DCM
TMS
f1 (ppm)

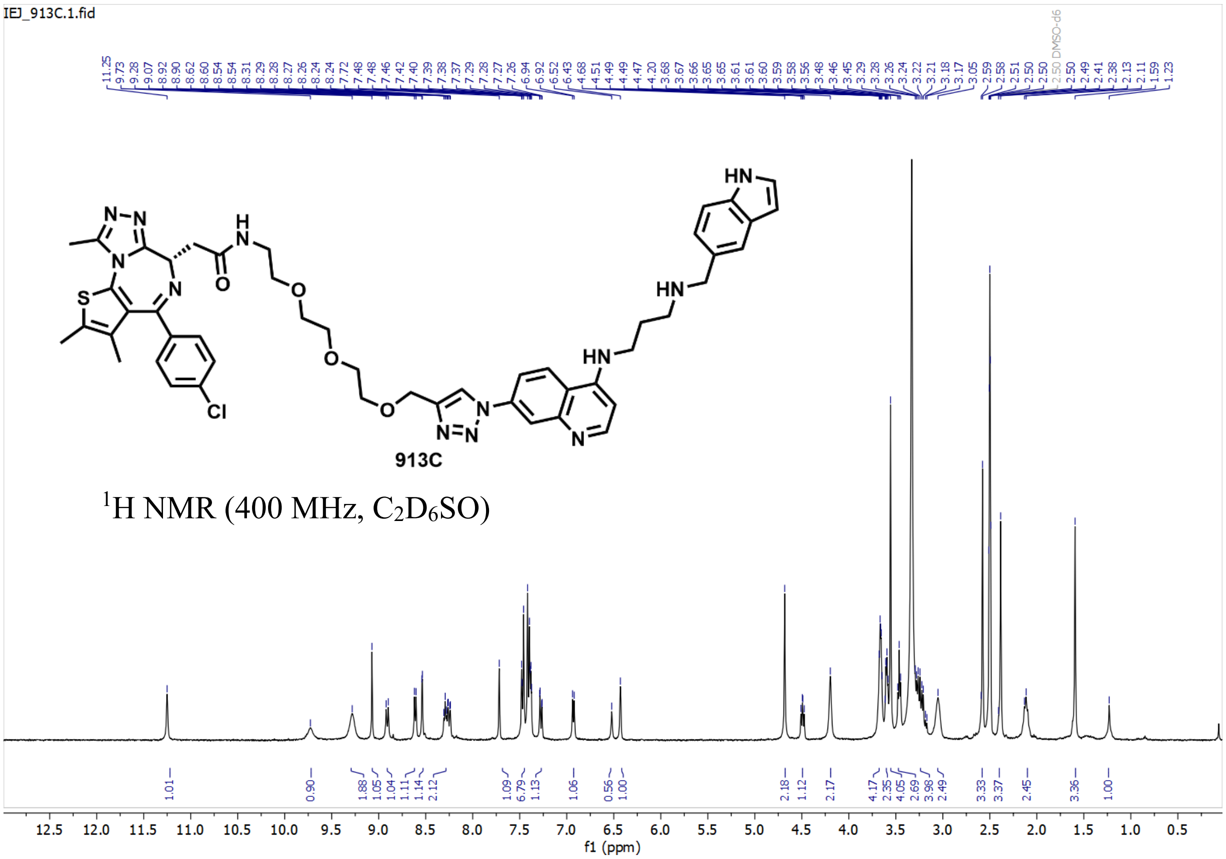
IEJ_913C.1.fid
913C
1H NMR (400 MHz, C2D6SO)
f1 (ppm)

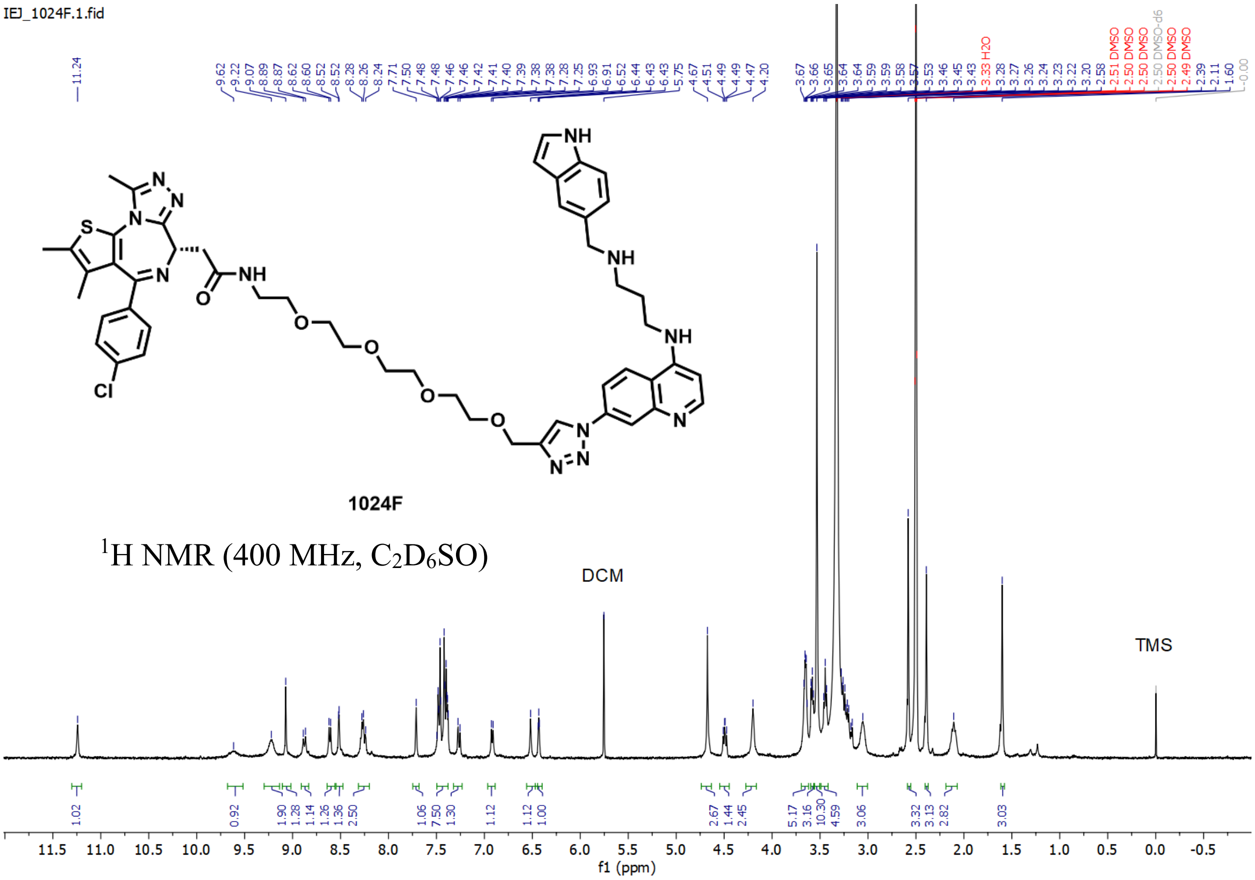
IEJ_1024F.1.fid
1024F
1H NMR (400 MHz, C2D6SO)
DCM
TMS
f1 (ppm)

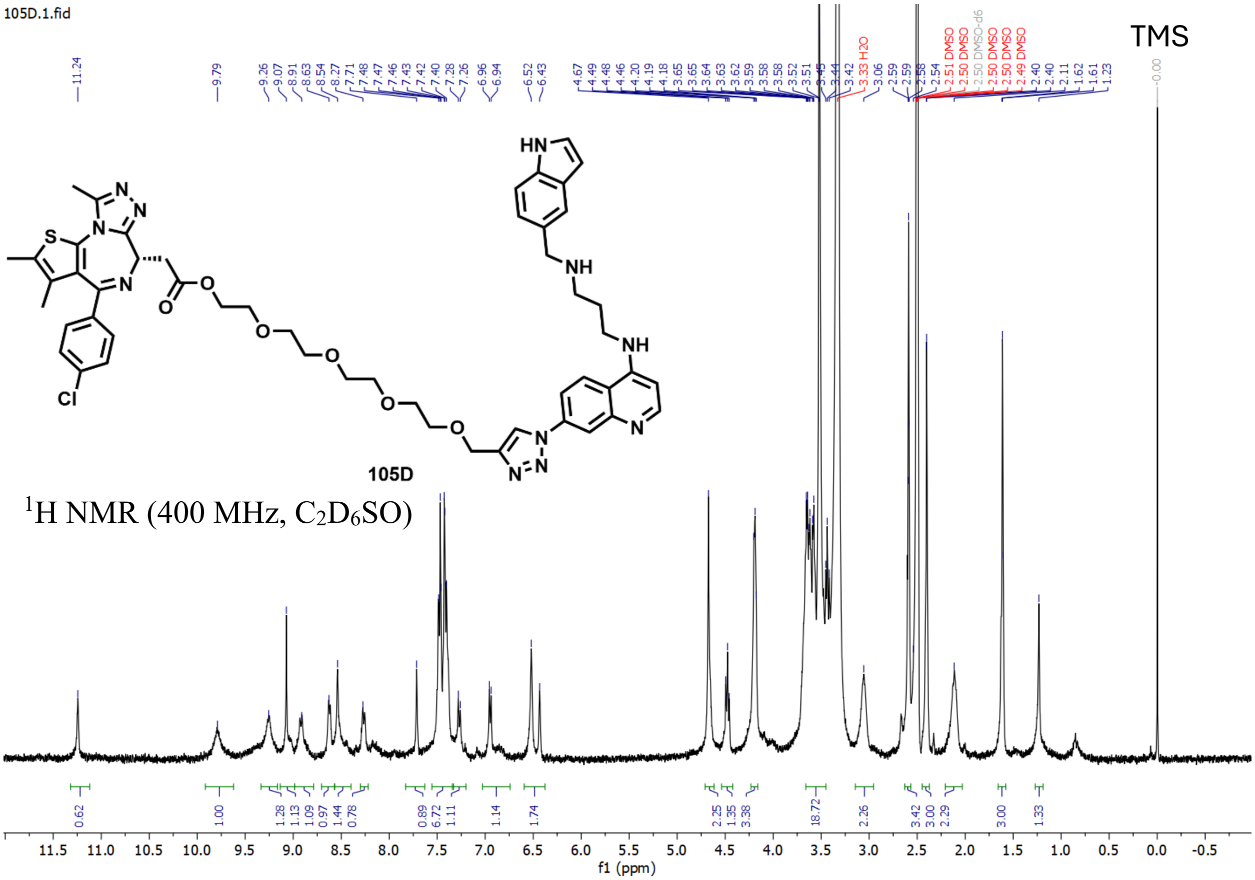
105D.1.fid
TMS
105D
$^{1}$H NMR (400 MHz, $C_2D_6SO$)
f1 (ppm)

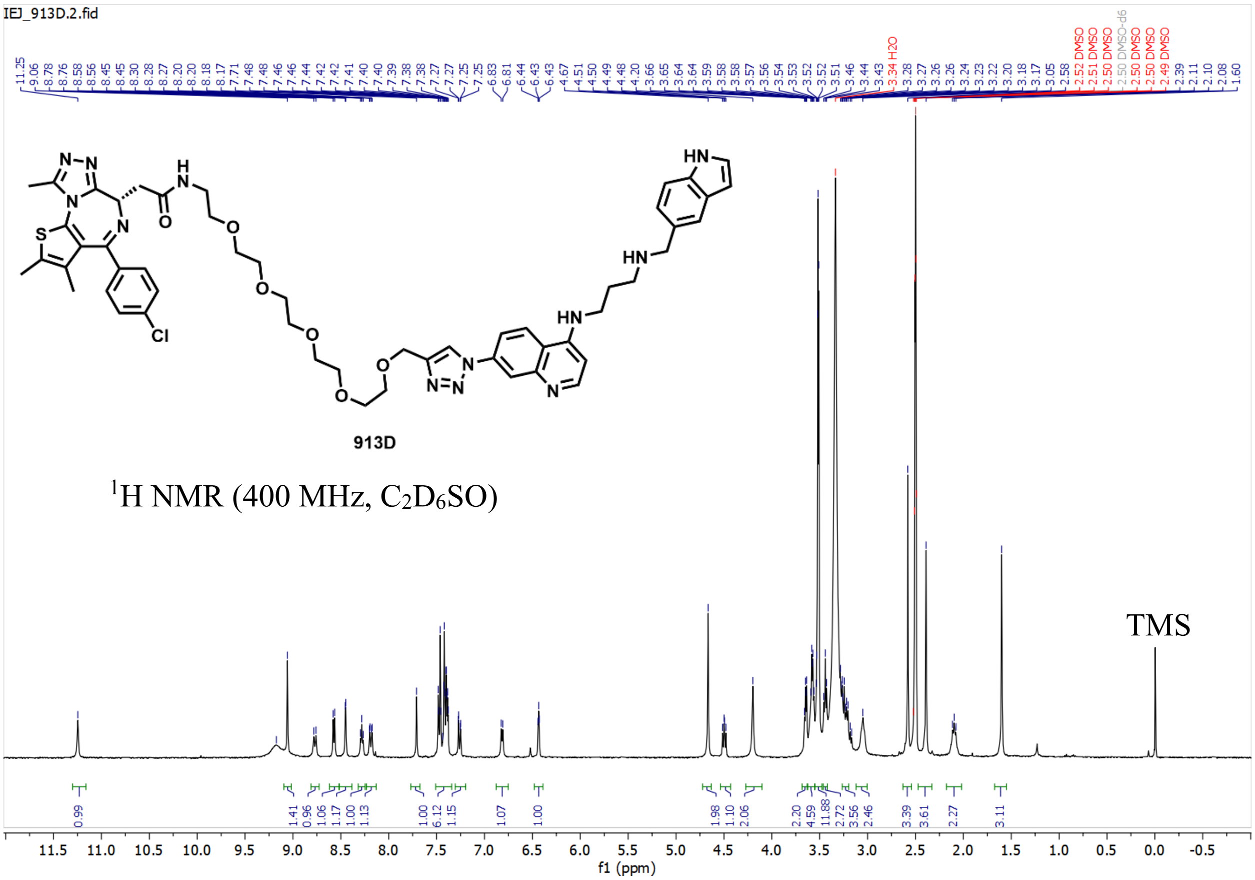
IEJ_913D.2.fid
913D
$^{1}$H NMR (400 MHz, $C_2D_6SO$)
TMS
f1 (ppm)

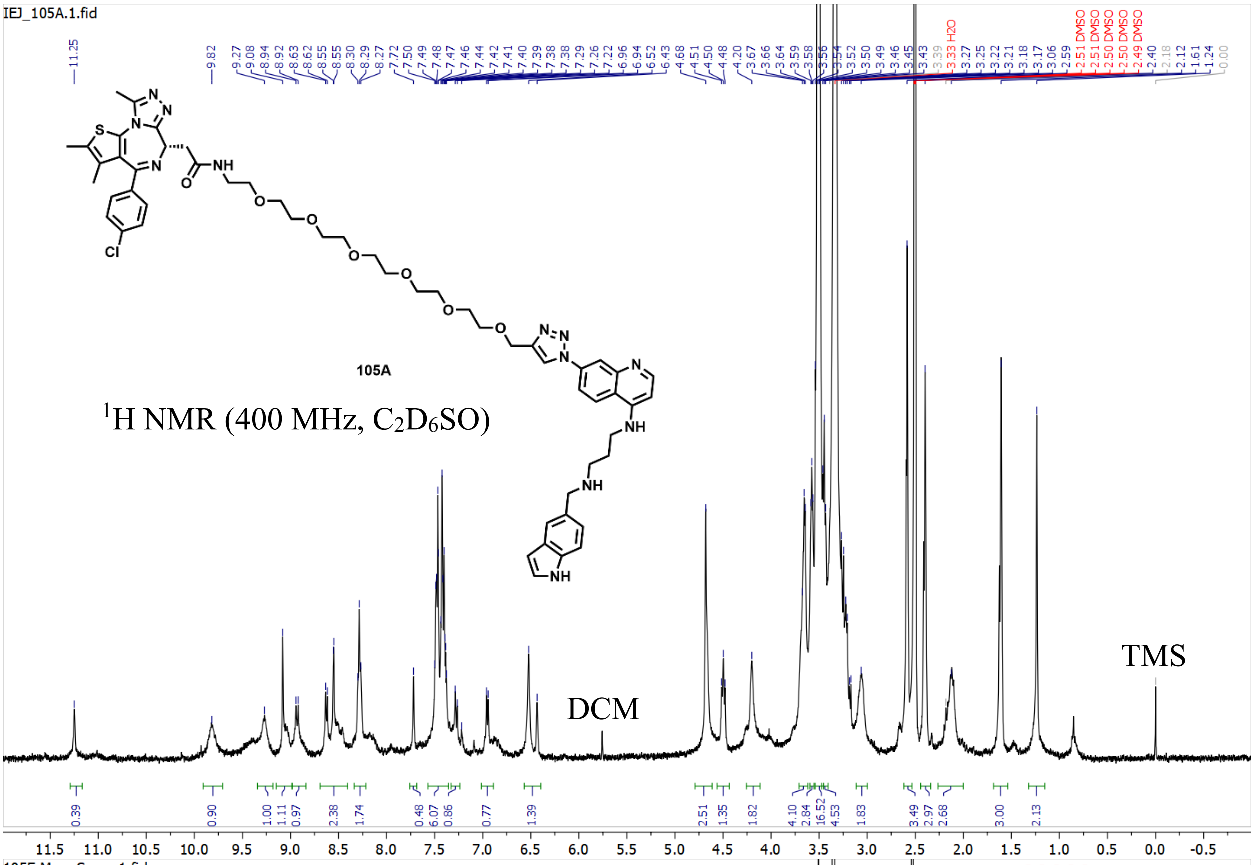
IEJ_105A.1.fid
105A
$^1$H NMR (400 MHz, $C_2D_6SO$)
DCM
TMS

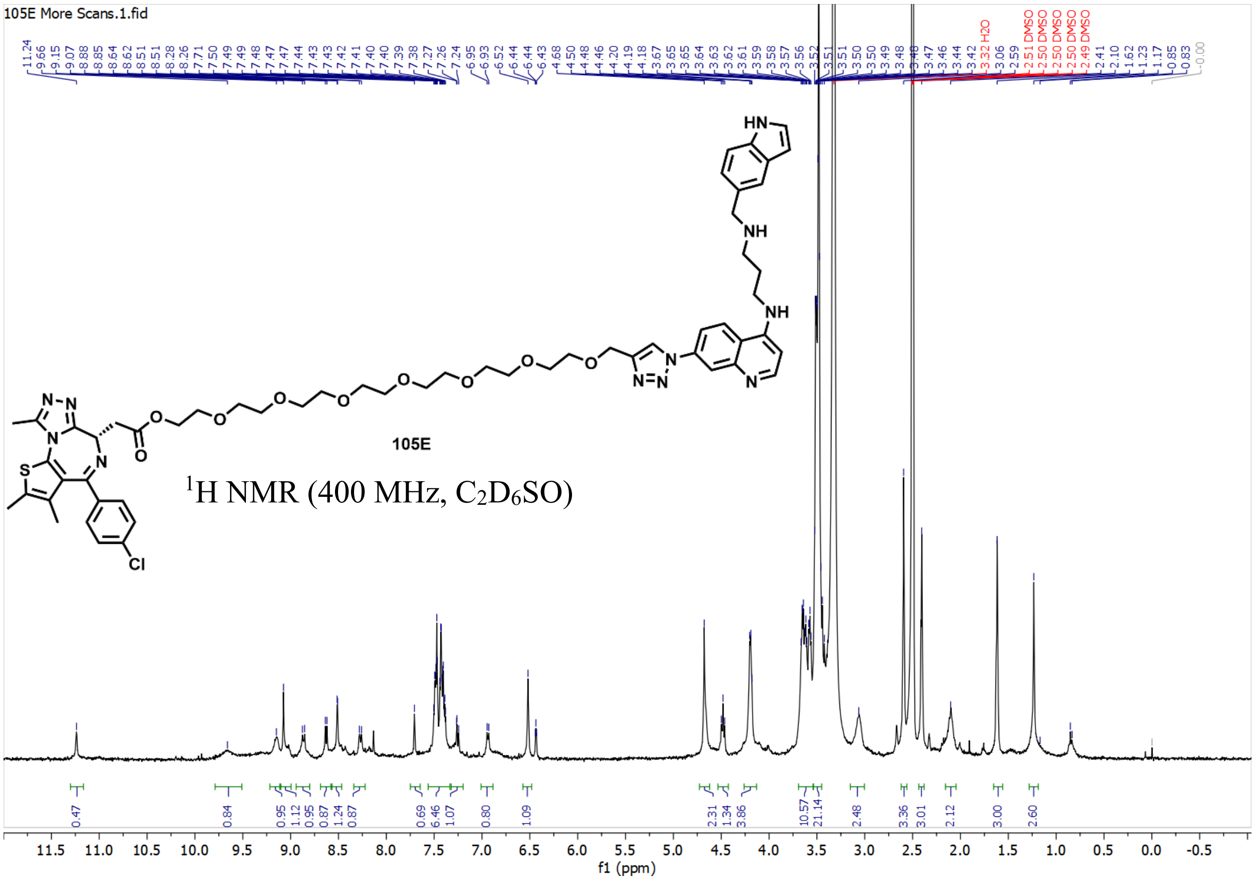
105E More Scans.1.fid
105E
$^1$H NMR (400 MHz, $C_2D_6SO$)
f1 (ppm)

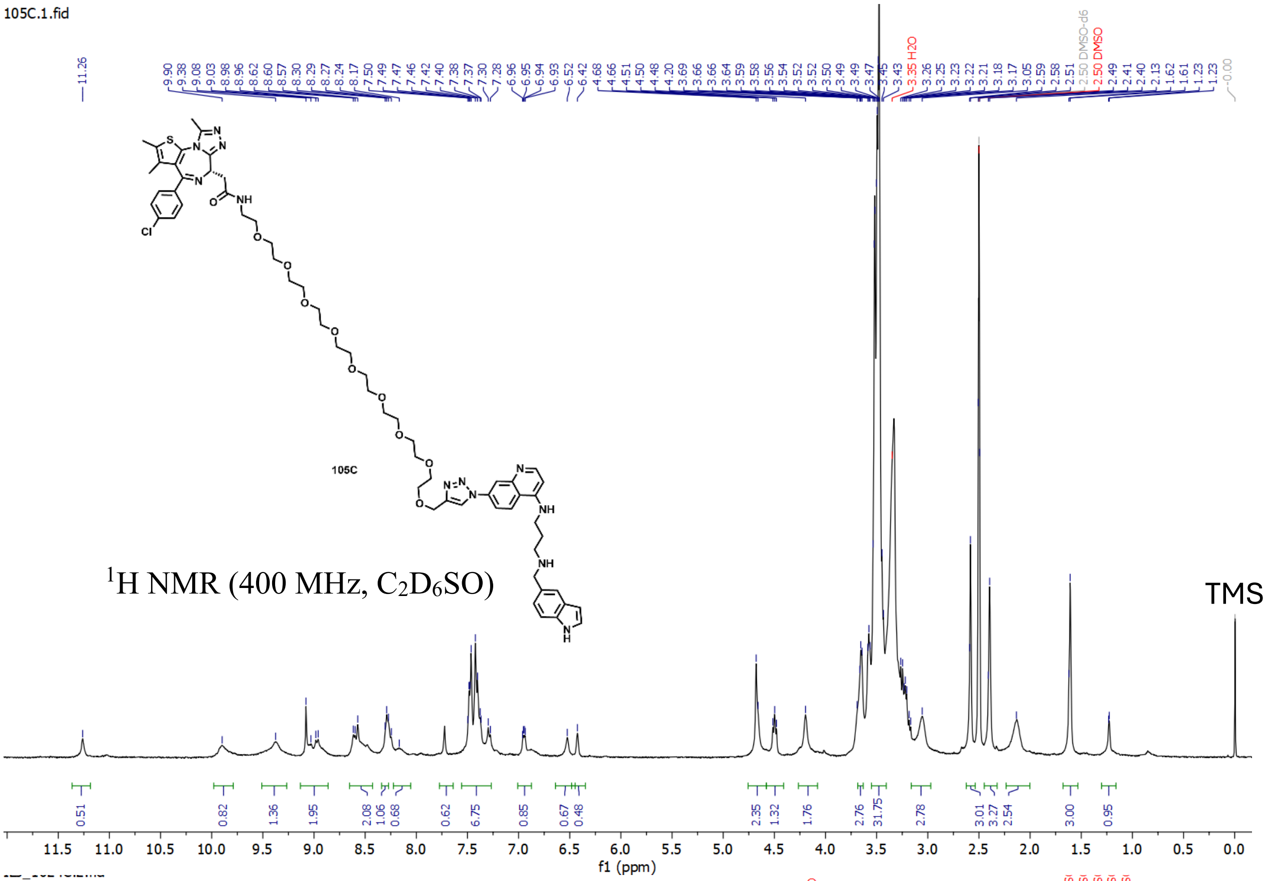
105C.1.fid
105C
¹H NMR (400 MHz, C2D6SO)
TMS
f1 (ppm)

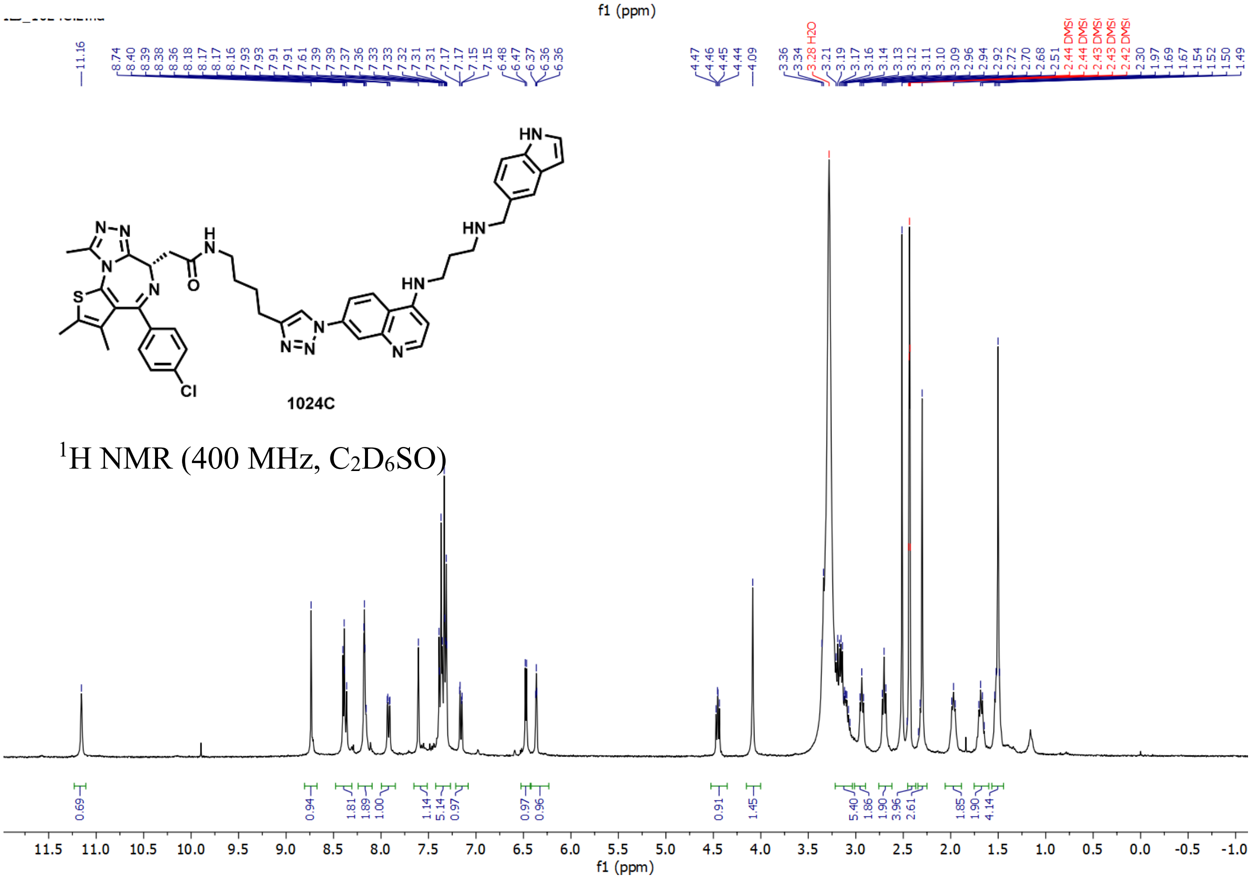
1024C
¹H NMR (400 MHz, C2D6SO)
f1 (ppm)

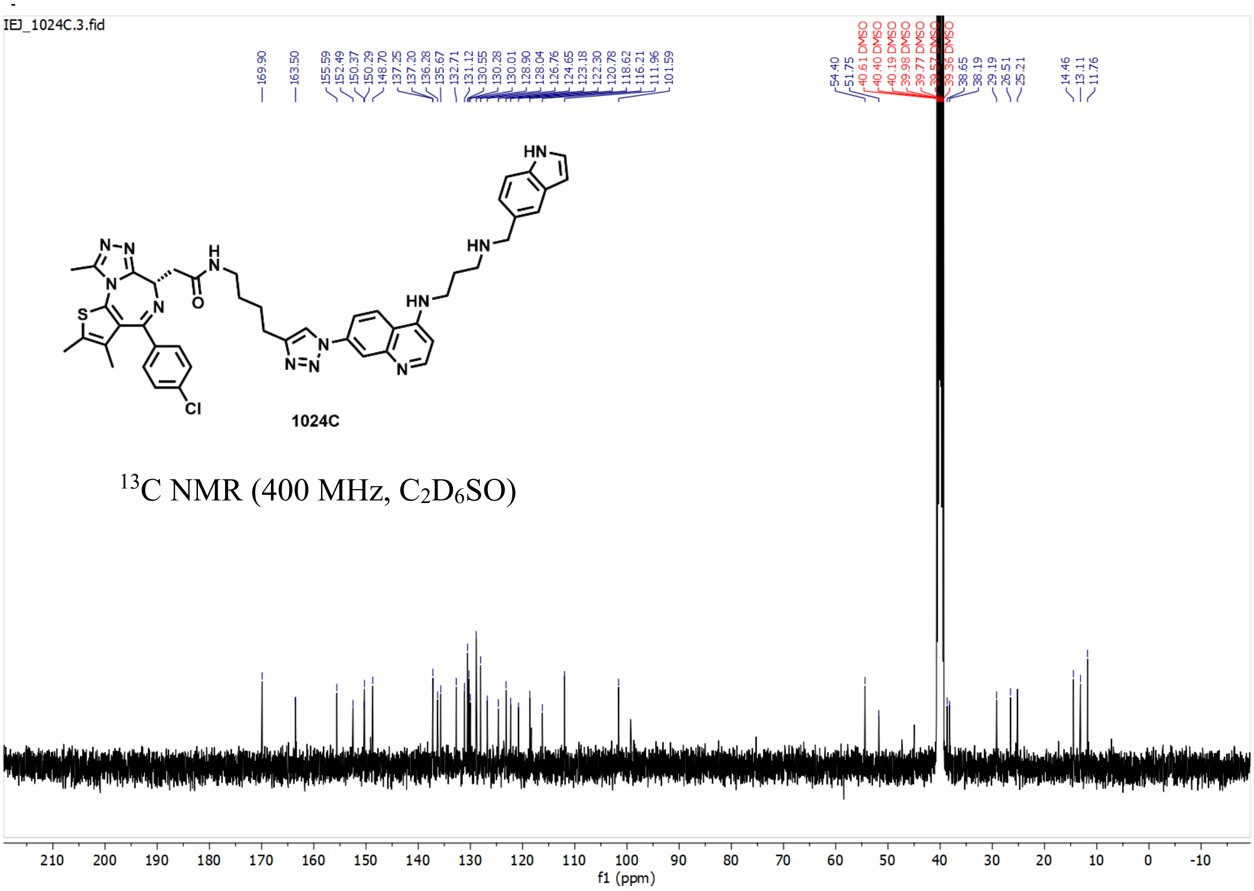

IEJ_1024C.3.fid
169.90
163.50
155.59
152.49
150.37
150.29
148.70
137.25
137.20
136.28
135.67
132.71
131.12
130.55
130.28
130.01
128.90
128.04
126.76
124.65
123.18
122.30
120.78
118.62
116.21
111.96
101.59
54.40
51.75
40.61 DMSO
40.40 DMSO
40.19 DMSO
39.98 DMSO
39.77 DMSO
39.57 DMSO
39.36 DMSO
38.65
38.19
29.19
26.51
25.21
14.46
13.11
11.76
1024C
$^{13}$C NMR (400 MHz, $C_2D_6SO$)
f1 (ppm)

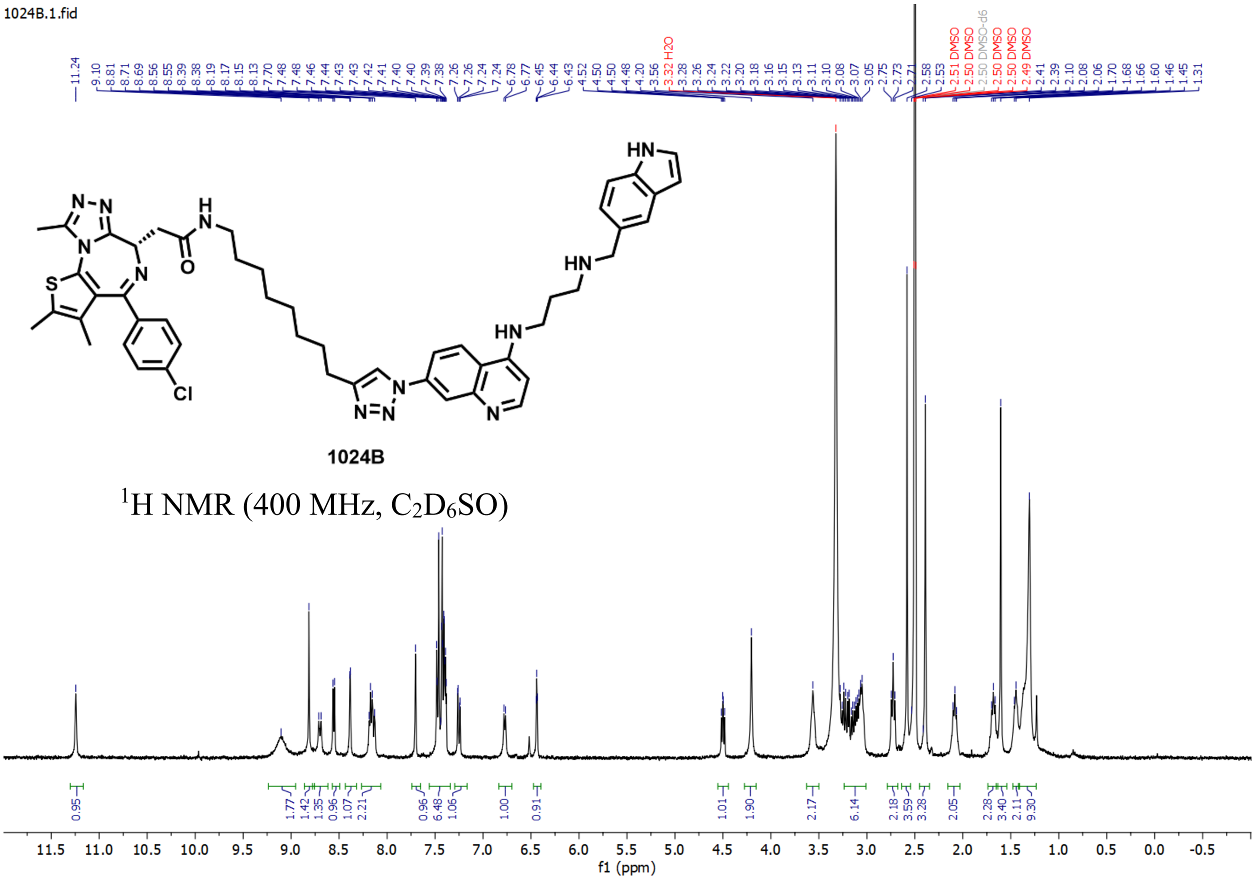

1024B.1.fid
11.24
9.10
8.81
8.71
8.69
8.56
8.55
8.39
8.38
8.19
8.17
8.15
8.13
7.70
7.48
7.46
7.44
7.43
7.42
7.41
7.40
7.39
7.38
7.26
7.26
7.24
6.78
6.77
6.45
6.44
6.43
4.52
4.50
4.50
4.48
4.20
3.56
3.32 H2O
3.28
3.26
3.24
3.22
3.20
3.18
3.16
3.15
3.13
3.11
3.10
3.08
3.07
3.05
2.75
2.73
2.71
2.58
2.53
2.51 DMSO
2.50 DMSO
2.50 DMSO-d6
2.50 DMSO
2.50 DMSO
2.49 DMSO
2.41
2.39
2.10
2.08
2.06
1.70
1.68
1.66
1.60
1.46
1.45
1.31
1024B
$^{1}$H NMR (400 MHz, $C_2D_6SO$)
f1 (ppm)

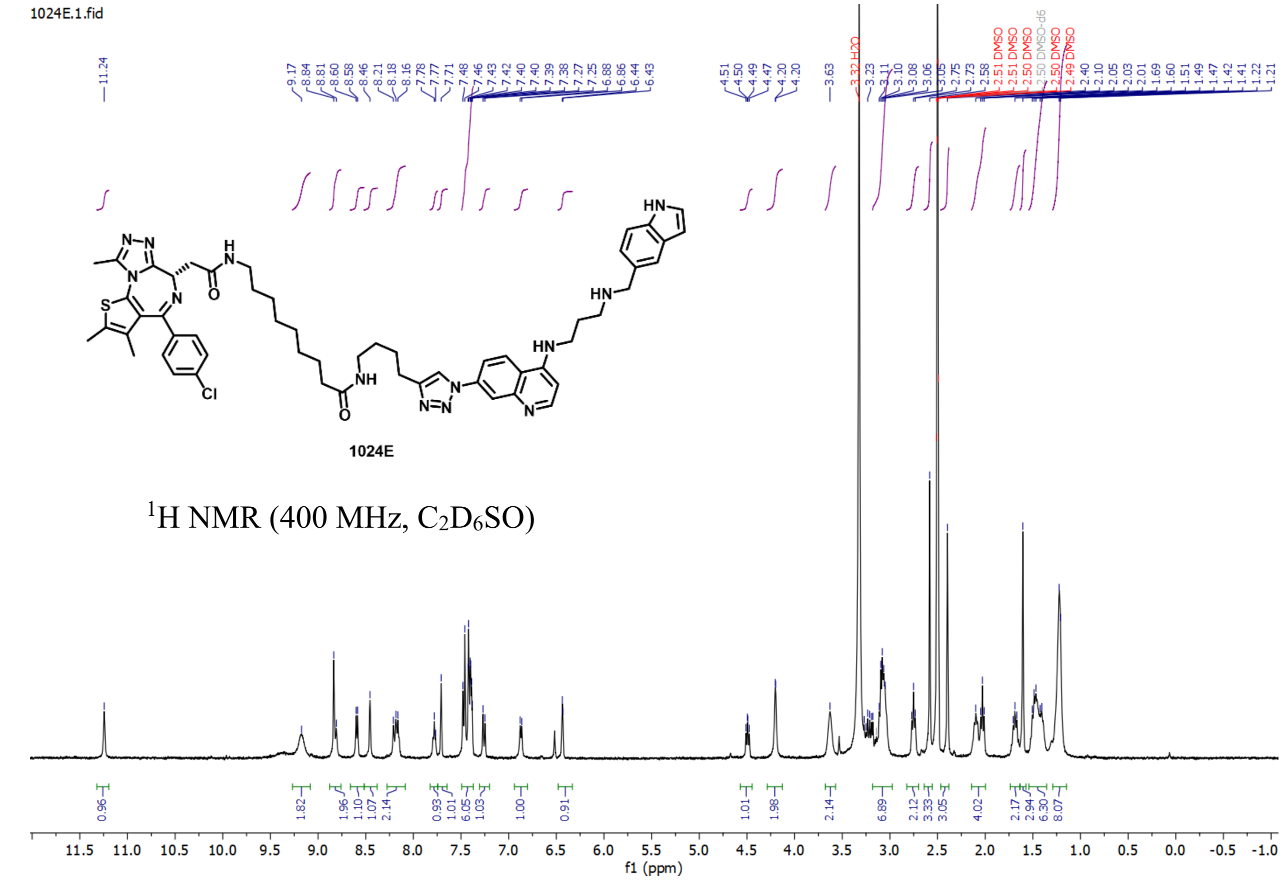
1024E.1.fid
1024E
$^{1}$H NMR (400 MHz, $C_2D_6SO$)
f1 (ppm)

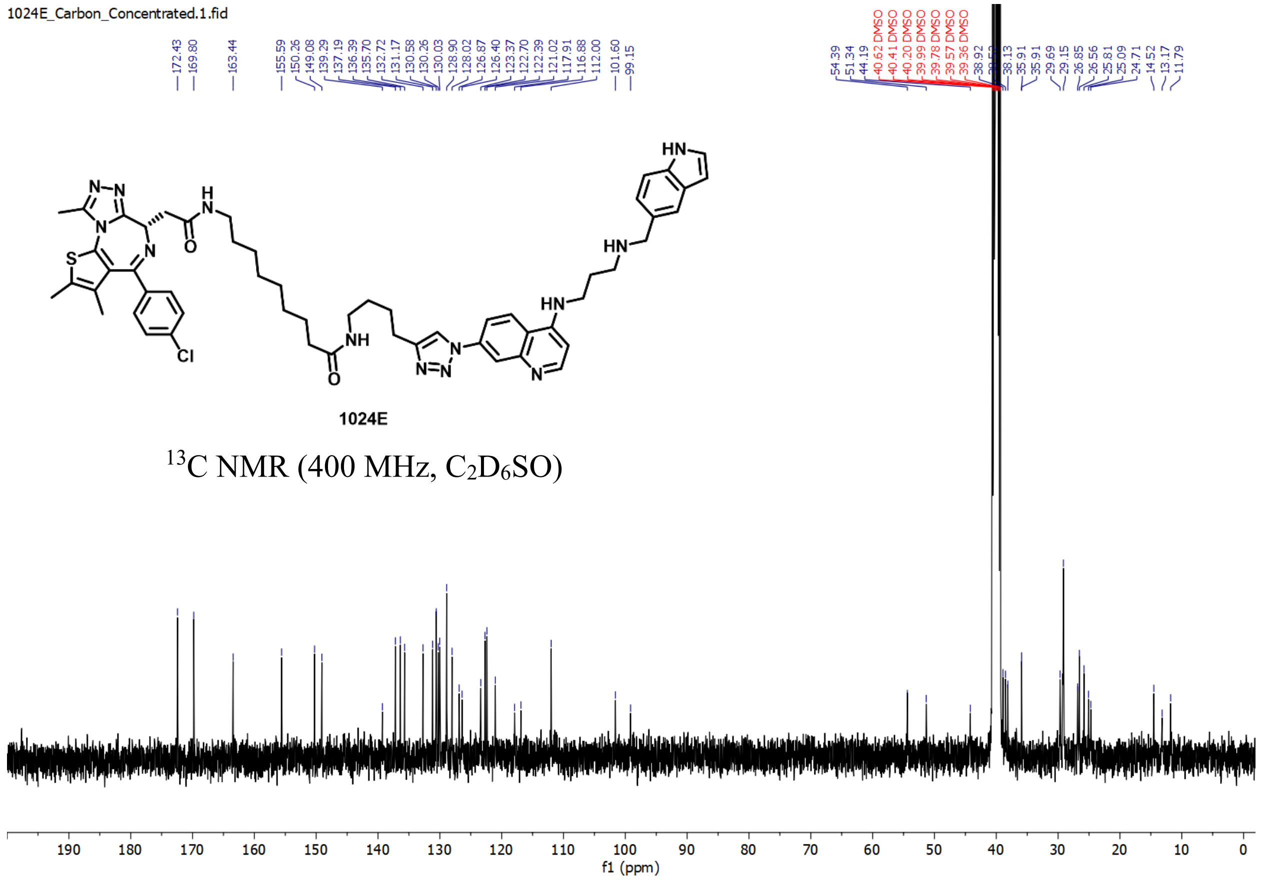
1024E_Carbon_Concentrated.1.fid
1024E
$^{13}$C NMR (400 MHz, $C_2D_6SO$)
f1 (ppm)

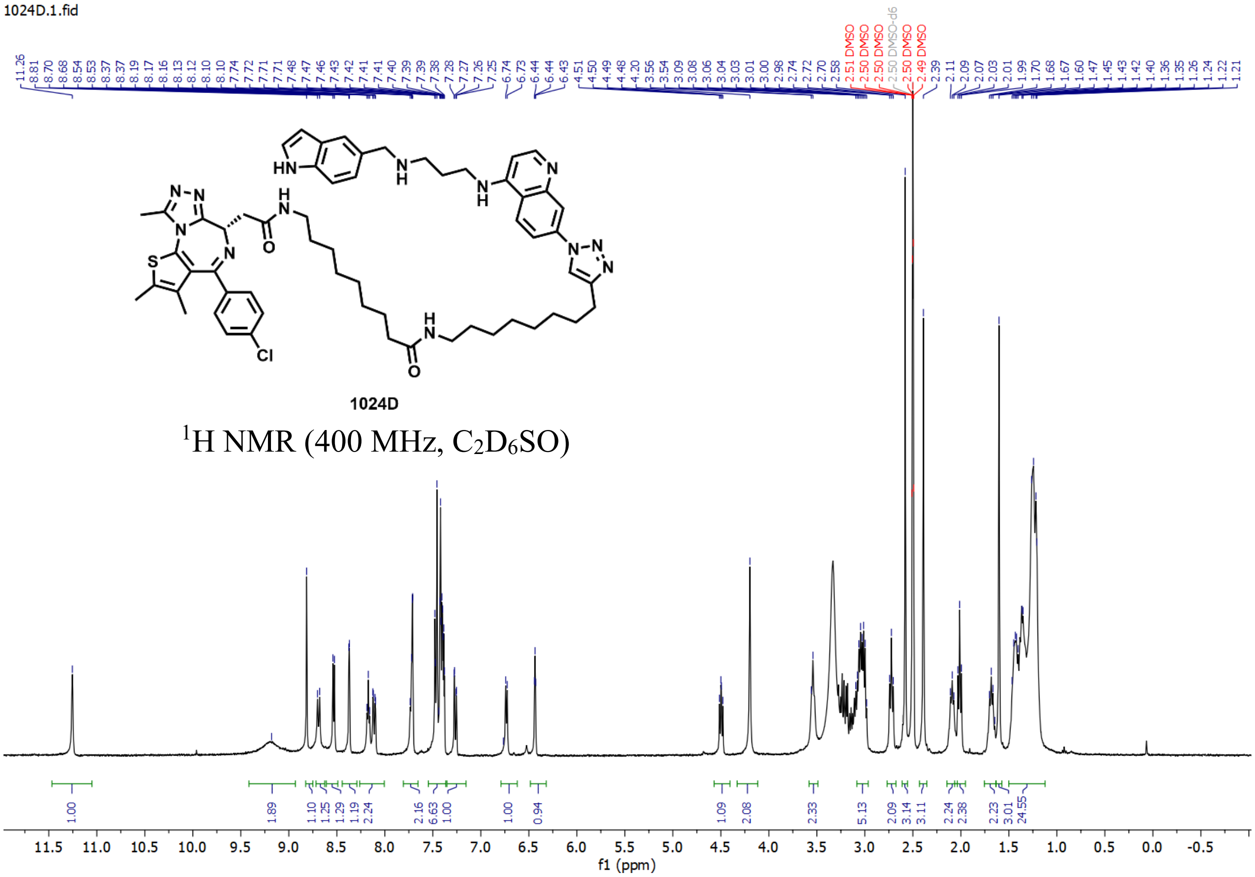

$^1$H NMR (400 MHz, $C_2D_6SO$)

### HPLC

**105B**

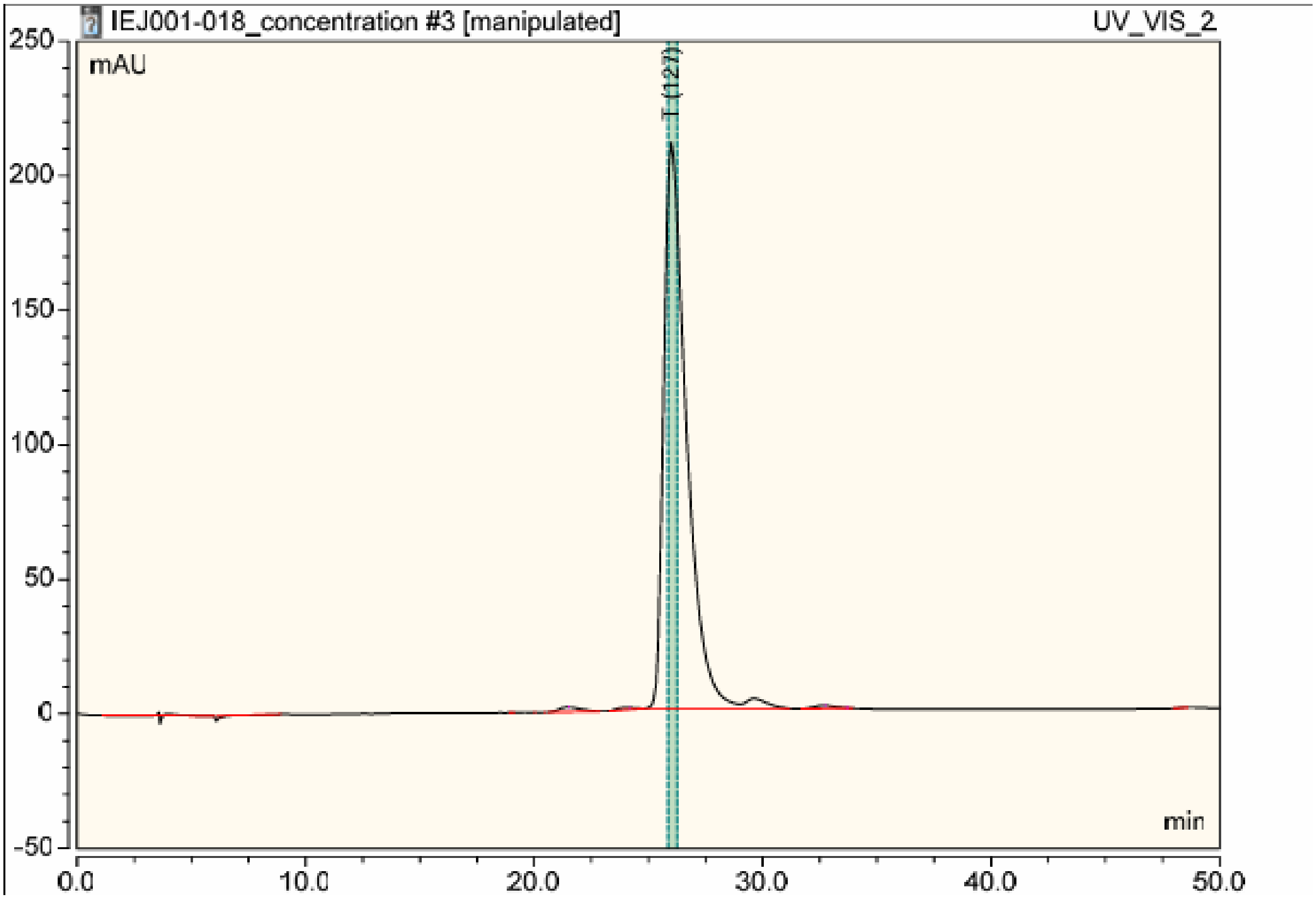

| No. | Ret.Time min | Peak Name | Height mAU | Area mAU*min | Rel.Area % | Amount n.a. | Type |
|---|---|---|---|---|---|---|---|
| 1 | 3.39 | | 0.056 | 0.182 | 0.07 | n.a. | BMb* |
| 2 | 3.58 | | 1.210 | 0.057 | 0.02 | n.a. | bM * |
| 3 | 3.90 | | 0.456 | 0.040 | 0.02 | n.a. | MB* |
| 4 | 8.91 | | 0.000 | 0.299 | 0.12 | n.a. | MB* |
| 5 | 19.06 | | 0.031 | 0.005 | 0.00 | n.a. | BMB |
| 6 | 20.00 | | 0.019 | 0.001 | 0.00 | n.a. | BMB |
| 7 | 20.50 | | 0.013 | 0.001 | 0.00 | n.a. | BMB |
| 8 | 21.44 | | 1.739 | 1.881 | 0.75 | n.a. | BMB |
| 9 | 21.48 | | 0.010 | 0.000 | 0.00 | n.a. | Rd |
| 10 | 24.15 | | 0.738 | 0.748 | 0.30 | n.a. | BMb |
| 11 | 24.18 | | 0.008 | 0.000 | 0.00 | n.a. | Rd |
| 12 | 26.00 | | 210.476 | 241.666 | 96.52 | n.a. | bM |
| 13 | 29.64 | | 3.808 | 4.142 | 1.65 | n.a. | MB |
| 14 | 32.64 | | 1.181 | 1.329 | 0.53 | n.a. | BMB |
| 15 | 32.71 | | 0.011 | 0.000 | 0.00 | n.a. | Rd |
| 16 | 33.69 | | 0.014 | 0.001 | 0.00 | n.a. | Rd |
| 17 | 48.59 | | 0.035 | 0.030 | 0.01 | n.a. | BMB |
| Total: | | | 219.805 | 250.383 | 100.00 | 0.000 | |

**SJ1008066**

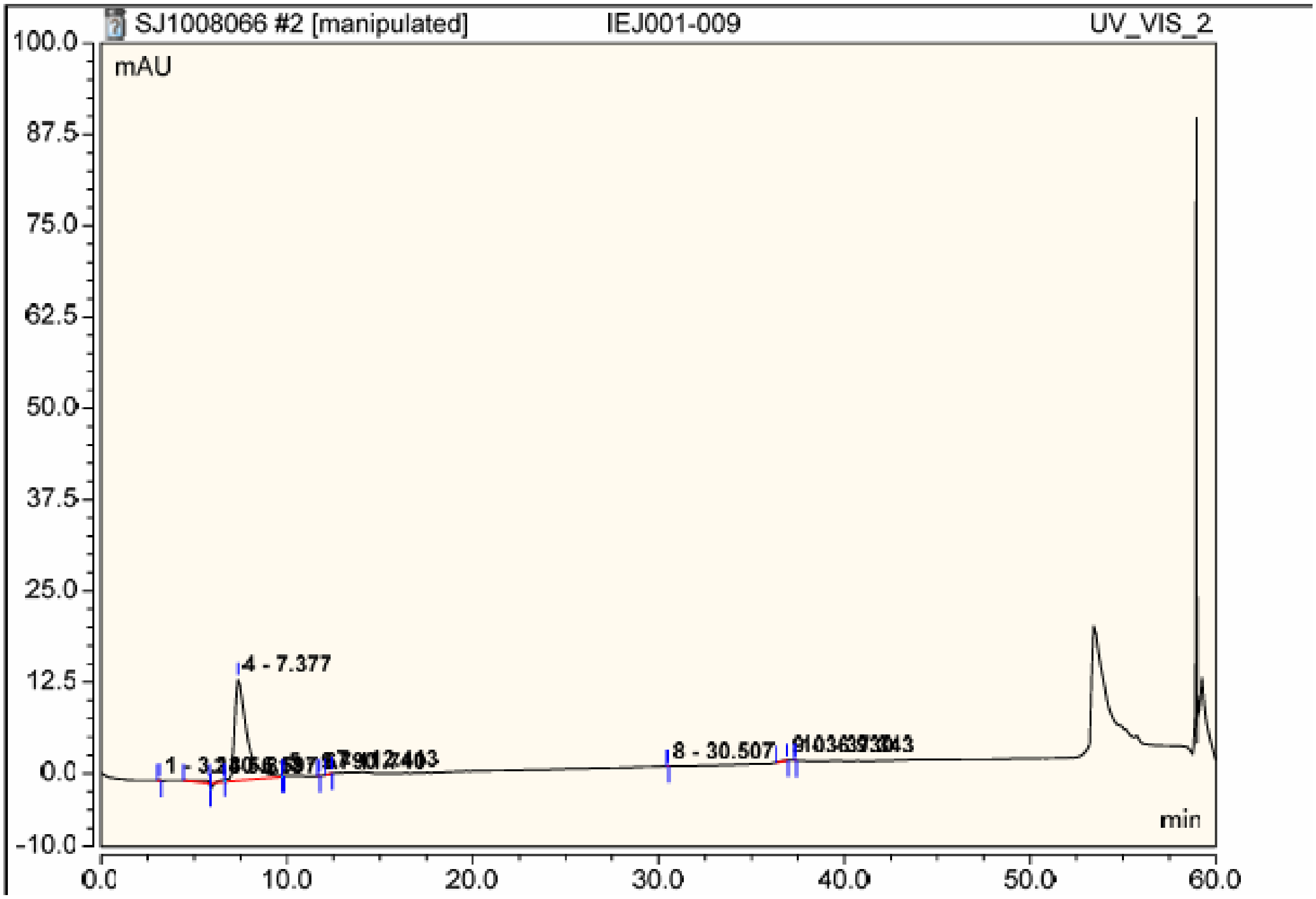

| No. | Ret.Time min | Peak Name | Height mAU | Area mAU*min | Rel.Area % | Amount n.a. | Type |
|---|---|---|---|---|---|---|---|
| 1 | 3.14 | | 0.093 | 0.010 | 0.09 | n.a. | BMB |
| 2 | 5.81 | | 0.417 | 0.227 | 2.05 | n.a. | BM * |
| 3 | 6.60 | | 0.259 | 0.009 | 0.08 | n.a. | M * |
| 4 | 7.38 | | 13.916 | 10.771 | 97.39 | n.a. | M |
| 5 | 9.79 | | 0.107 | 0.011 | 0.10 | n.a. | M * |
| 6 | 11.74 | | 0.015 | 0.001 | 0.01 | n.a. | BMB |
| 7 | 12.41 | | 0.013 | 0.006 | 0.06 | n.a. | BMB |
| 8 | 30.51 | | 0.017 | 0.001 | 0.01 | n.a. | BMB |
| 9 | 36.93 | | 0.023 | 0.023 | 0.21 | n.a. | BMB |
| 10 | 37.34 | | 0.018 | 0.001 | 0.01 | n.a. | BMB |
| Total: | | | 14.879 | 11.060 | 100.00 | 0.000 | |

**908B**

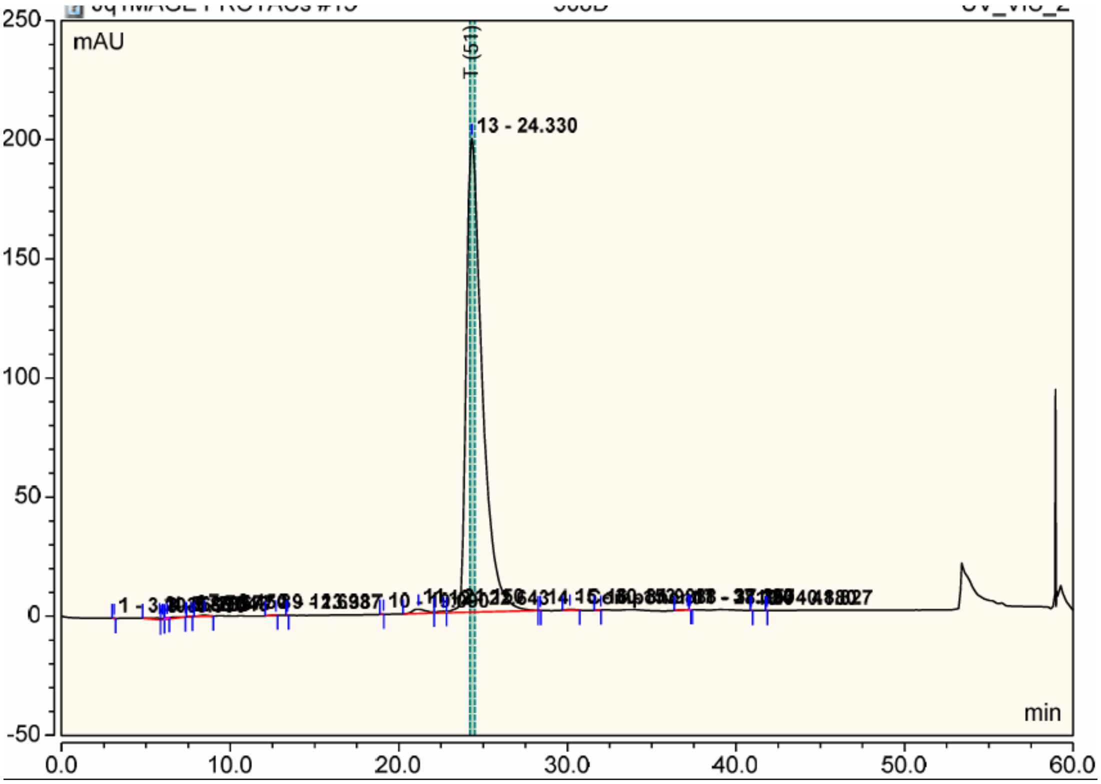

| No. | Ret.Time min | Peak Name | Height mAU | Area mAU*min | Rel.Area % | Amount micromol | Type |
|---|---|---|---|---|---|---|---|
| 1 | 3.10 | | 0.091 | 0.010 | 0.00 | n.a. | BMB |
| 2 | 5.81 | | 0.950 | 0.409 | 0.18 | n.a. | BMB |
| 3 | 5.99 | | 0.567 | 0.098 | 0.04 | n.a. | BM |
| 4 | 6.23 | | 0.197 | 0.036 | 0.02 | n.a. | Ru |
| 5 | 7.34 | | 0.000 | 0.429 | 0.19 | n.a. | MB |
| 6 | 7.68 | | 0.046 | 0.018 | 0.01 | n.a. | BMB |
| 7 | 8.45 | | 0.457 | 0.236 | 0.11 | n.a. | BMB |
| 8 | 12.69 | | 0.083 | 0.046 | 0.02 | n.a. | BMB |
| 9 | 13.39 | | 0.017 | 0.001 | 0.00 | n.a. | BMB |
| 10 | 19.09 | | 0.010 | 0.003 | 0.00 | n.a. | BMB |
| 11 | 21.15 | | 1.809 | 1.703 | 0.77 | n.a. | BM |
| 12 | 22.64 | | 0.649 | 0.406 | 0.18 | n.a. | M |
| 13 | 24.33 | | 198.867 | 217.530 | 98.29 | n.a. | M |
| 14 | 28.39 | Component 1 | 0.018 | 0.003 | 0.00 | n.a. | MB |
| 15 | 30.15 | | 0.411 | 0.242 | 0.11 | n.a. | BMB |
| 16 | 31.98 | | 0.011 | 0.005 | 0.00 | n.a. | BMB |
| 17 | 37.13 | | 0.148 | 0.125 | 0.06 | n.a. | BMB |
| 18 | 37.26 | | 0.020 | 0.001 | 0.00 | n.a. | Rd |
| 19 | 40.88 | | 0.019 | 0.001 | 0.00 | n.a. | BMB |
| 20 | 41.83 | | 0.018 | 0.001 | 0.00 | n.a. | BMB |
| Total: | | | 204.388 | 221.303 | 100.00 | 0.000 | |

**913B**

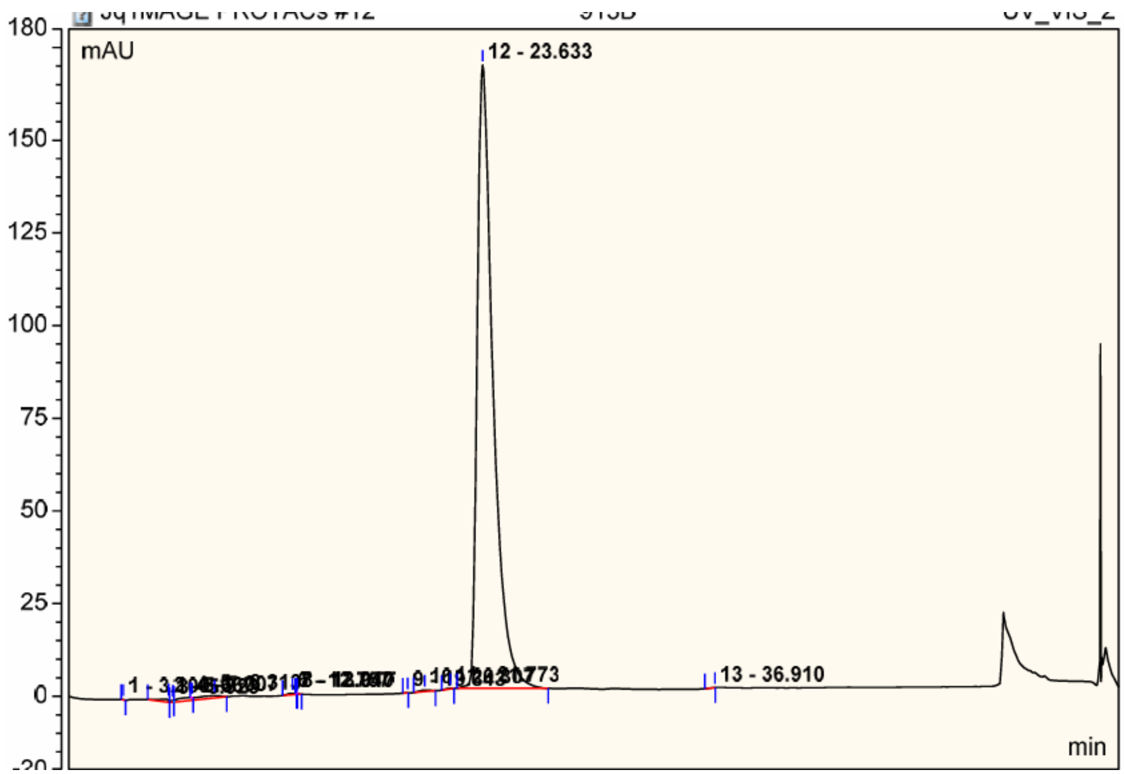

| No. | Ret.Time min | Peak Name | Height mAU | Area mAU*min | Rel.Area % | Amount n.a. | Type |
|---|---|---|---|---|---|---|---|
| 1 | 3.10 | | 0.101 | 0.012 | 0.01 | n.a. | BMB |
| 2 | 5.67 | | 0.917 | 0.525 | 0.29 | n.a. | BMB |
| 3 | 5.92 | | 0.532 | 0.098 | 0.05 | n.a. | BM |
| 4 | 7.01 | | 0.021 | 0.002 | 0.00 | n.a. | Ru |
| 5 | 8.31 | | 0.629 | 1.860 | 1.03 | n.a. | MB |
| 6 | 12.76 | | 0.409 | 0.257 | 0.14 | n.a. | BMB |
| 7 | 12.94 | | 0.010 | 0.000 | 0.00 | n.a. | Rd |
| 8 | 13.02 | | 0.011 | 0.000 | 0.00 | n.a. | Rd |
| 9 | 19.34 | | 0.019 | 0.005 | 0.00 | n.a. | BMB |
| 10 | 20.31 | | 0.406 | 0.289 | 0.16 | n.a. | BMB |
| 11 | 21.77 | | 0.160 | 0.084 | 0.05 | n.a. | BMB |
| 12 | 23.63 | | 168.153 | 177.247 | 98.26 | n.a. | BMB |
| 13 | 36.91 | | 0.017 | 0.013 | 0.01 | n.a. | BMB |
| Total: | | | 171.385 | 180.393 | 100.00 | 0.000 | |

**913C**

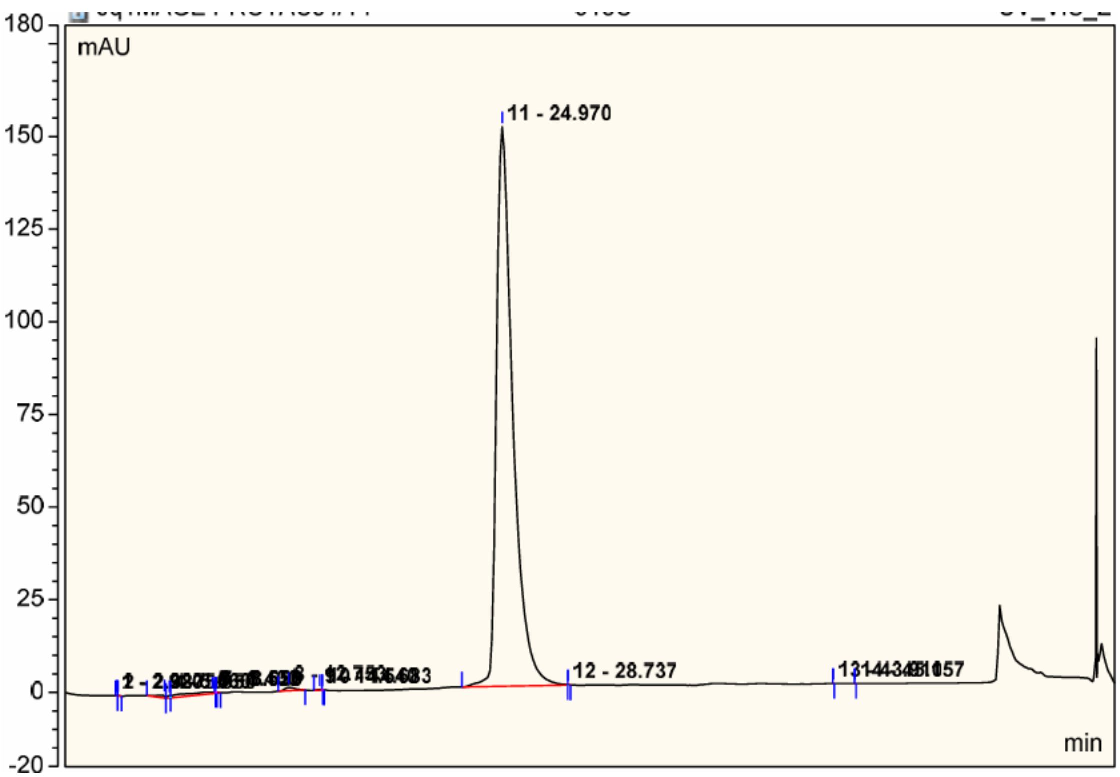

| No. | Ret.Time min | Peak Name | Height mAU | Area mAU*min | Rel.Area % | Amount n.a. | Type |
|---|---|---|---|---|---|---|---|
| 1 | 2.92 | | 0.031 | 0.002 | 0.00 | n.a. | BMb |
| 2 | 2.99 | | 0.013 | 0.012 | 0.01 | n.a. | bMB |
| 3 | 5.66 | | 0.831 | 0.411 | 0.25 | n.a. | BMB |
| 4 | 5.95 | | 0.582 | 0.125 | 0.07 | n.a. | BM |
| 5 | 8.40 | | 0.398 | 1.496 | 0.89 | n.a. | MB |
| 6 | 8.55 | | 0.017 | 0.000 | 0.00 | n.a. | Rd |
| 7 | 8.62 | | 0.017 | 0.001 | 0.00 | n.a. | Rd |
| 8 | 12.75 | | 0.847 | 0.615 | 0.37 | n.a. | BMB |
| 9 | 14.54 | | 0.188 | 0.066 | 0.04 | n.a. | BMB |
| 10 | 14.68 | | 0.015 | 0.000 | 0.00 | n.a. | Rd |
| 11 | 24.97 | | 150.937 | 164.923 | 98.37 | n.a. | BM |
| 12 | 28.74 | | 0.076 | 0.007 | 0.00 | n.a. | MB |
| 13 | 43.91 | | 0.031 | 0.001 | 0.00 | n.a. | BMB |
| 14 | 45.16 | | 0.027 | 0.001 | 0.00 | n.a. | BMB |
| Total: | | | 154.009 | 167.662 | 100.00 | 0.000 | |

**1024F**

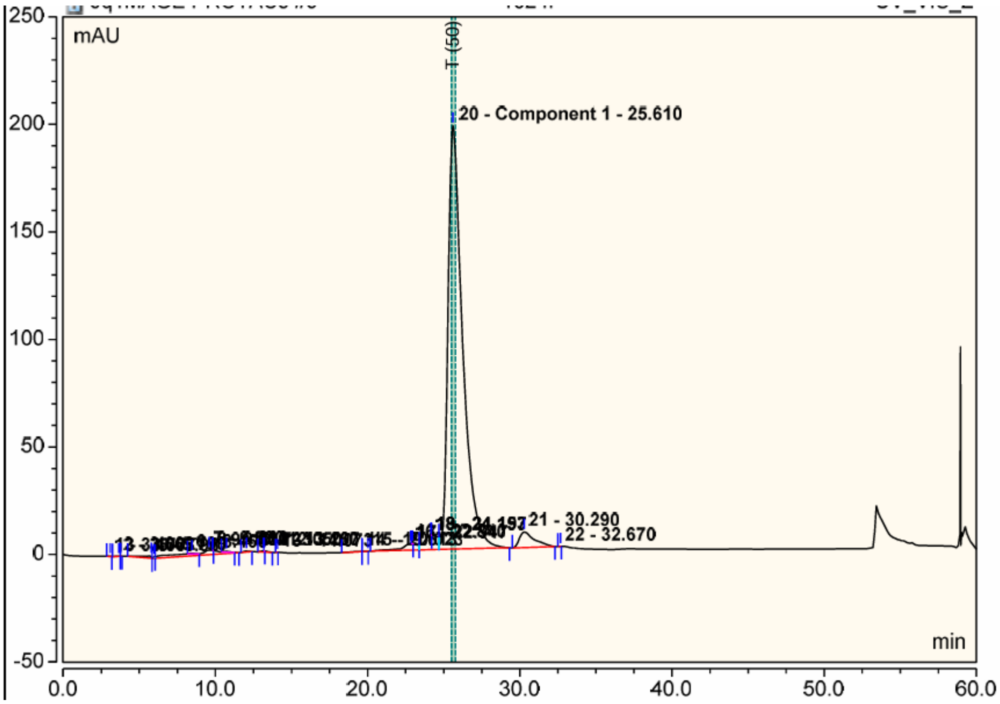

| No. | Ret.Time min | Peak Name | Height mAU | Area mAU*min | Rel.Area % | Amount micromol | Type |
|---|---|---|---|---|---|---|---|
| 1 | 3.10 | | 0.157 | 0.030 | 0.01 | n.a. | BMB |
| 2 | 3.66 | | 0.096 | 0.031 | 0.01 | n.a. | BM |
| 3 | 3.78 | | 0.055 | 0.003 | 0.00 | n.a. | MB |
| 4 | 5.79 | | 1.115 | 0.832 | 0.34 | n.a. | BMB |
| 5 | 5.97 | | 0.652 | 0.085 | 0.03 | n.a. | BM |
| 6 | 8.46 | | 0.319 | 0.125 | 0.05 | n.a. | Ru |
| 7 | 9.59 | | 2.137 | 5.414 | 2.21 | n.a. | MB |
| 8 | 9.83 | | 0.008 | 0.000 | 0.00 | n.a. | Rd |
| 9 | 10.73 | | 0.359 | 0.160 | 0.07 | n.a. | Rd |
| 10 | 12.05 | | 0.432 | 0.176 | 0.07 | n.a. | BMB |
| 11 | 13.16 | | 0.504 | 0.258 | 0.11 | n.a. | BMB |
| 12 | 13.21 | | 0.011 | 0.001 | 0.00 | n.a. | Rd |
| 13 | 14.07 | | 0.021 | 0.002 | 0.00 | n.a. | BMB |
| 14 | 19.61 | | 0.046 | 0.086 | 0.04 | n.a. | BM |
| 15 | 20.02 | | 0.009 | 0.002 | 0.00 | n.a. | MB |
| 16 | 22.84 | | 3.030 | 3.618 | 1.48 | n.a. | BM |
| 17 | 22.95 | | 0.022 | 0.002 | 0.00 | n.a. | Rd |
| 18 | 24.15 | | 6.631 | 6.736 | 2.75 | n.a. | M |
| 19 | 24.20 | | 0.007 | 0.000 | 0.00 | n.a. | Rd |
| 20 | 25.61 | Component 1 | 196.857 | 218.698 | 89.42 | n.a. | MB |
| 21 | 30.29 | | 7.269 | 8.319 | 3.40 | n.a. | BMB |
| 22 | 32.67 | | 0.040 | 0.006 | 0.00 | n.a. | BMB |
| Total: | | | 219.778 | 244.584 | 100.00 | 0.000 | |

**105D**

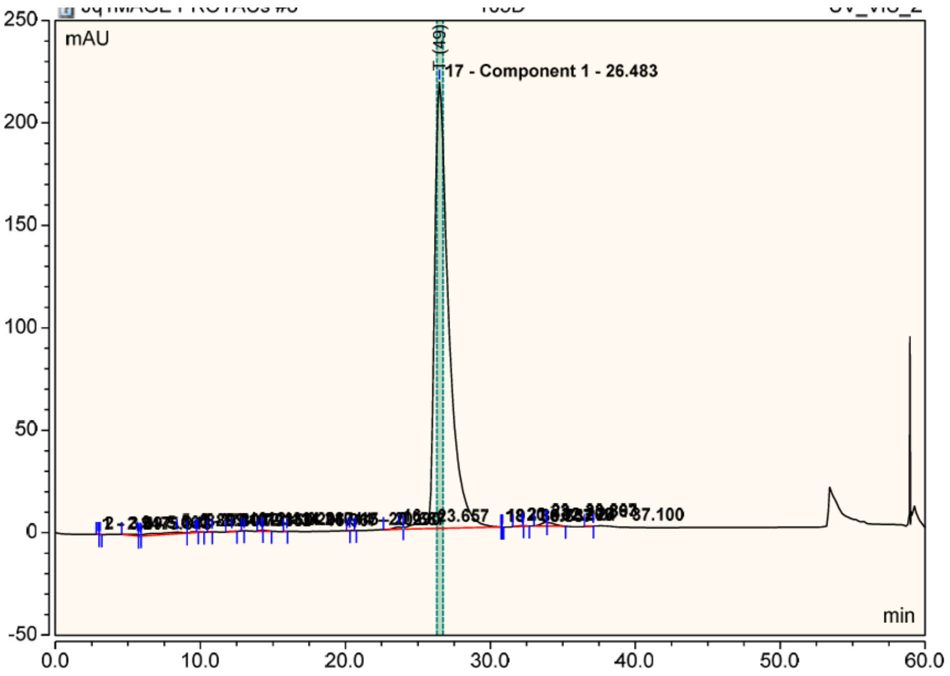

| No. | Ret.Time min | Peak Name | Height mAU | Area mAU*min | Rel.Area % | Amount micromol | Type |
|---|---|---|---|---|---|---|---|
| 1 | 2.95 | | 0.028 | 0.002 | 0.00 | n.a. | BMb |
| 2 | 3.10 | | 0.088 | 0.010 | 0.00 | n.a. | bMB |
| 3 | 5.66 | | 0.906 | 0.482 | 0.19 | n.a. | BMB |
| 4 | 5.84 | | 0.496 | 0.061 | 0.02 | n.a. | BM |
| 5 | 8.37 | | 0.809 | 2.226 | 0.88 | n.a. | M |
| 6 | 9.64 | | 0.468 | 0.377 | 0.15 | n.a. | MB |
| 7 | 9.81 | | 0.015 | 0.001 | 0.00 | n.a. | Rd |
| 8 | 10.79 | | 0.032 | 0.012 | 0.00 | n.a. | BMB |
| 9 | 12.15 | | 0.202 | 0.086 | 0.03 | n.a. | BMB |
| 10 | 13.02 | | 0.000 | 0.002 | 0.00 | n.a. | BMB |
| 11 | 14.24 | | 0.497 | 0.270 | 0.11 | n.a. | BMB |
| 12 | 14.29 | | 0.004 | 0.000 | 0.00 | n.a. | Rd |
| 13 | 15.97 | | 0.030 | 0.007 | 0.00 | n.a. | BMB |
| 14 | 20.29 | | 0.017 | 0.003 | 0.00 | n.a. | BMB |
| 15 | 20.69 | | 0.023 | 0.001 | 0.00 | n.a. | BMB |
| 16 | 23.66 | | 1.057 | 0.749 | 0.30 | n.a. | BM |
| 17 | 26.48 | Component 1 | 217.727 | 247.478 | 97.47 | n.a. | M |
| 18 | 30.76 | | 0.069 | 0.004 | 0.00 | n.a. | M |
| 19 | 30.84 | | 0.046 | 0.003 | 0.00 | n.a. | MB |
| 20 | 32.20 | | 0.309 | 0.211 | 0.08 | n.a. | BMB |
| 21 | 32.27 | | 0.006 | 0.000 | 0.00 | n.a. | Rd |
| 22 | 33.87 | | 1.817 | 1.886 | 0.74 | n.a. | BMB |
| 23 | 33.89 | | 0.009 | 0.000 | 0.00 | n.a. | Rd |
| 24 | 37.10 | | 0.013 | 0.020 | 0.01 | n.a. | BMB |
| Total: | | | 224.670 | 253.891 | 100.00 | 0.000 | |

**913D**

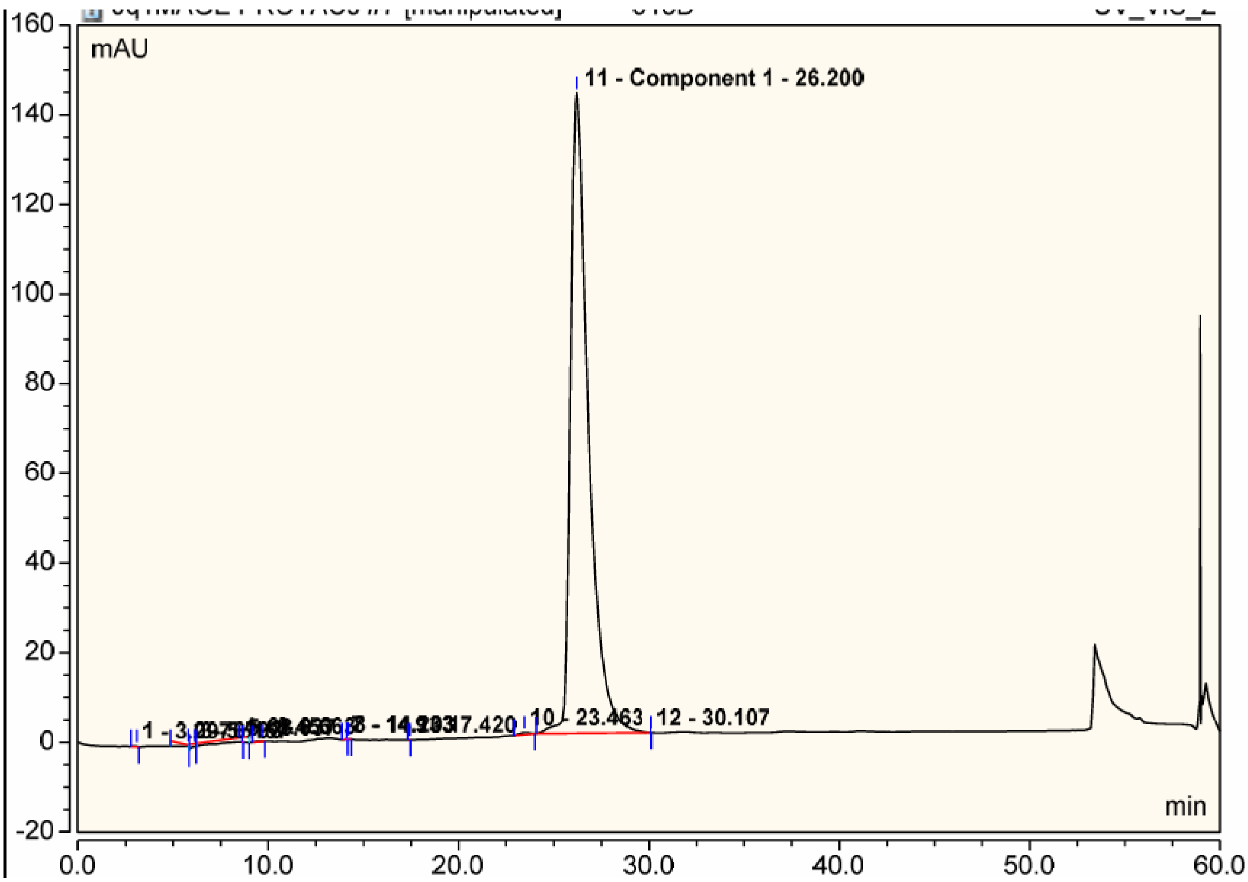

| No. | Ret.Time min | Peak Name | Height mAU | Area mAU*min | Rel.Area % | Amount micromol | Type |
|---|---|---|---|---|---|---|---|
| 1 | 3.10 | | 0.151 | 0.034 | 0.02 | n.a. | BMB |
| 2 | 5.79 | | 0.141 | 0.718 | 0.45 | n.a. | M * |
| 3 | 6.16 | | 0.497 | 0.262 | 0.16 | n.a. | M * |
| 4 | 8.45 | | 0.875 | 2.086 | 1.30 | n.a. | M * |
| 5 | 8.64 | | 0.015 | 0.001 | 0.00 | n.a. | Rd |
| 6 | 9.66 | | 0.081 | 0.048 | 0.03 | n.a. | BMB |
| 7 | 14.13 | | 0.010 | 0.000 | 0.00 | n.a. | Ru |
| 8 | 14.20 | | 0.083 | 0.021 | 0.01 | n.a. | BMB |
| 9 | 17.42 | | 0.023 | 0.002 | 0.00 | n.a. | BMB |
| 10 | 23.46 | | 0.445 | 0.268 | 0.17 | n.a. | BMB |
| 11 | 26.20 | Component 1 | 142.813 | 156.517 | 97.85 | n.a. | BM |
| 12 | 30.11 | | 0.030 | 0.001 | 0.00 | n.a. | MB |
| Total: | | | 145.164 | 159.958 | 100.00 | 0.000 | |

**105A**

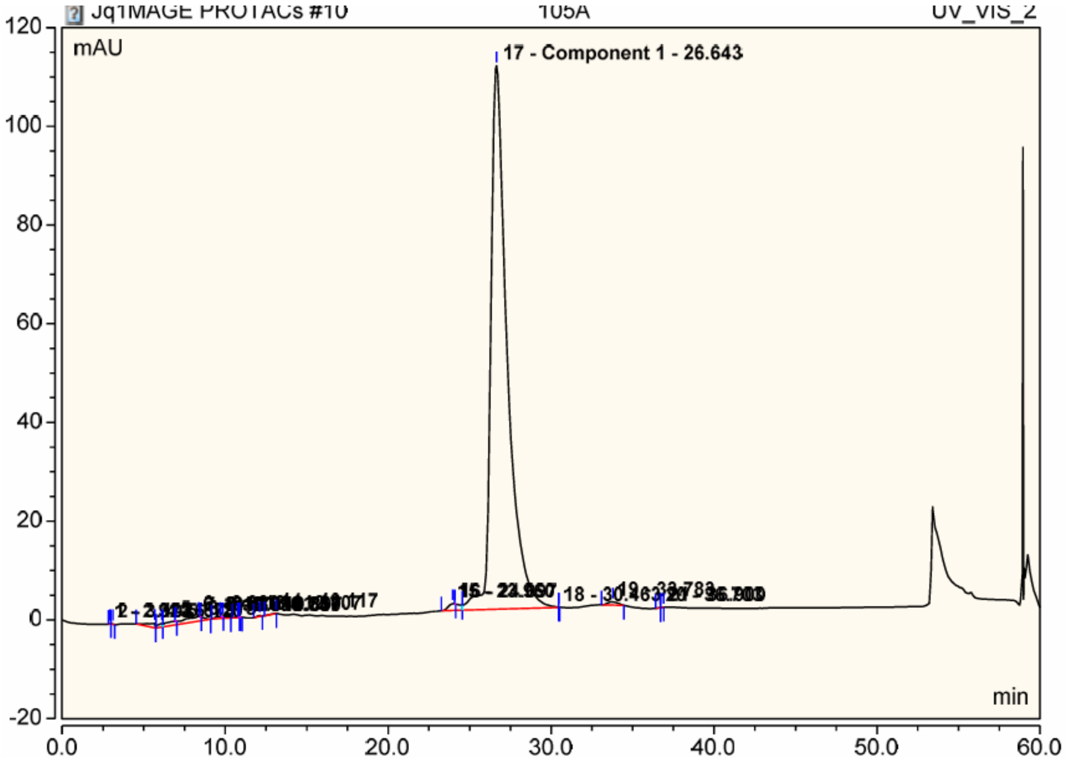

| No. | Ret.Time min | Peak Name | Height mAU | Area mAU*min | Rel.Area % | Amount micromol | Type |
|---|---|---|---|---|---|---|---|
| 1 | 2.92 | | 0.045 | 0.003 | 0.00 | n.a. | BMb |
| 2 | 3.10 | | 0.118 | 0.013 | 0.01 | n.a. | bMB |
| 3 | 5.67 | | 0.906 | 0.508 | 0.38 | n.a. | BMB |
| 4 | 6.12 | | 0.746 | 0.238 | 0.18 | n.a. | BM |
| 5 | 6.97 | | 0.016 | 0.001 | 0.00 | n.a. | Ru |
| 6 | 8.38 | | 0.915 | 1.943 | 1.46 | n.a. | MB |
| 7 | 8.47 | | 0.021 | 0.001 | 0.00 | n.a. | Rd |
| 8 | 9.69 | | 0.377 | 0.273 | 0.21 | n.a. | BMB |
| 9 | 9.84 | | 0.021 | 0.002 | 0.00 | n.a. | Rd |
| 10 | 10.76 | | 0.168 | 0.056 | 0.04 | n.a. | BMB |
| 11 | 10.84 | | 0.018 | 0.001 | 0.00 | n.a. | Rd |
| 12 | 10.90 | | 0.013 | 0.000 | 0.00 | n.a. | Rd |
| 13 | 12.11 | | 0.179 | 0.062 | 0.05 | n.a. | BMB |
| 14 | 13.12 | | 0.035 | 0.041 | 0.03 | n.a. | BMB |
| 15 | 23.96 | | 1.392 | 1.126 | 0.84 | n.a. | BM |
| 16 | 24.10 | | 0.027 | 0.001 | 0.00 | n.a. | Rd |
| 17 | 26.64 | Component 1 | 110.114 | 128.567 | 96.44 | n.a. | M |
| 18 | 30.46 | | 0.028 | 0.001 | 0.00 | n.a. | MB |
| 19 | 33.78 | | 0.641 | 0.469 | 0.35 | n.a. | BMB |
| 20 | 36.70 | | 0.013 | 0.004 | 0.00 | n.a. | BM |
| 21 | 36.90 | | 0.016 | 0.002 | 0.00 | n.a. | MB |
| Total: | | | 115.811 | 133.313 | 100.00 | 0.000 | |

**105E**

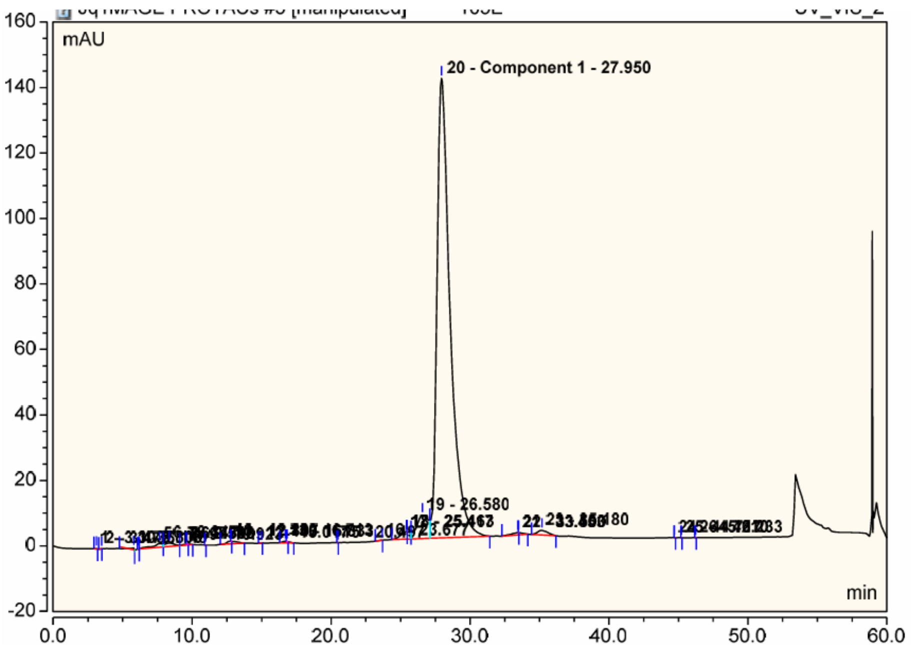

| No. | Ret.Time min | Peak Name | Height mAU | Area mAU*min | Rel.Area % | Amount micromol | Type |
|---|---|---|---|---|---|---|---|
| 1 | 3.10 | | 0.172 | 0.021 | 0.01 | n.a. | BMB |
| 2 | 3.48 | | 0.011 | 0.004 | 0.00 | n.a. | BMB |
| 3 | 5.80 | | 0.539 | 0.101 | 0.06 | n.a. | M * |
| 4 | 6.14 | | 0.194 | 0.011 | 0.01 | n.a. | BM * |
| 5 | 7.65 | | 1.158 | 0.997 | 0.60 | n.a. | M |
| 6 | 8.35 | | 1.055 | 0.830 | 0.50 | n.a. | M |
| 7 | 9.54 | | 0.317 | 0.198 | 0.12 | n.a. | MB |
| 8 | 9.59 | | 0.019 | 0.002 | 0.00 | n.a. | Rd |
| 9 | 10.92 | | 0.023 | 0.001 | 0.00 | n.a. | BMB |
| 10 | 12.73 | | 0.836 | 0.691 | 0.41 | n.a. | BMB |
| 11 | 12.81 | | 0.015 | 0.001 | 0.00 | n.a. | Rd |
| 12 | 15.02 | | 0.041 | 0.010 | 0.01 | n.a. | BMB |
| 13 | 16.71 | | 0.459 | 0.241 | 0.14 | n.a. | BMB |
| 14 | 16.83 | | 0.027 | 0.003 | 0.00 | n.a. | Rd |
| 15 | 20.50 | | 0.018 | 0.001 | 0.00 | n.a. | BMB |
| 16 | 23.68 | | 0.020 | 0.024 | 0.01 | n.a. | BMB |
| 17 | 25.42 | | 2.212 | 1.763 | 1.05 | n.a. | BM |
| 18 | 25.46 | | 0.012 | 0.000 | 0.00 | n.a. | Rd |
| 19 | 26.58 | | 7.195 | 6.795 | 4.06 | n.a. | M |
| 20 | 27.95 | Component 1 | 140.362 | 153.554 | 91.85 | n.a. | MB |
| 21 | 33.45 | | 0.749 | 0.715 | 0.43 | n.a. | BMB |
| 22 | 33.50 | | 0.004 | 0.000 | 0.00 | n.a. | Rd |
| 23 | 35.18 | | 1.400 | 1.218 | 0.73 | n.a. | BMB |
| 24 | 44.72 | | 0.020 | 0.001 | 0.00 | n.a. | BMB |
| 25 | 45.21 | | 0.022 | 0.001 | 0.00 | n.a. | BMB |
| 26 | 46.23 | | 0.020 | 0.001 | 0.00 | n.a. | BMB |
| Total: | | | 156.899 | 167.185 | 100.00 | 0.000 | |

**105B**

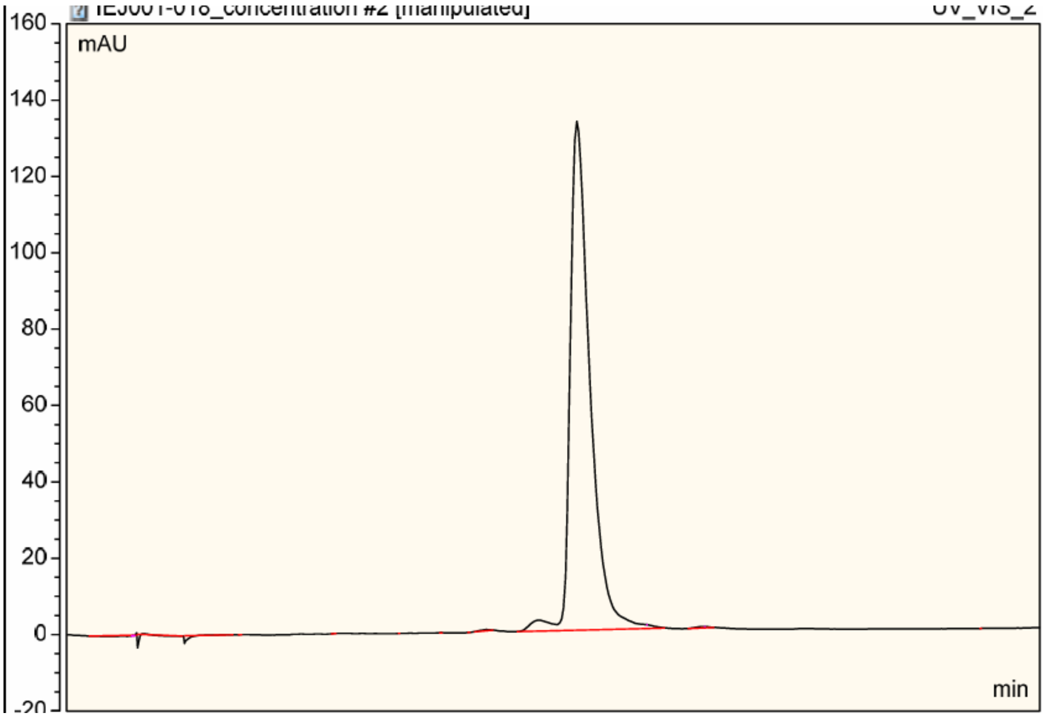

| No. | Ret.Time min | Peak Name | Height mAU | Area mAU*min | Rel.Area % | Amount n.a. | Type |
|---|---|---|---|---|---|---|---|
| 1 | 3.40 | | 0.095 | 0.010 | 0.01 | n.a. | Ru |
| 2 | 3.58 | | 0.848 | 0.447 | 0.28 | n.a. | BM * |
| 3 | 3.89 | | 0.187 | 0.335 | 0.21 | n.a. | MB* |
| 4 | 8.16 | | 0.105 | 0.281 | 0.17 | n.a. | M * |
| 5 | 8.42 | | 0.049 | 0.004 | 0.00 | n.a. | MB* |
| 6 | 8.99 | | 0.013 | 0.002 | 0.00 | n.a. | BMB |
| 7 | 13.79 | | 0.036 | 0.006 | 0.00 | n.a. | BMB |
| 8 | 17.07 | | 0.019 | 0.001 | 0.00 | n.a. | BMB |
| 9 | 19.29 | | 0.019 | 0.004 | 0.00 | n.a. | BMB |
| 10 | 20.71 | | 0.012 | 0.002 | 0.00 | n.a. | BMB |
| 11 | 21.54 | | 0.345 | 0.197 | 0.12 | n.a. | BMB |
| 12 | 24.30 | | 2.848 | 3.543 | 2.21 | n.a. | BM |
| 13 | 26.20 | | 133.290 | 154.698 | 96.45 | n.a. | M |
| 14 | 29.67 | | 1.084 | 0.572 | 0.36 | n.a. | MB |
| 15 | 29.80 | | 0.015 | 0.001 | 0.00 | n.a. | Rd |
| 16 | 32.68 | | 0.380 | 0.290 | 0.18 | n.a. | BMB |
| 17 | 32.78 | | 0.011 | 0.000 | 0.00 | n.a. | Rd |
| 18 | 46.97 | | 0.018 | 0.001 | 0.00 | n.a. | BMB |
| **Total:** | | | 139.374 | 160.391 | 100.00 | 0.000 | |

**105C**

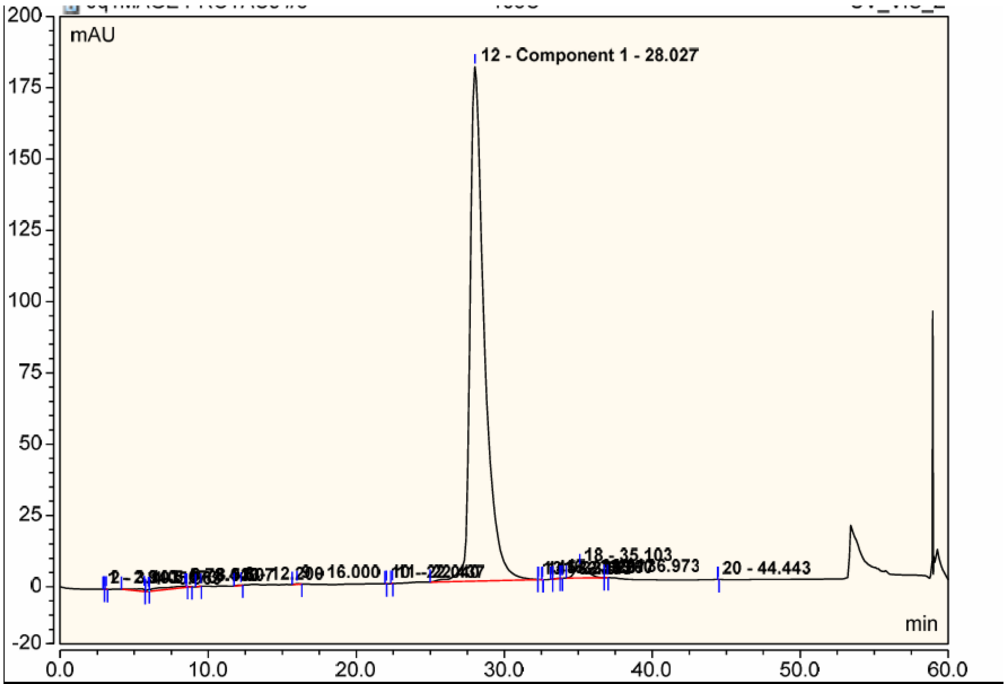

| No. | Ret.Time min | Peak Name | Height mAU | Area mAU*min | Rel.Area % | Amount micromol | Type |
|---|---|---|---|---|---|---|---|
| 1 | 2.94 | | 0.027 | 0.001 | 0.00 | n.a. | BMB |
| 2 | 3.10 | | 0.085 | 0.009 | 0.00 | n.a. | BMB |
| 3 | 5.67 | | 0.988 | 0.730 | 0.35 | n.a. | BMB |
| 4 | 5.96 | | 0.590 | 0.117 | 0.06 | n.a. | BM |
| 5 | 8.44 | | 0.480 | 1.707 | 0.82 | n.a. | MB |
| 6 | 8.53 | | 0.020 | 0.001 | 0.00 | n.a. | Rd |
| 7 | 9.51 | | 0.022 | 0.008 | 0.00 | n.a. | BMB |
| 8 | 12.20 | | 0.097 | 0.050 | 0.02 | n.a. | BMB |
| 9 | 16.00 | | 0.215 | 0.083 | 0.04 | n.a. | BMB |
| 10 | 22.04 | | 0.021 | 0.001 | 0.00 | n.a. | BMB |
| 11 | 22.44 | | 0.020 | 0.002 | 0.00 | n.a. | BMB |
| 12 | 28.03 | Component 1 | 180.393 | 202.740 | 96.80 | n.a. | BM |
| 13 | 32.29 | | 0.118 | 0.019 | 0.01 | n.a. | MB |
| 14 | 32.58 | | 0.018 | 0.001 | 0.00 | n.a. | Rd |
| 15 | 33.27 | | 0.046 | 0.003 | 0.00 | n.a. | BM |
| 16 | 33.76 | | 0.077 | 0.039 | 0.02 | n.a. | M |
| 17 | 33.92 | | 0.026 | 0.006 | 0.00 | n.a. | MB |
| 18 | 35.10 | | 3.905 | 3.928 | 1.88 | n.a. | BM |
| 19 | 36.97 | | 0.013 | 0.002 | 0.00 | n.a. | MB |
| 20 | 44.44 | | 0.020 | 0.001 | 0.00 | n.a. | BMB |
| Total: | | | 187.181 | 209.448 | 100.00 | 0.000 | |

**1024C**

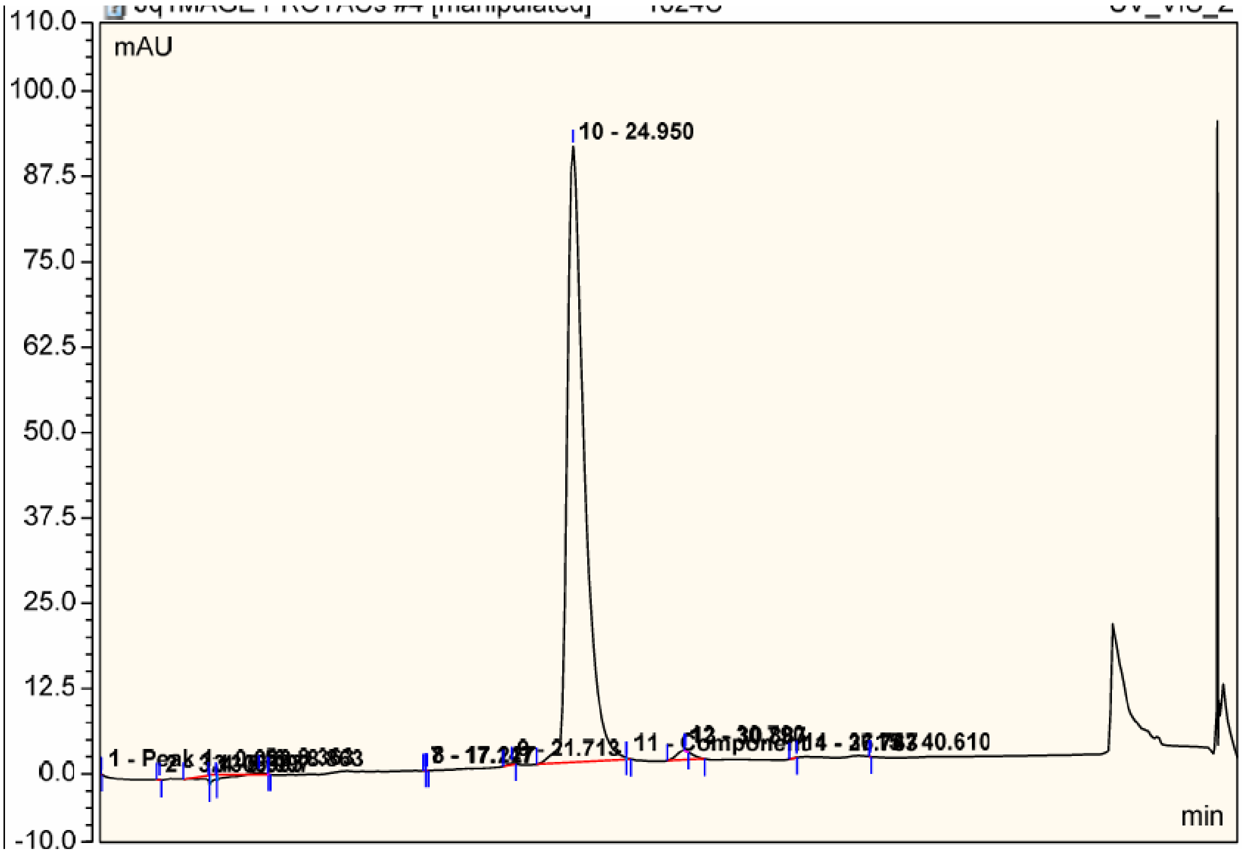

| No. | Ret.Time min | Peak Name | Height mAU | Area mAU*min | Rel.Area % | Amount micromol | Type |
|---|---|---|---|---|---|---|---|
| 1 | 0.06 | Peak 1 | 0.057 | 0.002 | 0.00 | n.a. | BMB |
| 2 | 3.13 | | 0.093 | 0.010 | 0.01 | n.a. | BMB |
| 3 | 5.68 | | 0.517 | 0.413 | 0.42 | n.a. | BM * |
| 4 | 6.10 | | 0.703 | 0.394 | 0.40 | n.a. | M * |
| 5 | 8.36 | | 0.294 | 0.358 | 0.37 | n.a. | MB |
| 6 | 8.86 | | 0.010 | 0.000 | 0.00 | n.a. | Rd |
| 7 | 17.15 | | 0.037 | 0.003 | 0.00 | n.a. | BM |
| 8 | 17.23 | | 0.032 | 0.003 | 0.00 | n.a. | MB |
| 9 | 21.71 | | 0.167 | 0.071 | 0.07 | n.a. | BMB |
| 10 | 24.95 | | 90.223 | 95.551 | 97.36 | n.a. | BM |
| 11 | 27.76 | Component 1 | 0.232 | 0.031 | 0.03 | n.a. | MB |
| 12 | 30.79 | | 1.398 | 1.285 | 1.31 | n.a. | BMB |
| 13 | 30.89 | | 0.053 | 0.008 | 0.01 | n.a. | Rd |
| 14 | 36.75 | | 0.017 | 0.006 | 0.01 | n.a. | BMB |
| 15 | 40.61 | | 0.019 | 0.001 | 0.00 | n.a. | BMB |
| Total: | | | 93.851 | 98.137 | 100.00 | 0.000 | |

**1024B**

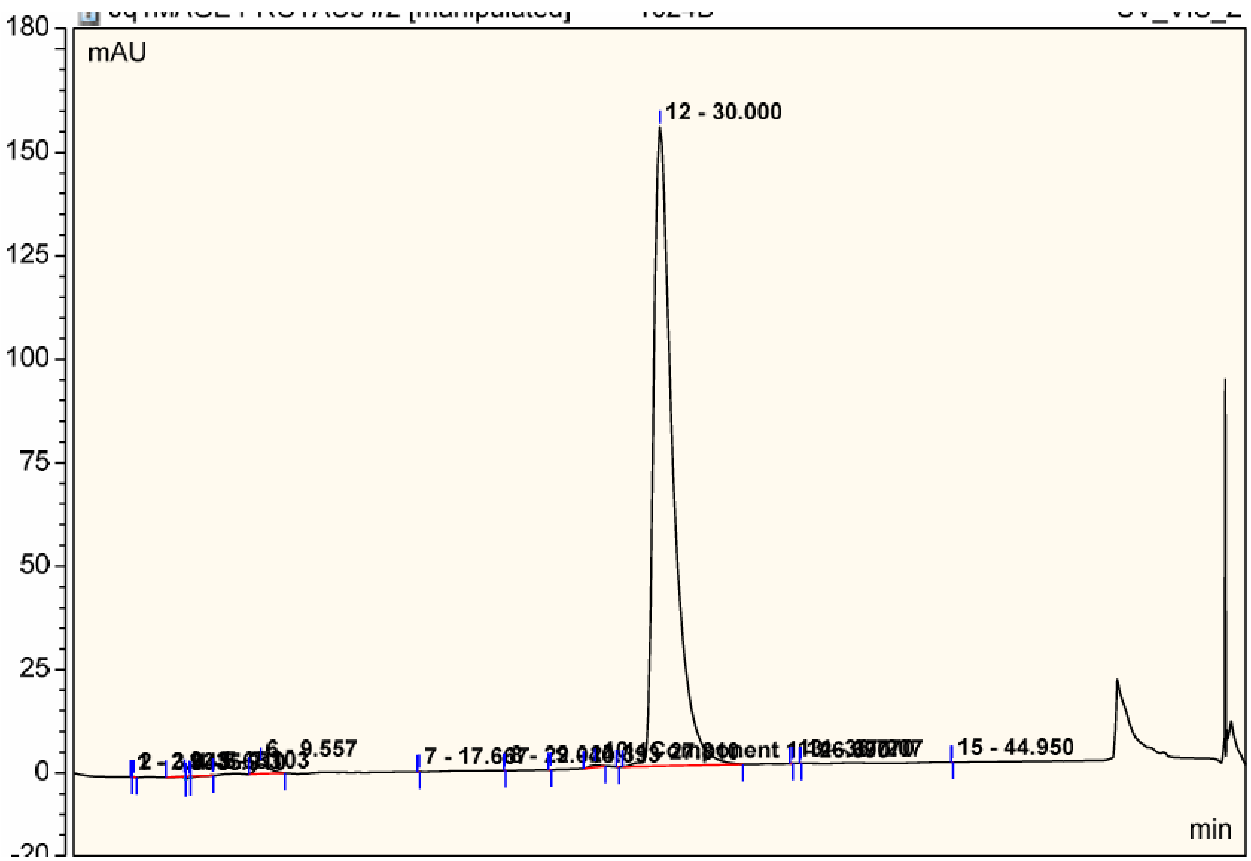

| No. | Ret.Time min | Peak Name | Height mAU | Area mAU*min | Rel.Area % | Amount micromol | Type |
|---|---|---|---|---|---|---|---|
| 1 | 2.93 | | 0.027 | 0.001 | 0.00 | n.a. | BMb |
| 2 | 3.04 | | 0.064 | 0.013 | 0.01 | n.a. | bMB |
| 3 | 5.65 | | 0.025 | 0.066 | 0.04 | n.a. | BM * |
| 4 | 5.91 | | 0.285 | 0.112 | 0.06 | n.a. | M * |
| 5 | 7.10 | | 0.002 | 0.079 | 0.04 | n.a. | MB |
| 6 | 9.56 | | 2.328 | 1.880 | 1.04 | n.a. | BMB |
| 7 | 17.67 | | 0.020 | 0.001 | 0.00 | n.a. | BMB |
| 8 | 22.04 | | 0.021 | 0.001 | 0.00 | n.a. | BMB |
| 9 | 24.39 | | 0.019 | 0.001 | 0.00 | n.a. | BMB |
| 10 | 26.69 | Component 1 | 0.468 | 0.298 | 0.16 | n.a. | BMB |
| 11 | 27.81 | | 0.015 | 0.001 | 0.00 | n.a. | BMB |
| 12 | 30.00 | | 154.402 | 178.751 | 98.64 | n.a. | BMB |
| 13 | 36.77 | | 0.014 | 0.001 | 0.00 | n.a. | BMB |
| 14 | 37.21 | | 0.022 | 0.001 | 0.00 | n.a. | BMB |
| 15 | 44.95 | | 0.021 | 0.001 | 0.00 | n.a. | BMB |
| **Total:** | | | 157.732 | 181.209 | 100.00 | 0.000 | |

**1024E**

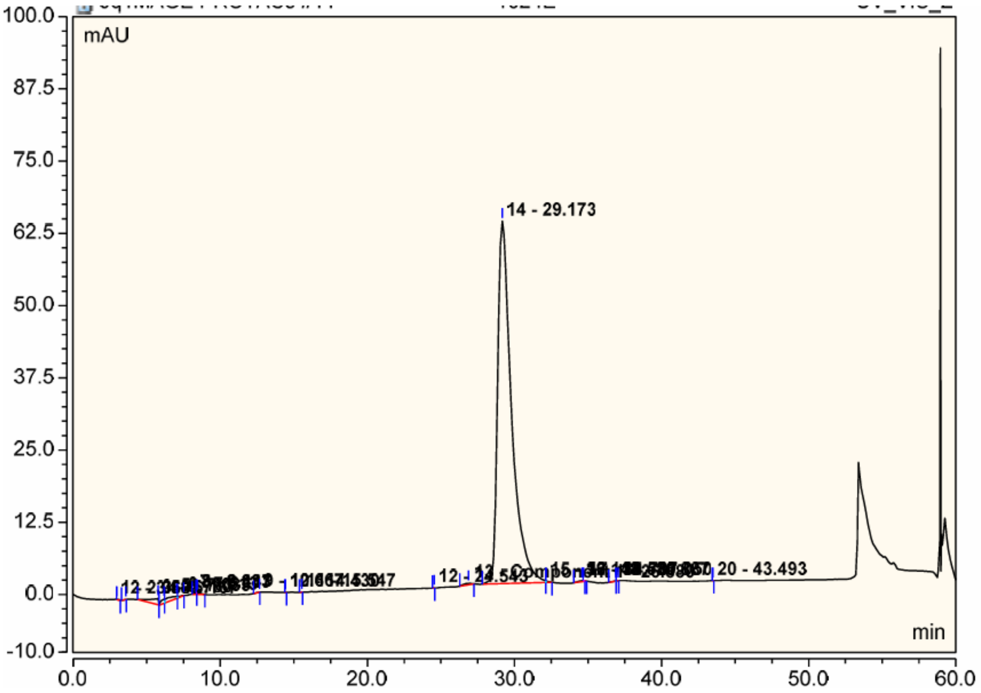

| No. | Ret.Time min | Peak Name | Height mAU | Area mAU*min | Rel.Area % | Amount micromol | Type |
|---|---|---|---|---|---|---|---|
| 1 | 2.96 | | 0.009 | 0.013 | 0.02 | n.a. | BMB |
| 2 | 3.58 | | 0.016 | 0.005 | 0.01 | n.a. | BMB |
| 3 | 5.78 | | 1.107 | 0.721 | 1.02 | n.a. | BMB |
| 4 | 6.17 | | 0.759 | 0.210 | 0.30 | n.a. | BM |
| 5 | 7.03 | | 0.308 | 0.443 | 0.63 | n.a. | M |
| 6 | 7.51 | | 0.023 | 0.053 | 0.07 | n.a. | MB |
| 7 | 8.38 | | 0.012 | 0.000 | 0.00 | n.a. | Ru |
| 8 | 8.45 | | 0.264 | 0.120 | 0.17 | n.a. | BMB |
| 9 | 12.67 | | 0.023 | 0.013 | 0.02 | n.a. | BMB |
| 10 | 14.43 | | 0.018 | 0.001 | 0.00 | n.a. | BMB |
| 11 | 15.55 | | 0.022 | 0.003 | 0.00 | n.a. | BMB |
| 12 | 24.54 | | 0.016 | 0.002 | 0.00 | n.a. | BMB |
| 13 | 26.88 | Component 1 | 0.235 | 0.162 | 0.23 | n.a. | BMB |
| 14 | 29.17 | | 62.740 | 68.471 | 97.27 | n.a. | BM |
| 15 | 32.14 | | 0.276 | 0.054 | 0.08 | n.a. | MB |
| 16 | 34.58 | | 0.199 | 0.100 | 0.14 | n.a. | BMB |
| 17 | 34.71 | | 0.016 | 0.001 | 0.00 | n.a. | Rd |
| 18 | 36.89 | | 0.050 | 0.015 | 0.02 | n.a. | BM |
| 19 | 37.05 | | 0.028 | 0.005 | 0.01 | n.a. | MB |
| 20 | 43.49 | | 0.023 | 0.001 | 0.00 | n.a. | BMB |
| **Total:** | | | 66.143 | 70.394 | 100.00 | 0.000 | |

**1024D**

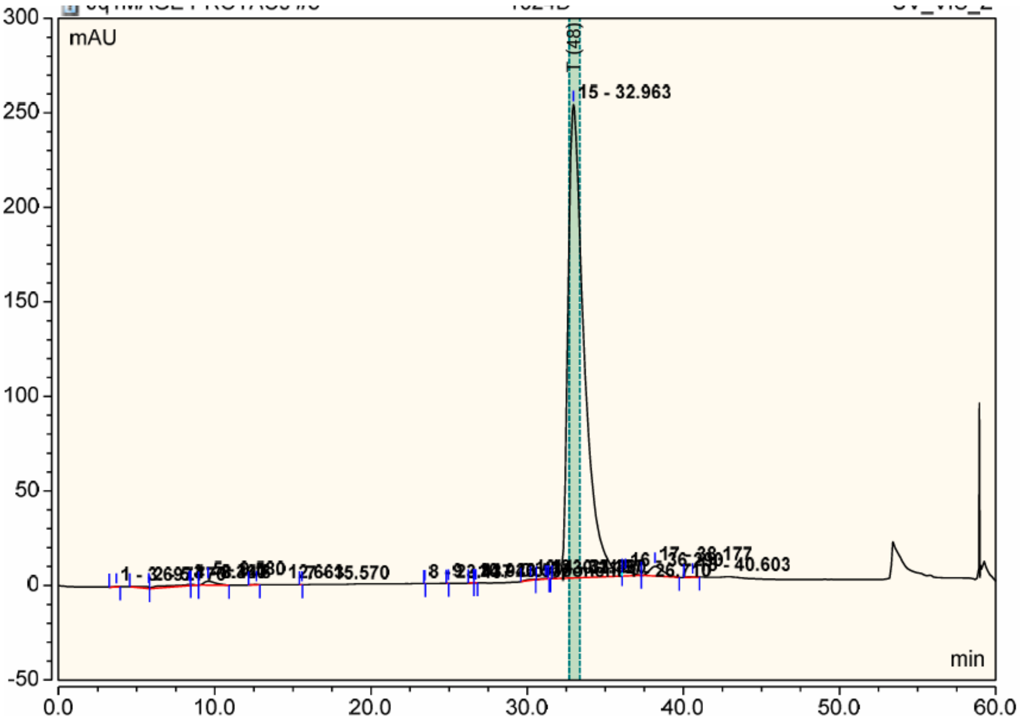

| No. | Ret.Time min | Peak Name | Height mAU | Area mAU*min | Rel.Area % | Amount micromol | Type |
|---|---|---|---|---|---|---|---|
| 1 | 3.70 | | 0.122 | 0.046 | 0.01 | n.a. | BMB |
| 2 | 5.77 | | 1.148 | 0.682 | 0.22 | n.a. | BMB |
| 3 | 8.36 | | 0.747 | 2.078 | 0.67 | n.a. | BMb |
| 4 | 8.45 | | 0.010 | 0.000 | 0.00 | n.a. | Rd |
| 5 | 9.58 | | 2.119 | 1.794 | 0.58 | n.a. | bMB |
| 6 | 12.66 | | 0.120 | 0.058 | 0.02 | n.a. | BMB |
| 7 | 15.57 | | 0.017 | 0.002 | 0.00 | n.a. | BMB |
| 8 | 23.44 | | 0.023 | 0.001 | 0.00 | n.a. | BMB |
| 9 | 24.94 | | 0.015 | 0.002 | 0.00 | n.a. | BMB |
| 10 | 26.58 | | 0.076 | 0.019 | 0.01 | n.a. | BM |
| 11 | 26.71 | Component 1 | 0.065 | 0.010 | 0.00 | n.a. | MB |
| 12 | 30.22 | | 0.412 | 0.272 | 0.09 | n.a. | BMB |
| 13 | 31.20 | | 0.214 | 0.140 | 0.05 | n.a. | BMB |
| 14 | 31.36 | | 0.013 | 0.000 | 0.00 | n.a. | Rd |
| 15 | 32.96 | | 250.490 | 298.299 | 96.14 | n.a. | BM |
| 16 | 36.29 | | 2.050 | 1.550 | 0.50 | n.a. | Mb |
| 17 | 38.18 | | 5.191 | 5.155 | 1.66 | n.a. | bMB |
| 18 | 40.60 | | 0.257 | 0.160 | 0.05 | n.a. | BMB |
| Total: | | | 263.090 | 310.269 | 100.00 | 0.000 | |

#### HRMS/LCMS

**105B**

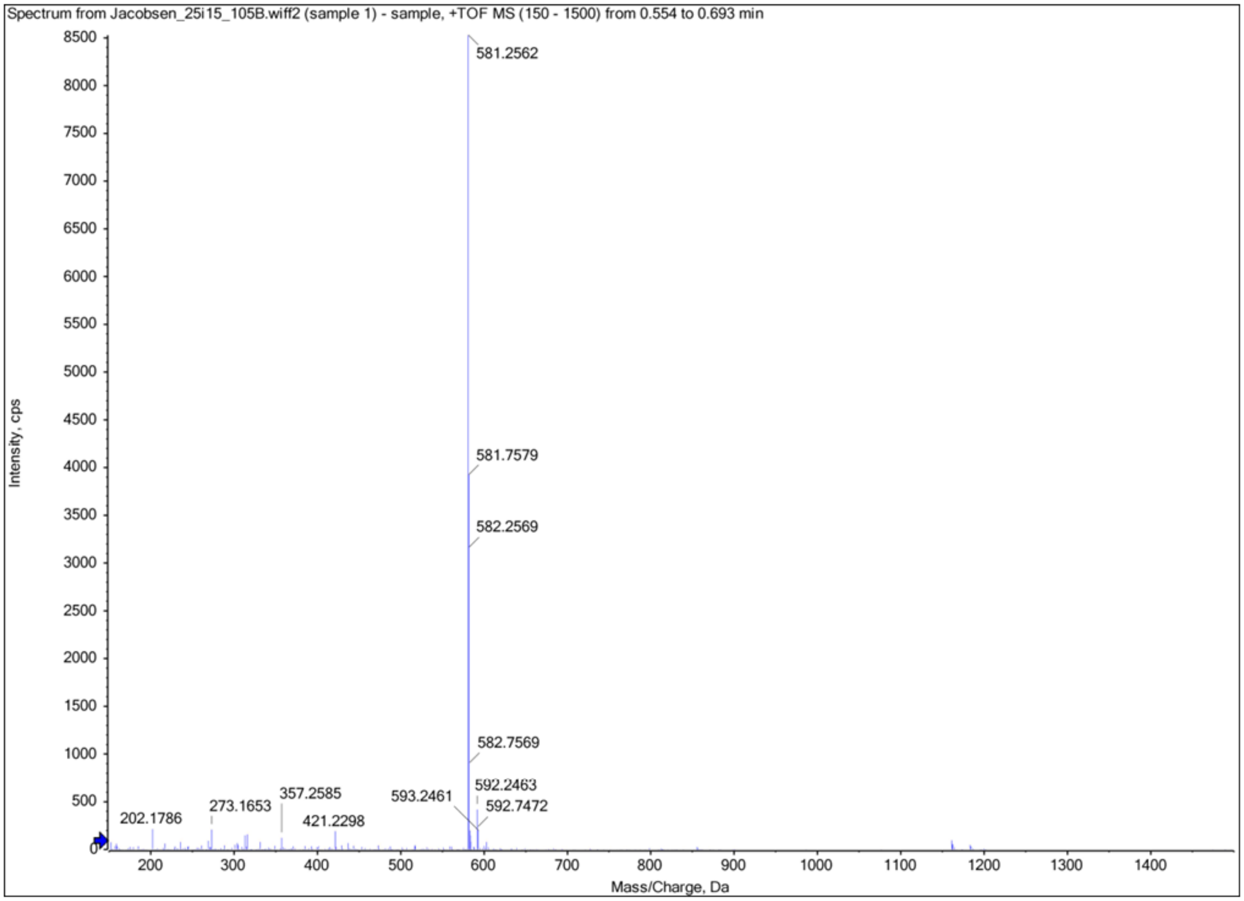

**SJ1008066**

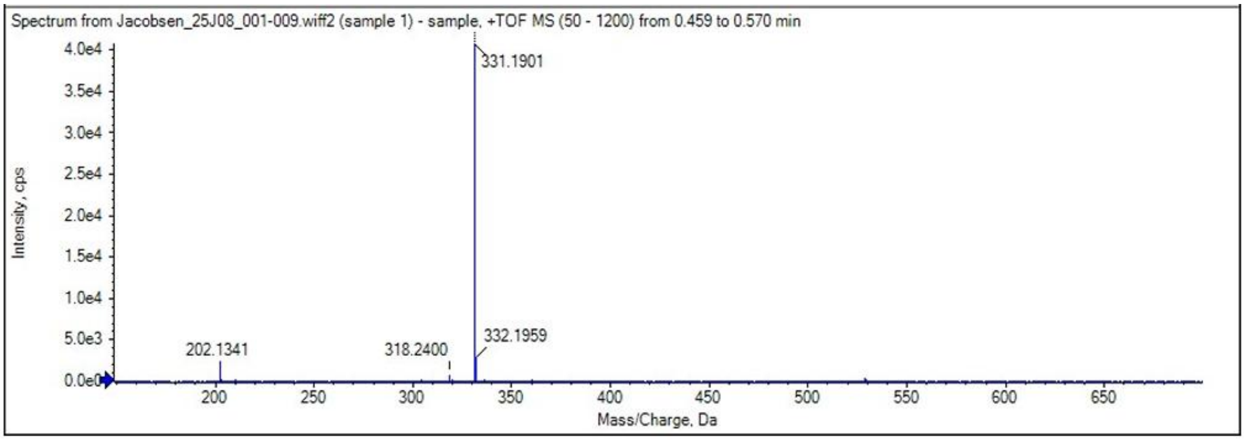

**913B**

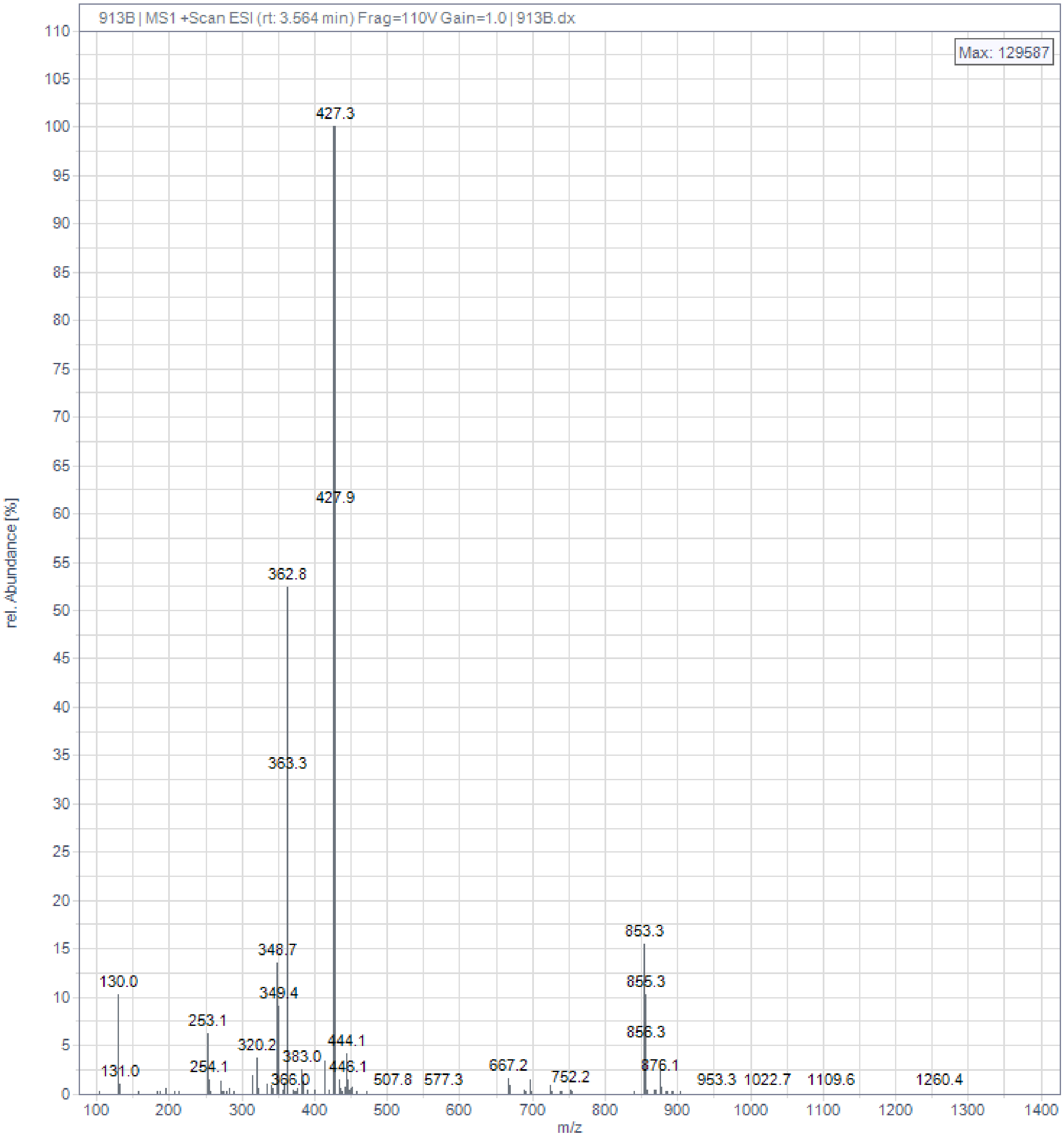
913B | MS1 +Scan ESI (rt: 3.564 min) Frag=110V Gain=1.0 | 913B.dx
Max: 129587
rel. Abundance [%]
m/z
427.3
427.9
362.8
363.3
348.7
349.4
130.0
131.0
253.1
254.1
320.2
383.0
366.0
444.1
446.1
507.8
577.3
667.2
752.2
853.3
855.3
856.3
876.1
953.3
1022.7
1109.6
1260.4

**908B**

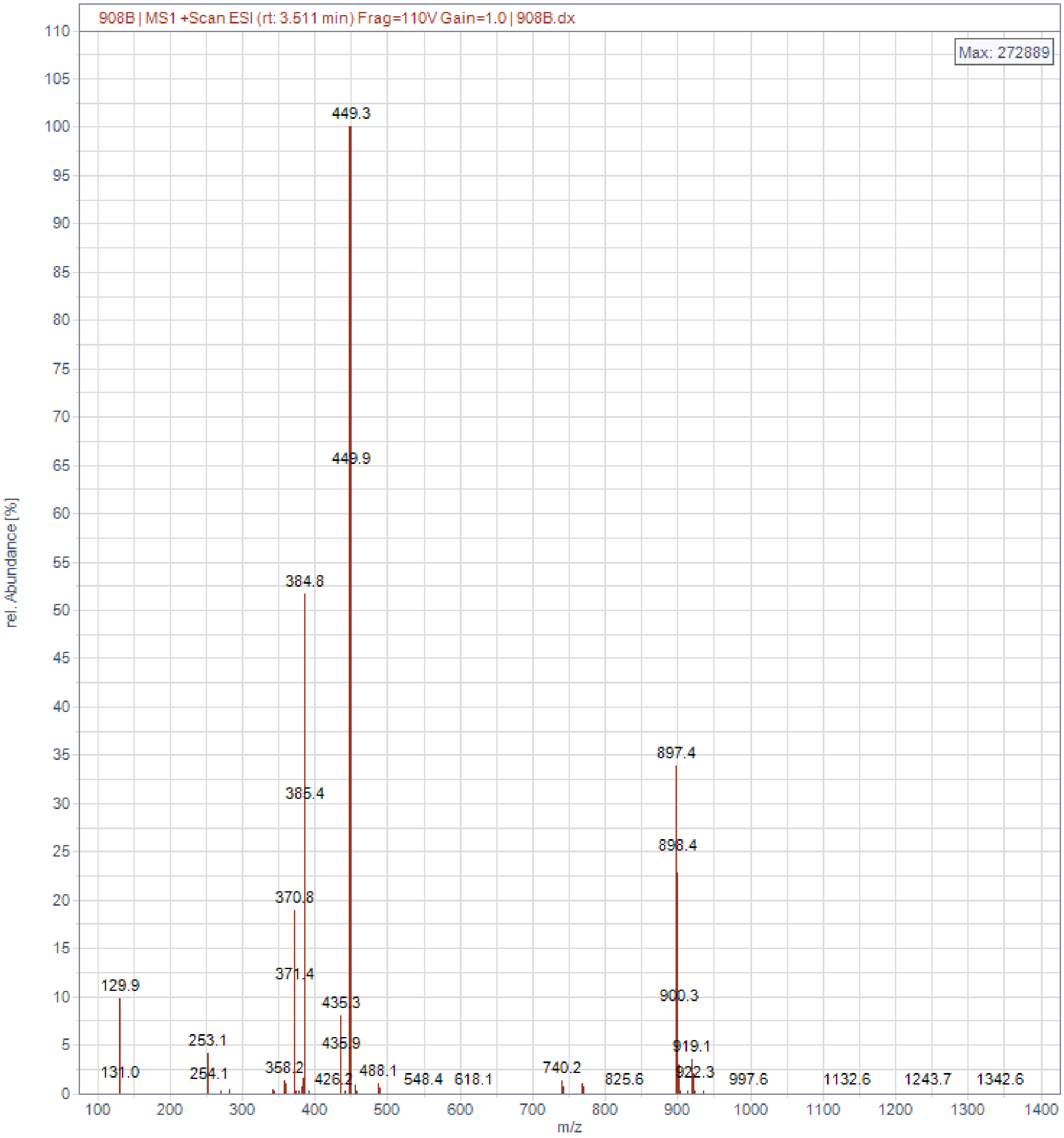
908B | MS1 +Scan ESI (rt: 3.511 min) Frag=110V Gain=1.0 | 908B.dx
Max: 272889
rel. Abundance [%]
m/z
449.3
449.9
384.8
385.4
370.8
371.4
129.9
131.0
253.1
254.1
358.2
426.2
435.3
435.9
488.1
548.4
618.1
740.2
825.6
897.4
898.4
900.3
919.1
922.3
997.6
1132.6
1243.7
1342.6

**913C**

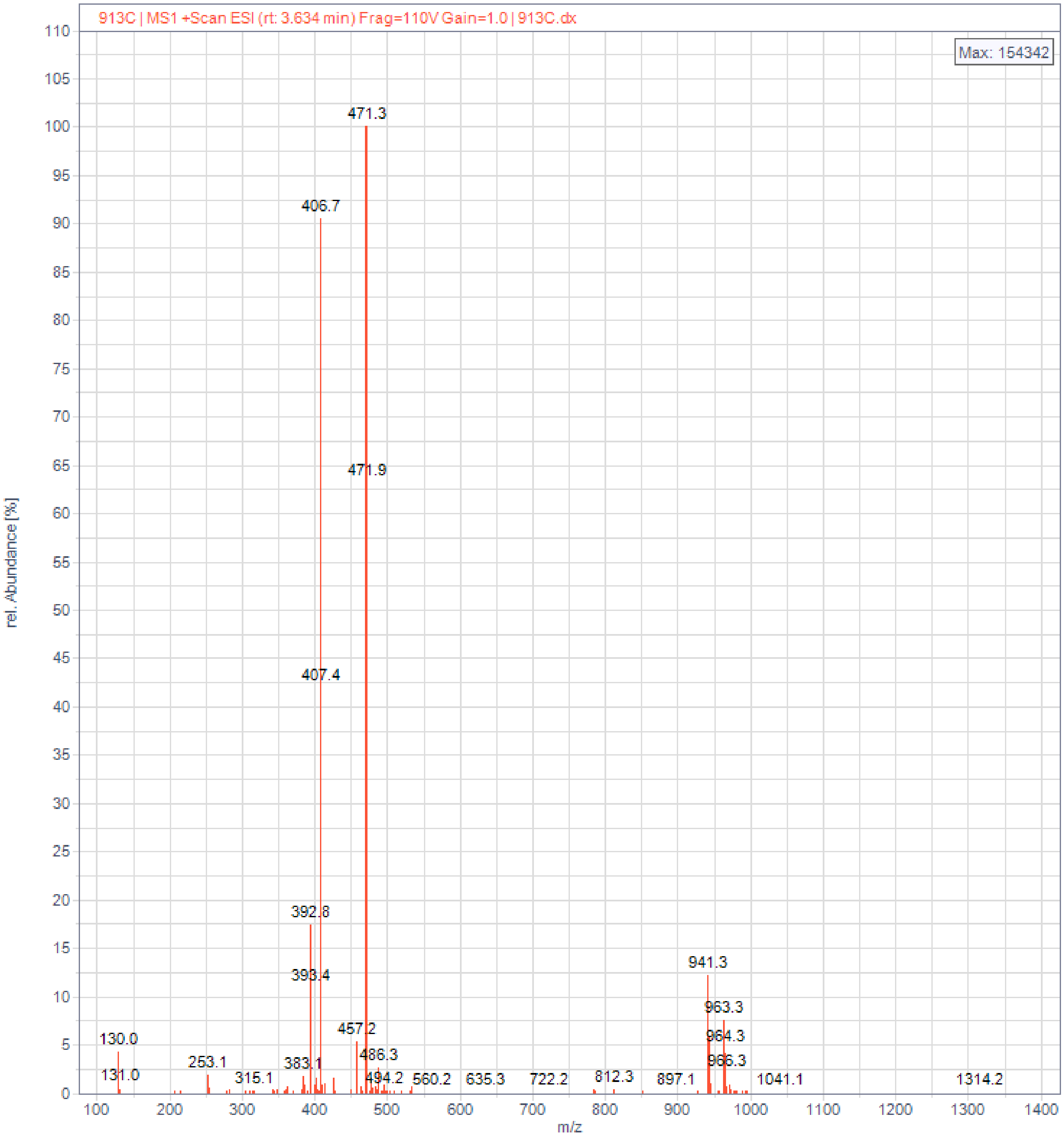

913C | MS1 +Scan ESI (rt: 3.634 min) Frag=110V Gain=1.0 | 913C.dx
Max: 154342
rel. Abundance [%]
m/z
471.3
406.7
471.9
407.4
392.8
393.4
941.3
963.3
964.3
966.3
457.2
486.3
494.2
130.0
131.0
253.1
315.1
383.1
560.2
635.3
722.2
812.3
897.1
1041.1
1314.2

**1024F**

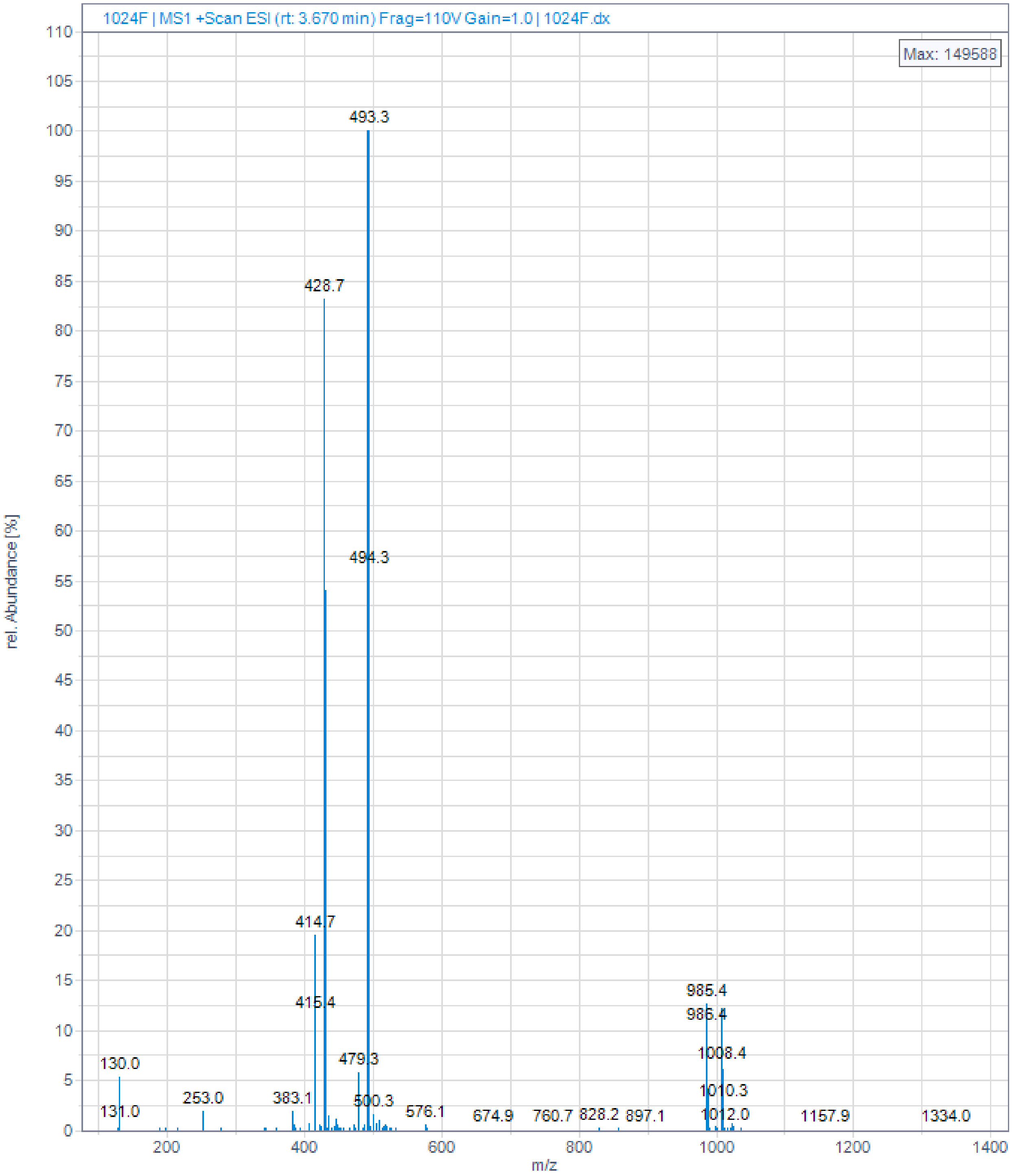

1024F | MS1 +Scan ESI (rt: 3.670 min) Frag=110V Gain=1.0 | 1024F.dx
Max: 149588
rel. Abundance [%]
m/z
493.3
428.7
494.3
414.7
415.4
985.4
986.4
1008.4
1010.3
1012.0
130.0
131.0
253.0
383.1
479.3
500.3
576.1
674.9
760.7
828.2
897.1
1157.9
1334.0

**105D**

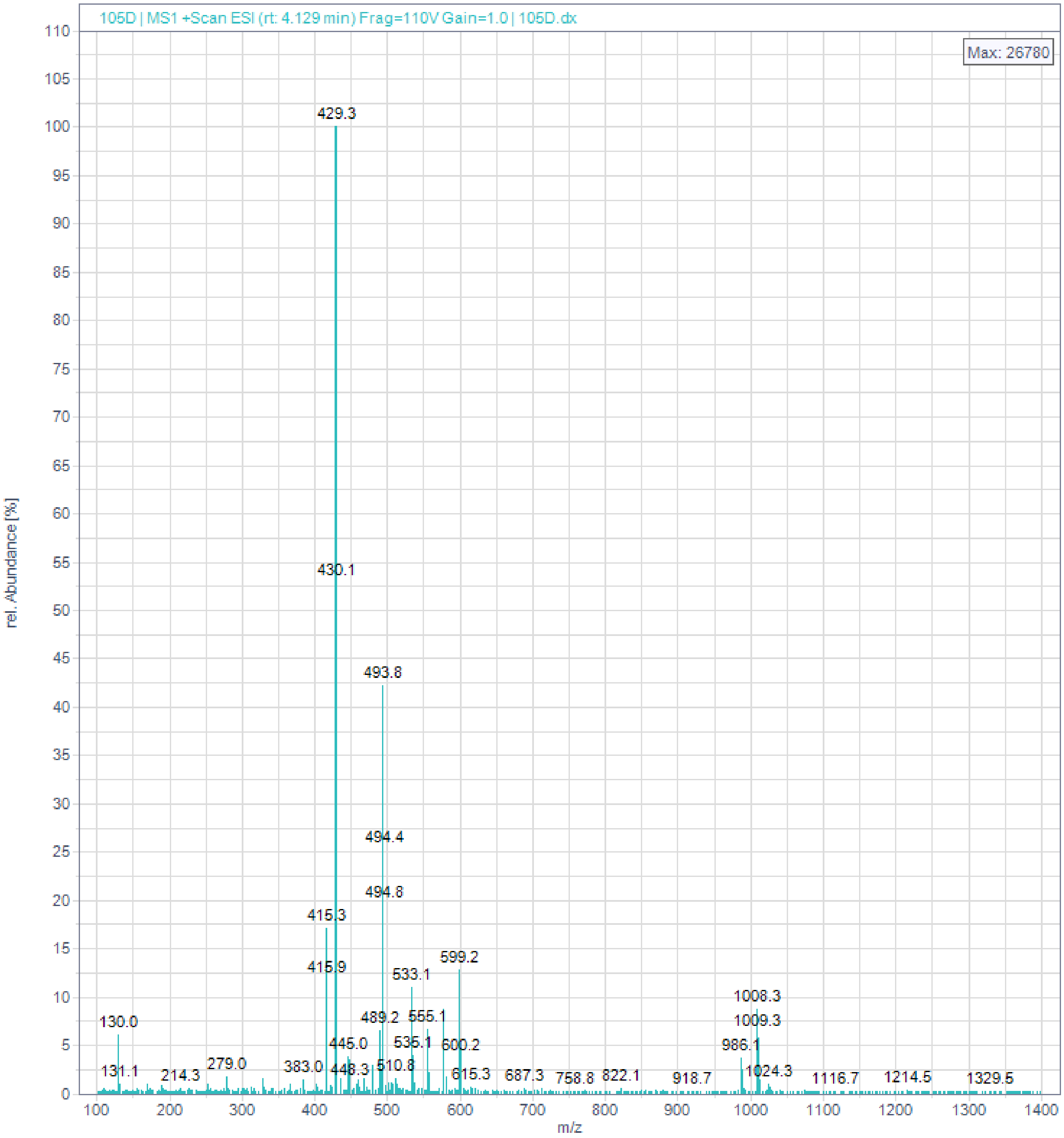

105D | MS1 +Scan ESI (rt: 4.129 min) Frag=110V Gain=1.0 | 105D.dx
Max: 26780
rel. Abundance [%]
m/z
429.3
430.1
493.8
494.4
494.8
415.3
415.9
599.2
533.1
1008.3
1009.3
130.0
489.2
555.1
445.0
535.1
600.2
986.1
279.0
383.0
448.3
510.8
131.1
214.3
615.3
687.3
758.8
822.1
918.7
1024.3
1116.7
1214.5
1329.5

**913D**

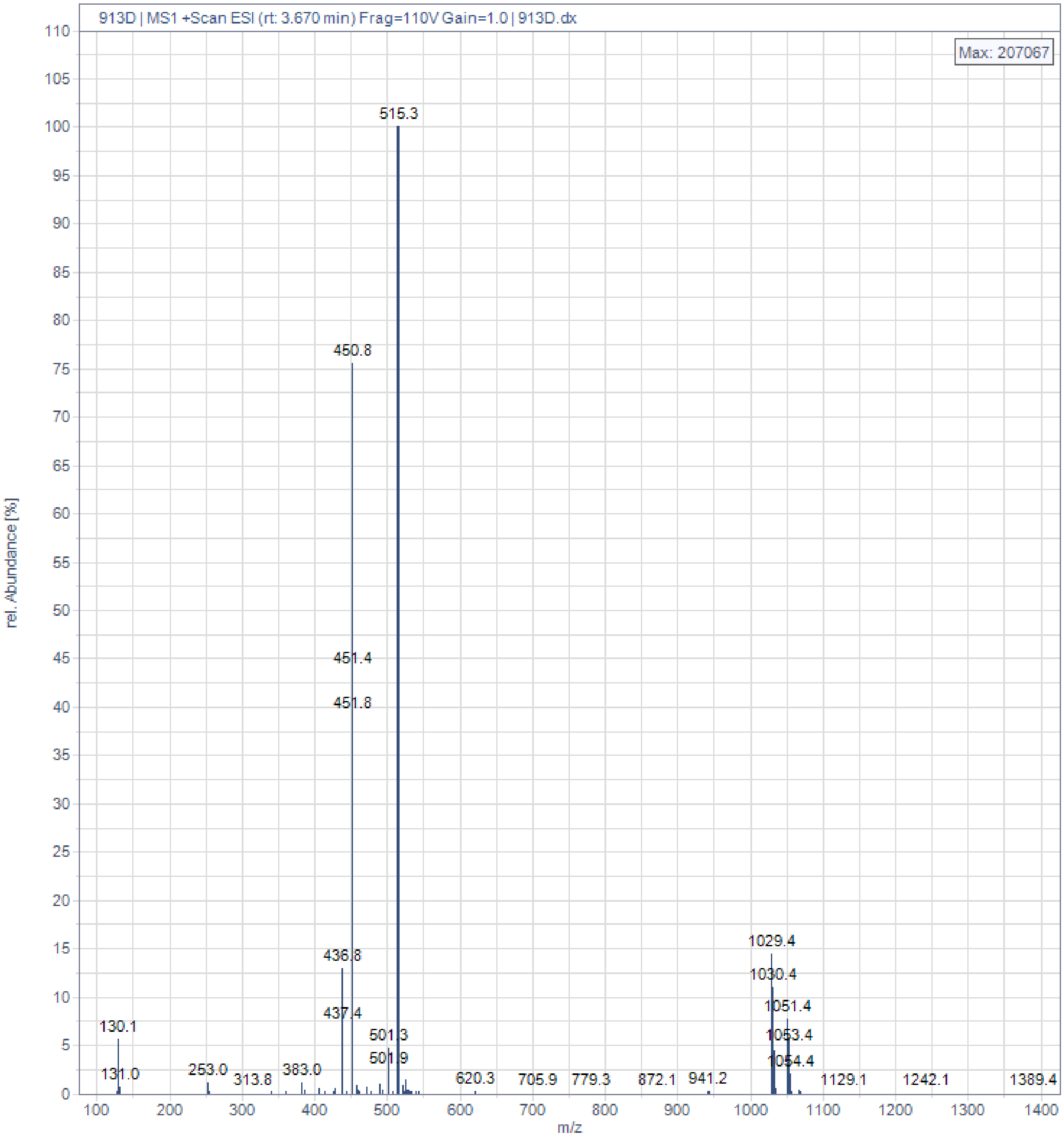

913D | MS1 +Scan ESI (rt: 3.670 min) Frag=110V Gain=1.0 | 913D.dx
Max: 207067
rel. Abundance [%]
m/z
515.3
450.8
451.4
451.8
436.8
437.4
501.3
501.9
130.1
131.0
253.0
313.8
383.0
620.3
705.9
779.3
872.1
941.2
1029.4
1030.4
1051.4
1053.4
1054.4
1129.1
1242.1
1389.4

**105A**

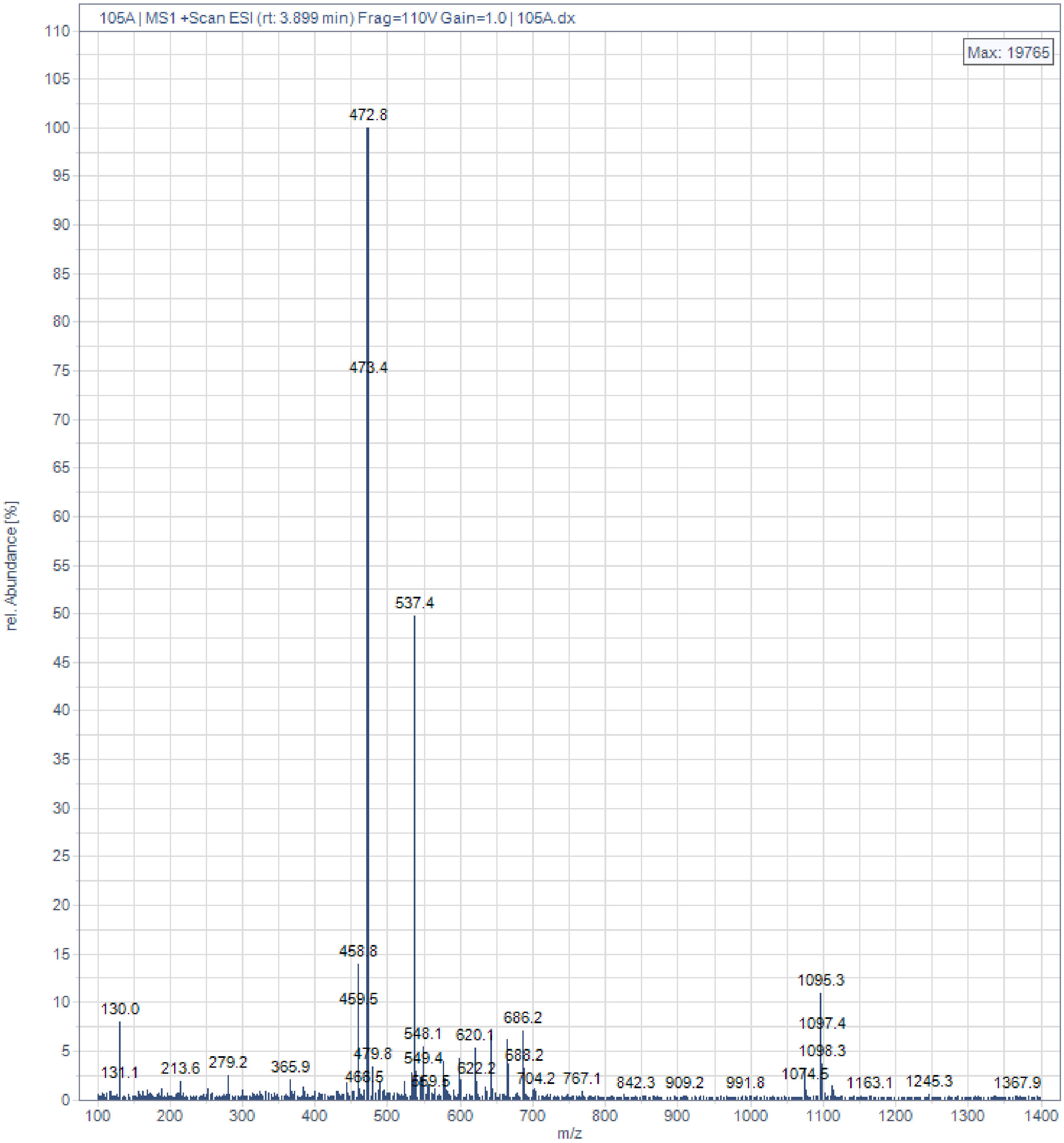
105A | MS1 +Scan ESI (rt: 3.899 min) Frag=110V Gain=1.0 | 105A.dx
Max: 19765
rel. Abundance [%]
m/z
472.8
473.4
537.4
458.8
459.5
1095.3
1097.4
1098.3
1074.5
130.0
131.1
213.6
279.2
365.9
479.8
466.5
548.1
549.4
559.5
620.1
622.2
686.2
688.2
704.2
767.1
842.3
909.2
991.8
1163.1
1245.3
1367.9

**105E**

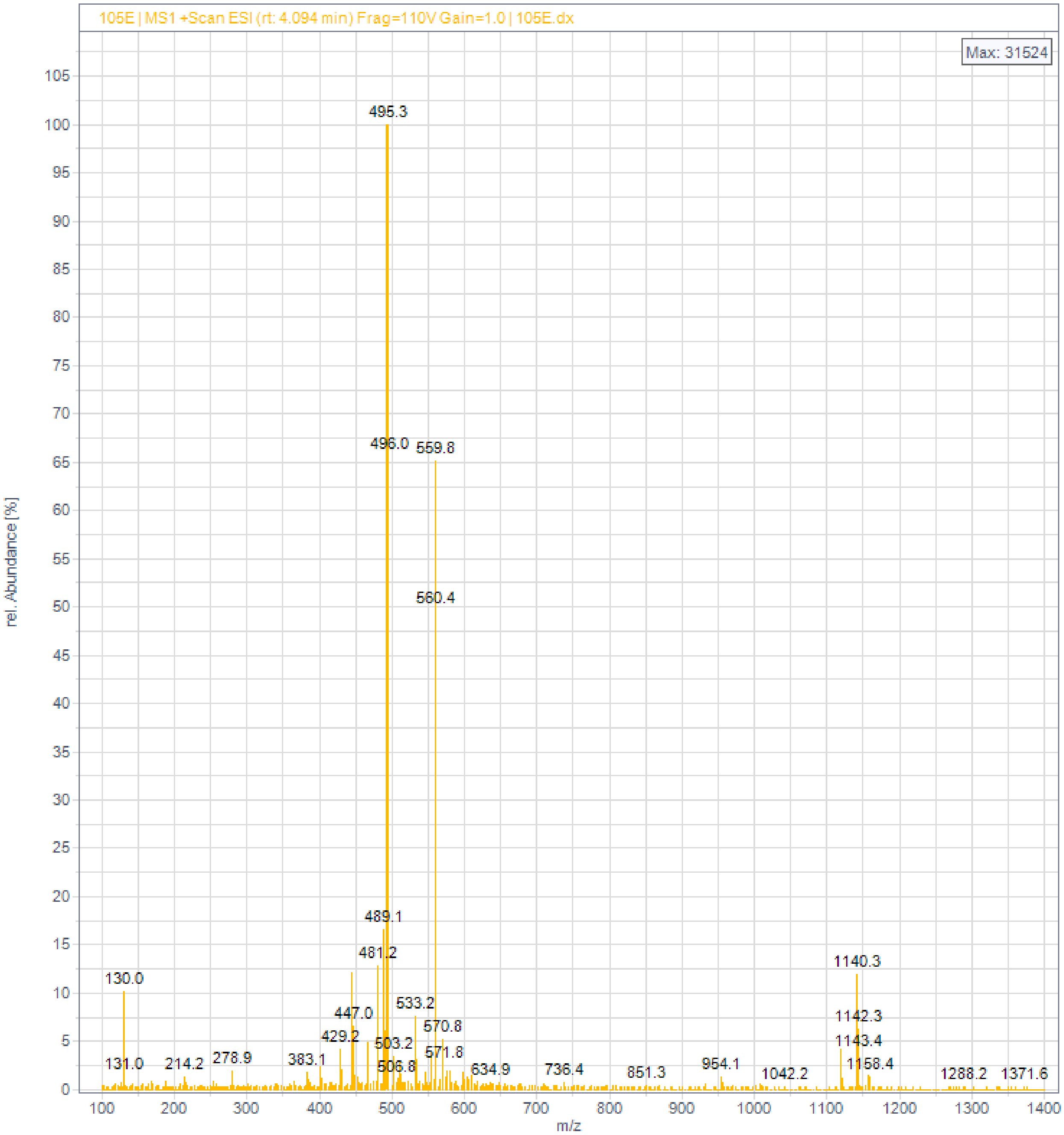
105E | MS1 +Scan ESI (rt: 4.094 min) Frag=110V Gain=1.0 | 105E.dx
Max: 31524
rel. Abundance [%]
m/z
495.3
496.0
559.8
560.4
489.1
481.2
1140.3
130.0
533.2
447.0
1142.3
570.8
429.2
1143.4
503.2
571.8
131.0
214.2
278.9
383.1
506.8
634.9
736.4
851.3
954.1
1042.2
1158.4
1288.2
1371.6

**105C**

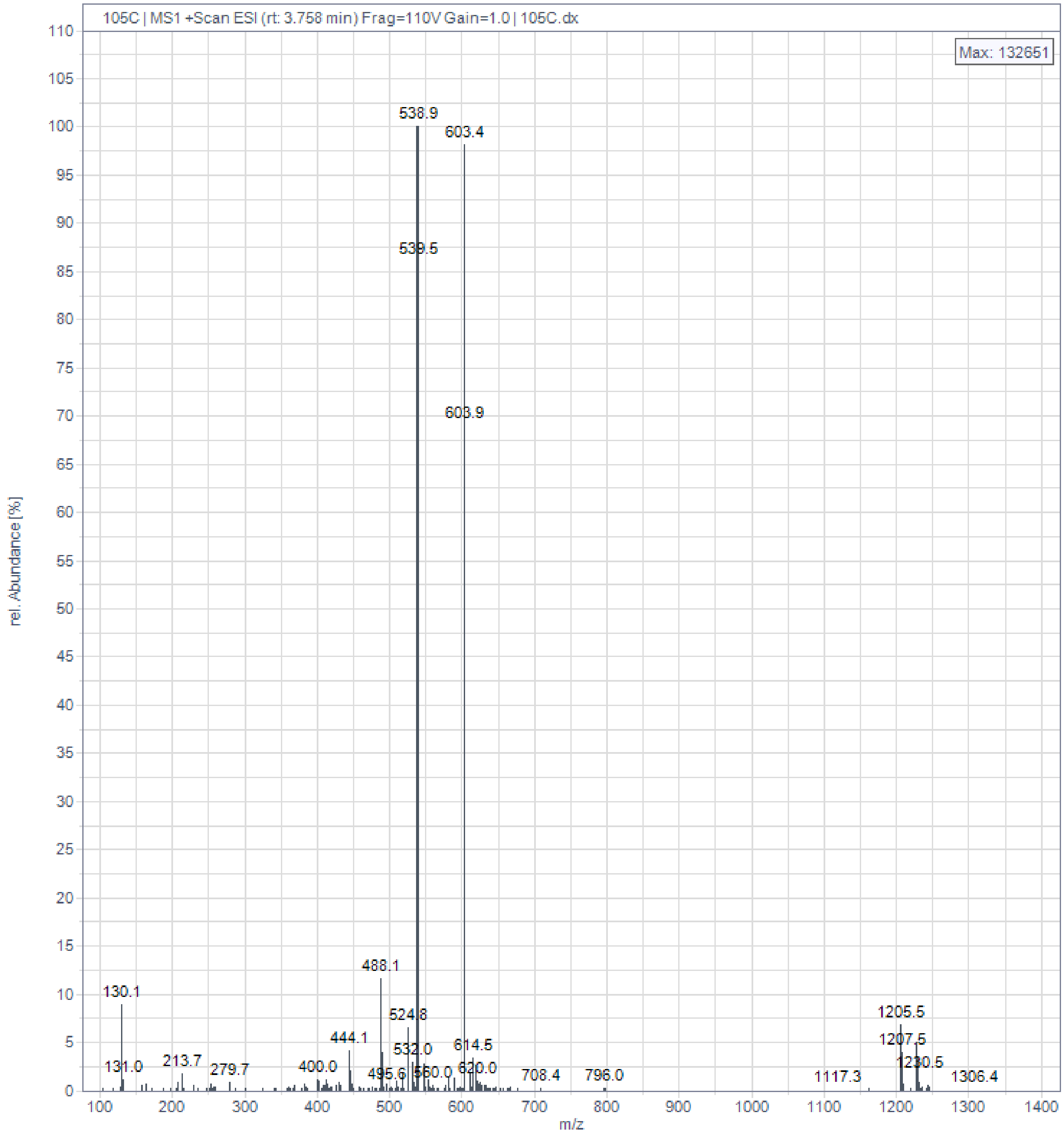

105C | MS1 +Scan ESI (rt: 3.758 min) Frag=110V Gain=1.0 | 105C.dx
Max: 132651
rel. Abundance [%]
m/z
538.9
603.4
539.5
603.9
488.1
130.1
524.8
444.1
532.0
614.5
131.0
213.7
279.7
400.0
495.6
560.0
620.0
708.4
796.0
1117.3
1205.5
1207.5
1230.5
1306.4

**1024C**

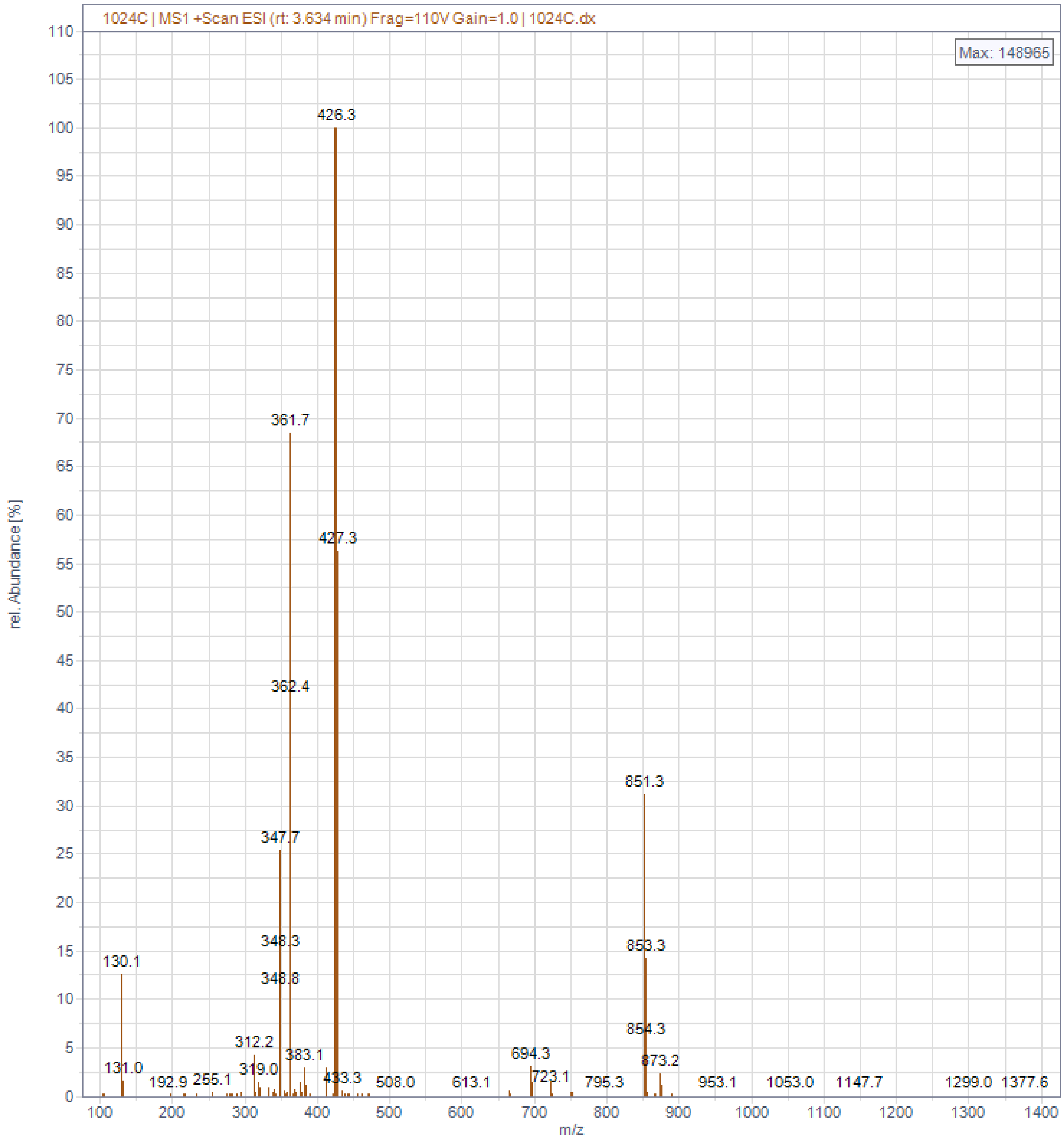
1024C | MS1 +Scan ESI (rt: 3.634 min) Frag=110V Gain=1.0 | 1024C.dx
Max: 148965
rel. Abundance [%]
m/z
426.3
361.7
427.3
362.4
851.3
347.7
348.3
348.8
853.3
130.1
854.3
312.2
383.1
694.3
873.2
131.0
319.0
433.3
723.1
192.9
255.1
508.0
613.1
795.3
953.1
1053.0
1147.7
1299.0
1377.6

**1024B**

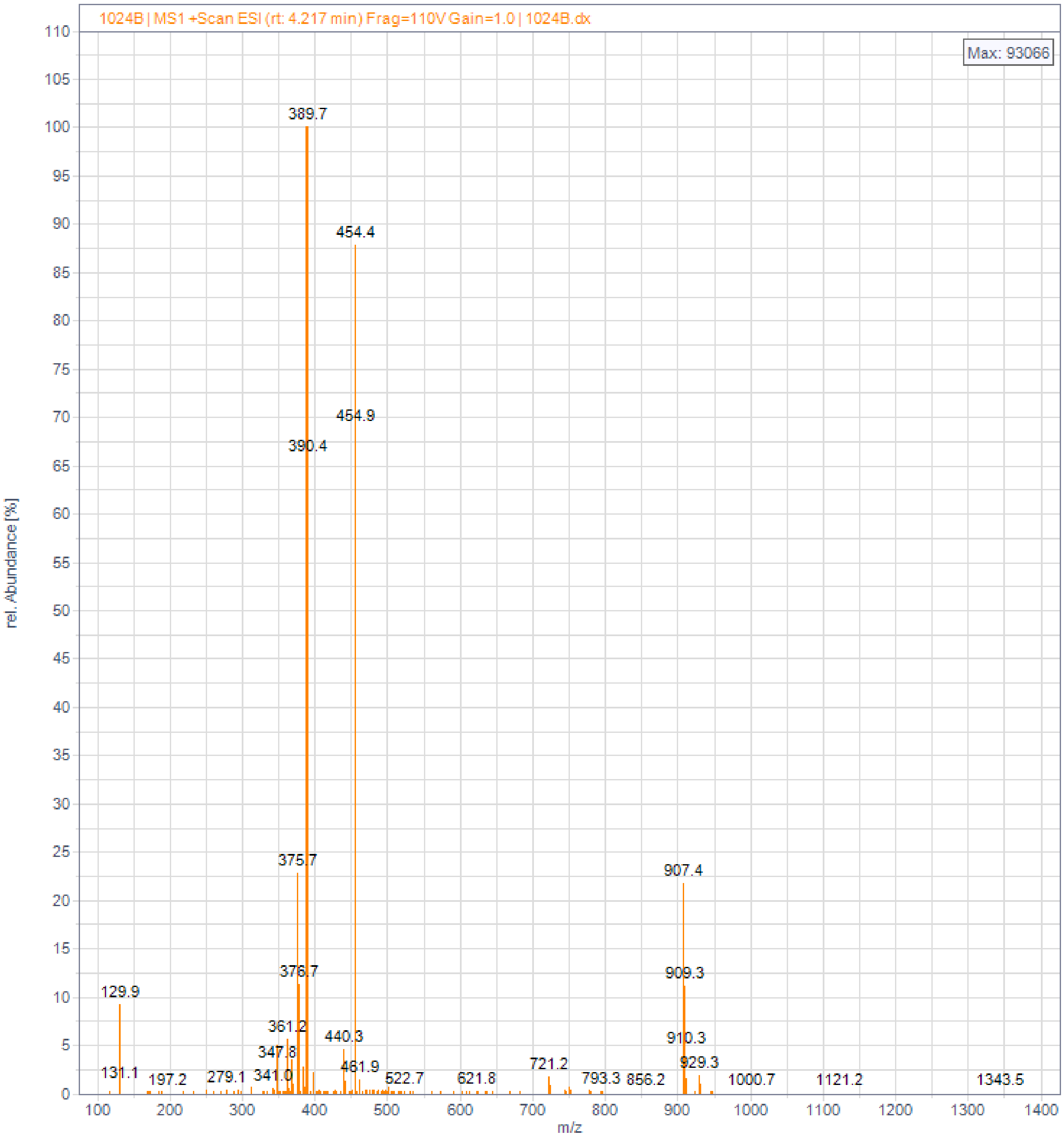

1024B | MS1 +Scan ESI (rt: 4.217 min) Frag=110V Gain=1.0 | 1024B.dx
Max: 93066
rel. Abundance [%]
m/z
389.7
454.4
454.9
390.4
375.7
376.7
361.2
347.8
341.0
440.3
461.9
522.7
621.8
721.2
793.3
856.2
907.4
909.3
910.3
929.3
1000.7
1121.2
1343.5
129.9
131.1
197.2
279.1

**1024E**

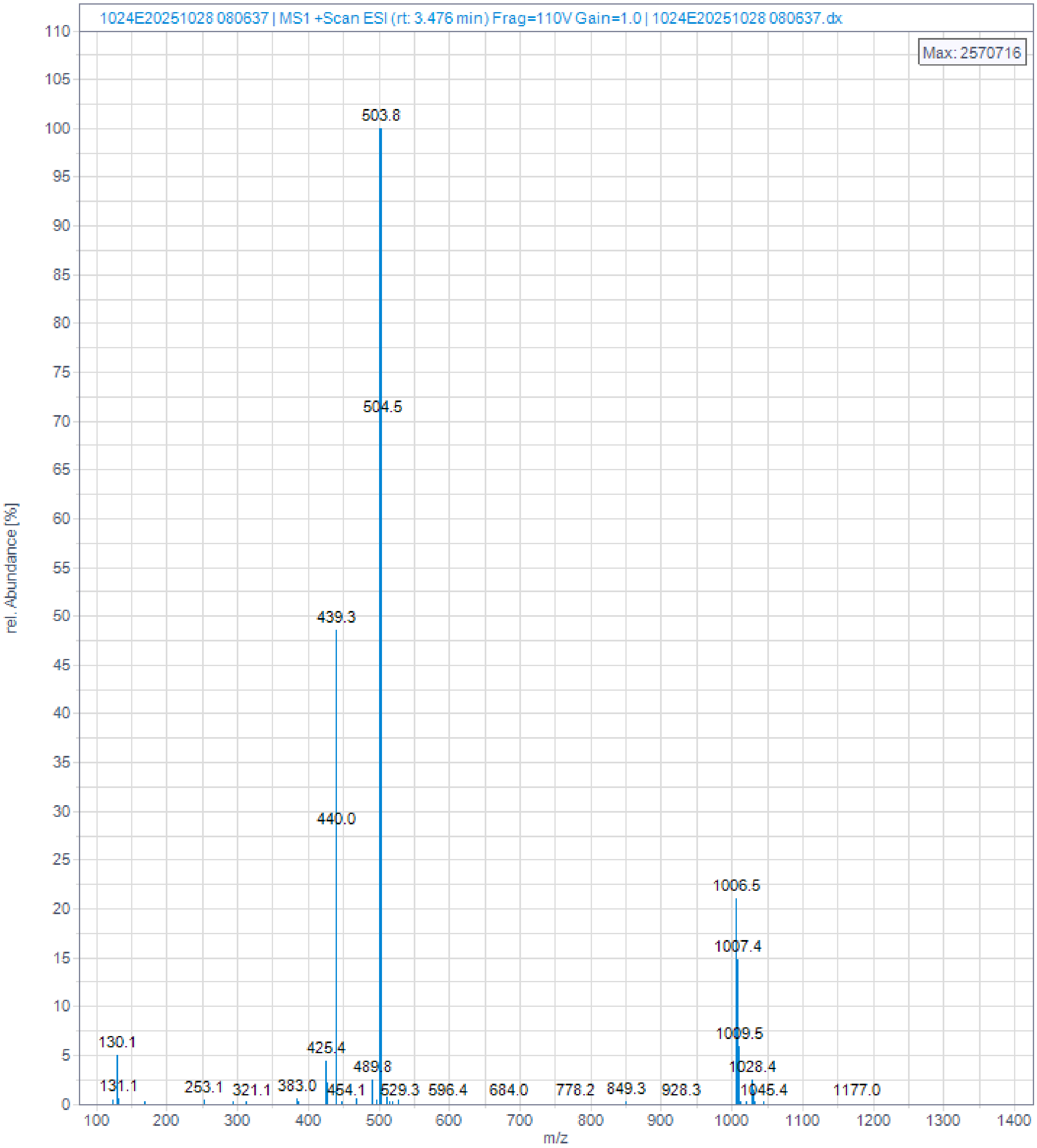

1024E20251028 080637 | MS1 +Scan ESI (rt: 3.476 min) Frag=110V Gain=1.0 | 1024E20251028 080637.dx
Max: 2570716
rel. Abundance [%]
m/z
503.8
504.5
439.3
440.0
1006.5
1007.4
1009.5
1028.4
1045.4
1177.0
130.1
131.1
253.1
321.1
383.0
425.4
454.1
489.8
529.3
596.4
684.0
778.2
849.3
928.3

**1024D**

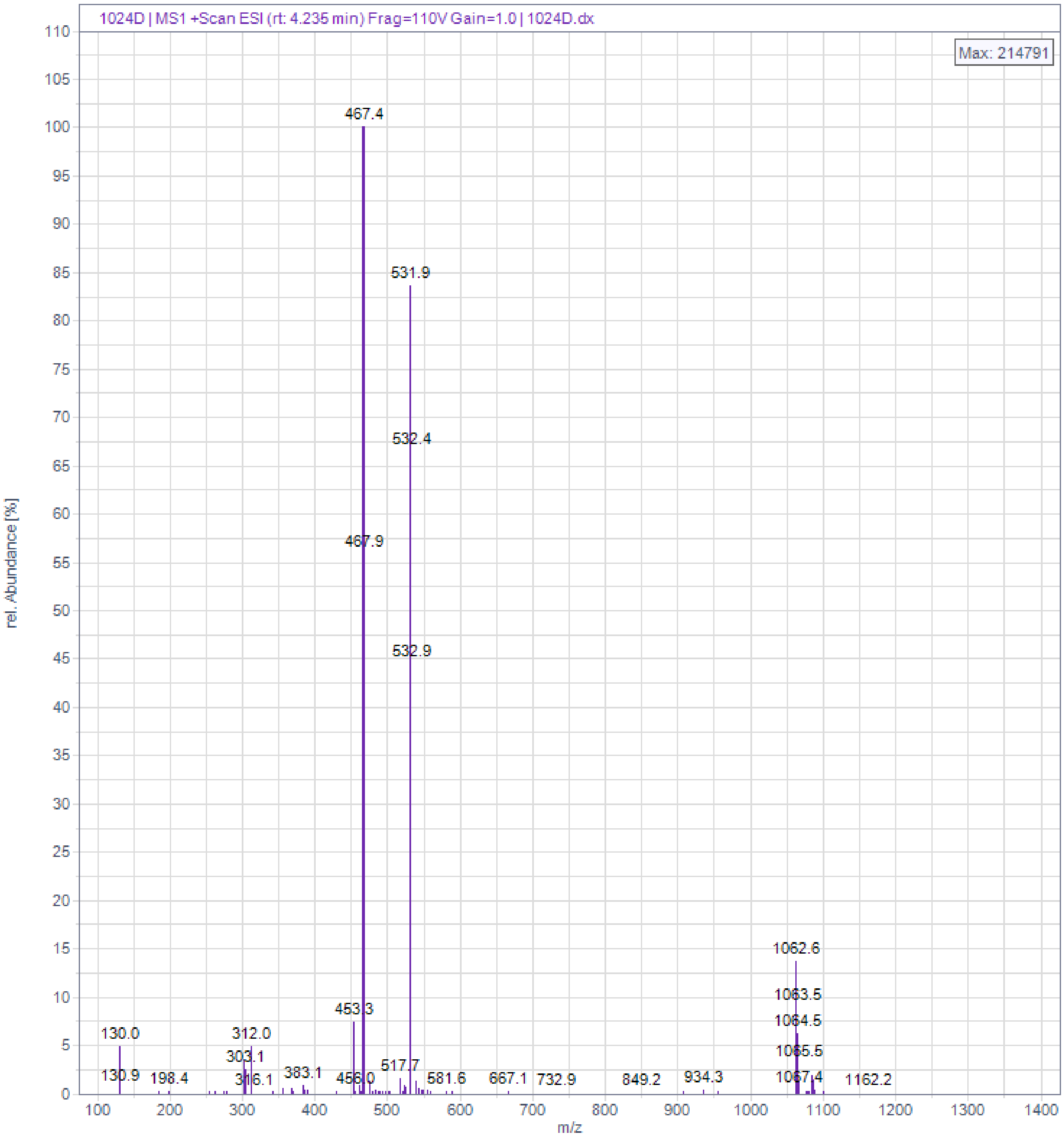

1024D | MS1 +Scan ESI (rt: 4.235 min) Frag=110V Gain=1.0 | 1024D.dx
Max: 214791
rel. Abundance [%]
m/z
467.4
531.9
532.4
467.9
532.9
1062.6
1063.5
1064.5
1065.5
1067.4
1162.2
453.3
130.0
312.0
303.1
130.9
198.4
316.1
383.1
456.0
517.7
581.6
667.1
732.9
849.2
934.3
